# Validation of Tau Antibodies for Use in Western Blotting and Immunohistochemistry

**DOI:** 10.1101/2023.04.13.536711

**Authors:** Michael J. Ellis, Christiana Lekka, Hanna Tulmin, Darragh P. O’Brien, Shalinee Dhayal, Marie-Louise Zeissler, Jakob G. Knudsen, Benedikt M. Kessler, Noel G. Morgan, John A. Todd, Sarah J. Richardson, M. Irina Stefana

## Abstract

**Background:** The microtubule-associated protein Tau has attracted diverse and increasing research interest, with Tau being mentioned in the title/abstract of nearly 34,000 PubMed-indexed publications to date. To accelerate studies into Tau biology, the characterisation of its multiple proteoforms, including disease-relevant post-translational modifications (PTMs), and its role in neurodegeneration, a multitude of Tau-targeting antibodies have been developed, with hundreds of distinct antibody clones currently available for purchase. Nonetheless, concerns over antibody specificity and limited understanding of the performance of many of these reagents has hindered research.

**Methods:** We have employed a range of techniques in combination with samples of murine and human origin to characterise the performance and specificity of 53 commercially-available Tau antibodies by Western blot, and a subset of these, 35 antibodies, in immunohistochemistry.

**Results:** Continued expression of residual protein was found in presumptive Tau “knockout” human cells and further confirmed through mass-spectrometry proteomics, providing evidence of Tau isoforms generated by exon skipping. Importantly, many total and isoform-specific antibodies failed to detect this residual Tau, as well as Tau expressed at low, endogenous levels, thus highlighting the importance of antibody choice. Our data further reveal that the binding of several “total” Tau antibodies, which are assumed to detect Tau independently of post-translational modifications, was partially inhibited by phosphorylation. Many antibodies also displayed non-specific cross-reactivity, with some total and phospho-Tau antibodies cross-reacting with MAP2 isoforms, while the “oligomer-specific” T22 antibody detected monomeric Tau on Western blot. Regardless of their specificity, with one exception, the phospho-Tau antibodies tested were found to not detect the unphosphorylated protein.

**Conclusions:** We identify Tau antibodies across all categories (total, PTM-dependent and isoform-specific) that can be employed in Western blot and/or immunohistochemistry applications to reliably detect even low levels of Tau expression with high specificity. This is of particular importance for studying Tau in non-neuronal cells and peripheral tissues, as well as for the confident validation of knockout cells and/or animal models. This work represents an extensive resource that serves as a point of reference for future studies. Our findings may also aid in the re-interpretation of existing data and improve reproducibility of Tau research.

## Background

Discovered over four and a half decades ago [1], the microtubule-associated protein Tau has attracted ample research interest owing to its association with a wide range of neurodegenerative diseases, particularly tauopathies, a family of dementias marked by abnormal accumulation of protein aggregates containing hyperphosphorylated Tau [2,3]. The Tau protein comprises an amino (N)-terminal region, proline-rich mid-region, microtubule-binding repeats (MTBR) domain, and a carboxy (C)-terminal region (Fig. 1B). *MAPT*, the gene encoding Tau, is made up of 16 exons of which the first (exon -1) is not transcribed, six are alternatively spliced (exons 2, 3, 4A, 6, 8 and 10), while the last (exon 14) together with part of exon 13, encode the 3’ UTR (Fig. 1A). When exons 4A, 6 and 8 are excluded, the alternative splicing of exons 2, 3 and 10 gives rise to the six common Tau splice isoforms that are expressed in the adult human brain, distinguished by the presence of zero, one, or two N-terminal domains (0N, 1N and 2N Tau isoforms) and either three or four MTBRs (3R and 4R Tau isoforms) (Fig. 1B). The resulting protein isoforms have predicted molecular weights (MW) ranging from 36.7 to 45.9 kDa, but the unmodified recombinant proteins migrate on sodium dodecyl sulphate-polyacrylamide gel electrophoresis (SDS-PAGE) as a series of closely spaced bands with apparent MWs ranging from 58 to 66 kDa, thus showing abnormal retardation in electrophoretic mobility [4]. Truncation or, less commonly, skipping of constitutively-included exons can generate Tau species with a MW below that of the shortest splice isoform [5–9], while inclusion of exons 4A and/or 6 gives rise to mid-MW and high-MW Tau isoforms, known as Big Tau or PNS-Tau [10–14]. Inclusion of exon 8 has been observed in bovine Tau [15], but has not been reported in human. The study of Tau is further complicated by the fact that each isoform is subjected to a large number of potential post-translational modifications (PTMs), including phosphorylation, acetylation, glycosylation, methylation, ubiquitination and truncation [16,17]. Among these, phosphorylation is one the earliest recognised and most conspicuous Tau PTMs, with the 2N4R brain Tau isoform containing 85 residues that can accept a phosphate group, over 45 of which have been found to be phosphorylated *in vivo* or *in vitro*. As a consequence, unless subjected to phosphatase treatment to remove phosphate groups prior to analysis, Tau present in denatured protein extracts is commonly detected on SDS-PAGE as an electrophoretically-heterogeneous “smear” or multiple, overlapping diffuse bands.

**Figure 1.**
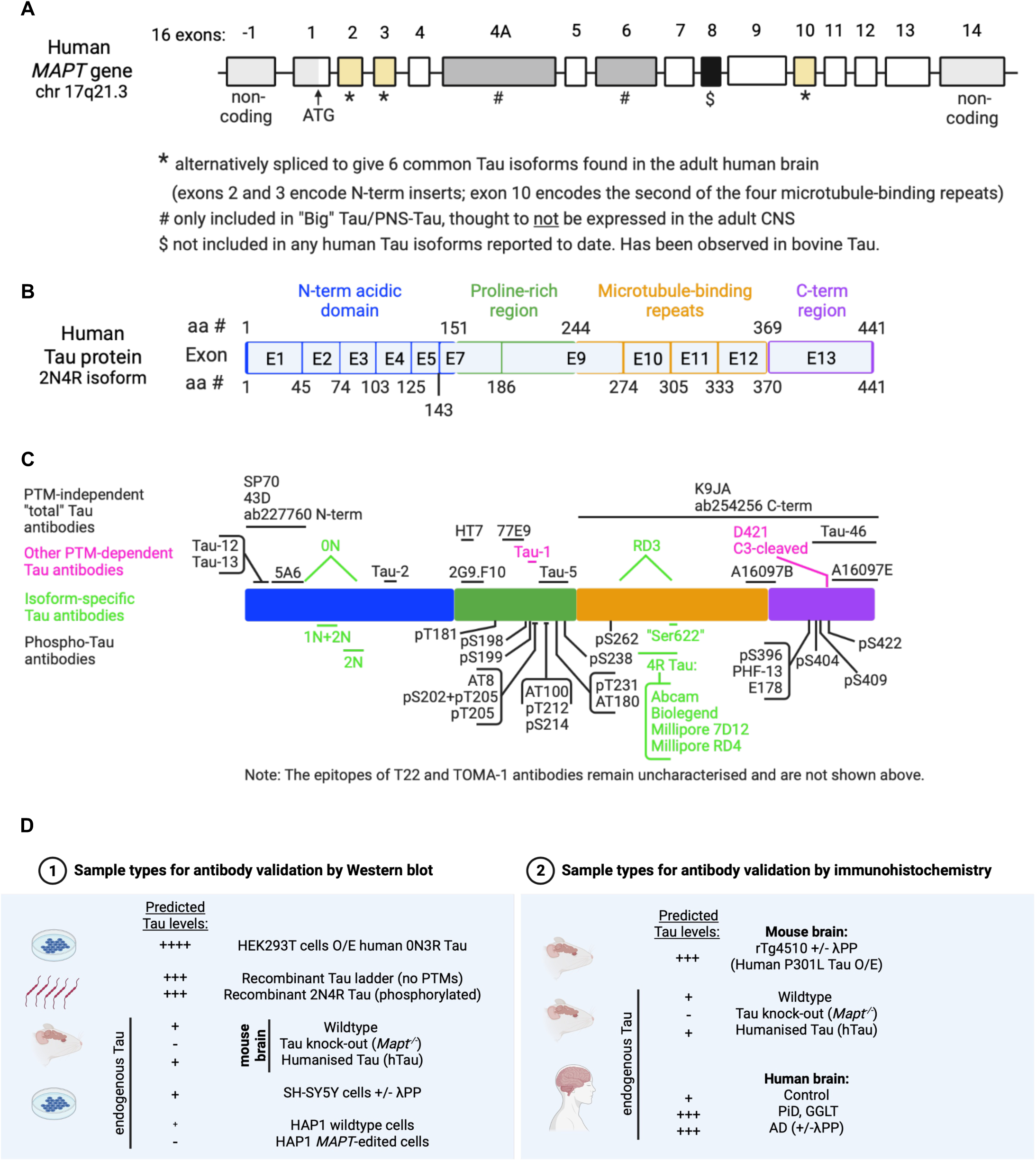
Overview of the human *MAPT* gene, Tau protein, antibody epitopes and antibody validation experimental strategies. **A:** Diagram of the human *MAPT* gene structure with currently-described exons depicted as rectangles and introns depicted as connecting lines. Exon numbering shown above. Canonical transcription start site (ATG) located in exon 1 is indicated (black arrow). Non-coding exonic regions are shown in light grey. Constitutively-included exons (1, 4, 5, 7, 9, 11, 12 and 13) are shown in white. Alternatively-spliced exons included in common brain Tau isoforms, Big Tau isoforms or not included in any human Tau isoforms described to date are shown in yellow (exon 2,3 and 10), light grey (exons 4A and 6) and black (exon 8), respectively. **B**: Diagram of the human Tau protein (2N4R isoform, longest canonical Tau isoform), showing the four main protein domains: N-terminal acidic region (blue), proline-rich mid region (green), microtubule-binding repeat region (orange) and C-terminal region (purple). Amino acid residues marking the domain boundaries are shown above. Protein regions encoded by different exons are indicated. Amino acid residues marking exon-exon boundaries are shown below. **C**: Schematic depiction of the epitopes targeted by the Tau antibodies included in this study: “total” Tau antibodies (above, black), phospho-Tau antibodies (below, black), other PTM-dependent antibodies (magenta) and isoform-specific antibodies (green). **D**: Overview of the experimental approaches employed in this study to validate Tau antibodies using WB (left) and IHC (right).

Originally thought to be a neuronal-specific protein, Tau expression has now been demonstrated in oligodendrocytes, astrocytes and various retinal cell types in the central nervous system (CNS), and has also been reported in many other tissues, including the peripheral nervous system (PNS), submandibular gland, tonsil, liver, colon, heart, skeletal muscle, breast, kidney, skin, pancreas, adrenal gland, reproductive organs and salivary glands [18–34]. Given its pathological relevance, studies to date have focussed largely on the role of Tau hyperphosphorylation and aggregation in driving neurodegeneration, mostly employing overexpression models or patient brain samples where Tau is expressed at high levels and is often found in aggregated states. In contrast, its primary physiological functions have received relatively little attention. Outside the brain, little is known about Tau expression or its roles. Understanding the physiological functions of Tau and how these are affected in the context of tauopathies and other diseases requires that endogenous, unaggregated Tau is detected. This, in turn, depends on the consistent availability of high-quality, validated reagents and protocols that can be employed reliably. Such research has been hindered by uncertainty over antibody specificity and the challenges of detecting the endogenous protein under non-disease conditions, a situation that is exacerbated in non-brain tissues where it is expressed at low levels.

Antibodies are among the most widely-employed tools in the study of proteins and, as such, an ever-increasing multiplicity of anti-Tau antibodies (referred to as “Tau antibodies” hereinafter) have been developed. Records from antibody databases demonstrate the breadth of Tau antibodies available commercially from different suppliers (**Table 1**), representing hundreds of distinct antibody clones with many others developed in-house by individual research groups [35–42]. Many of these have been widely employed in research studies to date, with the top 20 Tau antibody products on Citeab, representing 15 different antibody clones, having amassed altogether a total of 4,803 citations [43]. This overwhelming variety of antibody resources speaks to the widespread interest in the protein, but also highlights the difficulties of developing reliable, high-quality antibodies.

**Table 1.** Summary of existing Tau antibodies compiled from publicly-available antibody databases

| Database name | No. of Tau antibodies | Notes |
| --- | --- | --- |
| Benchsci [44] | 10,830 | Search term: "MAPT [human]" |
| Antibodypedia [45] | 6,454 | 53 suppliers |
| Alzforum [46] | 572 | Includes records of antibodies developed by individual research laboratories but which are not currently available commercially. |
| Labome [47] | 460 | 29 suppliers.<br>135 reagents have been cited in published studies.<br>87 reagents included on the Labome "validated antibodies" list |
| Citeab [43] | 3,502 | Search term: "MAPT" + filtered for "Protein = Tau (Human)" |

Quality issues with research antibodies contribute to the widely acknowledged “reproducibility crisis” in scientific research [48–52]. Despite an urgent need for validated reagents, and sustained efforts from both the scientific community and publishers [52–64], no universally-agreed standards exist for antibody validation and most antibodies used in research studies have not been adequately validated. This problem is compounded by the fact that antibody performance is both application-dependent and context-specific [58,65–67]. Poorly characterised antibodies can cause significant loss of time and financial resources, and lead to artefactual findings or misinterpretation of research data. Previous studies that have addressed Tau antibody validation have focused on either: (i) the reliability of a small number of phosphorylation-dependent antibodies to detect *in vitro*-phosphorylated recombinant Tau [68]; (ii) antibody performance in mouse brain samples by Western blotting (WB; [69]); (iii) characterising PTM-dependent antibodies by WB and immunocytochemistry [70]; or (iv) testing a small set of phospho-Tau antibodies by fluorescence-activated cell sorting [71]. Here, we use a combination of recombinant proteins, protein overexpression in human cells, human and murine cell/tissue samples that express endogenous Tau, and brain tissue from *Mapt^-/-^* mice as well as human cells that have undergone genomic editing to disrupt the *MAPT* gene, to characterise a panel of 53 commercially-available Tau antibodies for use in WB. A subset of these (35 antibodies) were further verified for use in immunohistochemistry (IHC) applications. WB and IHC were chosen owing to their status as the two most common uses of antibodies overall [58] and the two most common applications for Tau antibodies, according to antibody reagent portals BenchSci[44], Labome [47]and Antibodypedia [45]. We identify high-quality antibodies that demonstrate specificity towards the Tau protein and can be employed reliably to investigate the expression and roles of endogenous Tau proteoforms. To enable studies of clinical relevance, we focus on identifying antibodies that react with human Tau. By including only commercially-available antibodies, we have produced a resource defining readily-available reagents that has relevance to the entire research community. Identifying antibodies that can be used to reliably detect Tau in samples and tissue types where it is expressed at low levels will inevitably broaden Tau research, enabling a deeper understanding of its physiological roles in the CNS as well as its functions outside of the brain.

## Methods

### Antibodies

The complete list of antibodies used in this study is displayed in Supp. Table S1.

### Cell culture

Human embryonic kidney (HEK) 293T cells were maintained in Dulbecco’s Modified Eagle’s Medium (DMEM, Sigma, cat. no. D6429) supplemented with 10% heat-inactivated foetal bovine serum (hiFBS, Sigma, cat. no. F9665) at 37°C under a 5% CO_2_ humidified atmosphere. No antibiotics were used. Cells were split upon reaching 90-95% confluence (approximately every 2-3 days). HEK293T Lenti-X cells (Takara Bio, cat. no. 632180) were maintained in DMEM supplemented with 10% hiFBS, 1x GlutaMAX (Gibco, cat. no. 13462629) and 1 mM sodium pyruvate (Gibco, cat. no. 11360070), at 37°C under a 5% CO_2_ humidified atmosphere. No antibiotics were used. Cells were split upon reaching 90-95% confluence (approximately every 2-3 days). SH-SY5Y cells (European Collection of Authenticated Cell Cultures via Sigma-Merck, cat. no. 94030304) were maintained in DMEM:F12 (1:1) medium (Gibco, cat. no. 11330057) supplemented with 10% hiFBS (Sigma, cat. no. F9665) at 37°C under a 5% CO_2_ humidified atmosphere. No antibiotics were used. Cells were split upon reaching ∼90% confluency (approx. every 5 days) at a ratio of 1:5. Cells were collected for experiments when 85-90% confluent. Parental control and *MAPT*-edited HAP1 cells were purchased from Horizon Discovery (parental control cells cat. no. C631; 2 bp deletion cells cat. no. HZGHC003277c009; 14 bp deletion cells cat. no. HZGHC003277c003). The two *MAPT*-edited cell lines were generated by CRISPR/Cas9 editing using the same guide RNA (sequence: GACCAGCAGCTTCGTCTTCC) and carry either a 2 bp or a 14 bp deletion in *MAPT* exon 4 (exon numbering as shown in Fig. 1), in both cases leading to out-of-frame shifts and the introduction of early stop codons. Following the supplier’s instructions, cells were maintained in Iscove’s Modified Dulbecco’s Medium (Sigma, cat. no. I3390) supplemented with 10% hiFBS and 1% Pen/Strep at 37°C under a 5% CO_2_ humidified atmosphere. Cells were split upon reaching 70-75% confluency (approximately every 2-3 days). Cells were collected for experiments when ∼70% confluent.

### Tau overexpression in HEK293T cells

HEK293T cells were transfected with either Tau-encoding or control constructs using Fugene 6 (Promega, cat. no. E2691) according to the manufacturer’s instructions. Transfection rates of >90% and high expression of the transgenic construct were confirmed under the fluorescence microscope 48 hours post-transfection. Cells were lysed (as described below) for protein extraction 72 hours post-transfection. Plasmids used for tdTomato-Tau overexpression and the tdTomato-C1 control plasmid were a gift from Michael Davidson: tdTomato-MAPTau-C-10 (Addgene plasmid #58112), tdTomato-MAPTau-N-10 (Addgene plasmid #58113) and tdTomato-C1 (Addgene plasmid #54653). Control tdTomato-N1 plasmid was a gift from Michael Davidson, Nathan Shaner and Roger Tsien (Addgene plasmid #54642) [72].

### Expression of transgenic 2N4R Tau in SH-SY5Y cells

The cDNA sequence of the 441 amino acids-long 2N4R Tau isoform was cloned into the pSMPUW-IRES-Bsd vector backbone (Cell Biolabs, cat. no. VPK-219). The empty vector was used as a control. Lentiviral particles were generated by co-transfecting HEK293T Lenti-X cells with either the empty or the 2N4R-encoding pSMPUW-IRES-Bsd transfer vector and with packaging vectors – pRSV-Rev, pCMV-VSV-G and pCgpV (Cell Biolabs, ViraSafe Lentiviral Packaging System, cat. no. VPK-206) –, using the PEIPro DNA transfection reagent (Polyplus, cat. no. 115-010) following manufacturer’s instructions. The molar ratio of transfer vector to packaging vectors was 3:1:1:1. Media containing lentiviral particles was collected 72 hours post-transfection and the lentiviral particles were precipitated with 3 volumes of Lenti-X Concentrator (Takara, cat. no. 631232) at 4°C overnight, following manufacturer’s instructions. The lentiviral stock was aliquoted in single-use aliquots and stored at -80°C. To generate SH-SY5Y cells that stably express transgenic 2N4R Tau, lentiviral stock (either empty or Tau-encoding lentiviral particles) was added to cells in suspension, prior to seeding cells into flasks. Following 20 hours of transduction, the virus was removed, cells were split into fresh flasks and allowed to recover for 24 hours. Transduced cells were then selected with blasticidin (7 µg/mL) for 6 days, until all untransduced, positive-control cells had died. Fresh antibiotic was provided every 2 days during the selection protocol.

### Recombinant proteins

An equimolar mixture of recombinant versions of the six common Tau splice isoforms (0N3R, 0N4R, 1N3R, 1N4R, 2N3R and 2N4R), referred to as the Tau ladder (either Sigma cat. no. T7951, or Signal Chem cat. no. T08-07N-250), was used as a control for antibody validation by WB. Note that the Signal Chem Tau ladder contains a seventh Tau isoform of a MW ∼10 kDa lower than 0N3R, the smallest of the six common Tau isoforms. Recombinant 2N4R Tau that has been phosphorylated, *in vivo* and *in vitro*, with specific kinases was also used to validate phosphorylation-dependent antibodies and to test the impact of phosphorylation on binding of total and isoform-specific Tau antibodies: GSK3β-phosphorylated Tau (Signal Chem, cat. no. T08-50FN-20), DYRK1A-phosphorylated Tau (Signal Chem, cat. no. T08-50RN-20) and CAMK2A-phosphorylated Tau (Signal Chem, cat. no. T08-50CN-20). Recombinant MAP2c protein (Abcam, #ab114686), carrying a ∼26 kDa Glutathione-S-Transferase (GST) tag at the N-terminus, was used to test for cross-reactivity of Tau antibodies with MAP2. Recombinant proteins were resuspended, if applicable, according to manufacturer’s instructions, aliquoted and stored at -80°C until use.

### Animals

In this study, we used wildtype, *MAPT* knockout (*Mapt*^-/-^; [73]) and humanised Tau (hTau; [74]) mice, all of which were maintained on a C57Bl/6J background. hTau mice were generated using P1-derived artificial chromosome (PAC) technology to express the entire 143 kb wildtype human *MAPT* locus (H1 haplotype) in *Mapt*^-/-^ mice. The 143 kb transgene comprises of 9 kb of promoter sequence and 134 kb genomic DNA, including the two alternative polyadenylation sequences at the 3’ end of the human *MAPT* locus [75]. This allows the physiological expression of all human Tau isoforms under the control of the endogenous human *MAPT* promoter. Animals were sacrificed at five months of age by cervical dislocation, brains were dissected quickly, and half of the brain was snap-frozen in liquid nitrogen and stored at −80 °C until lysis. The other half of the brain was fixed in formalin and embedded in paraffin blocks. Sagittal sections of 4 µM thickness were used for immunohistochemistry. Tissues from female mice were used in this study. rTg4510 mice [76] used in the study were acquired from Eli Lily through collaboration with Dr Jonathan Brown (Exeter University, UK). Dissected brains were formalin-fixed and embedded in paraffin blocks. 4 µM coronal sections were cut from the forebrain of 9-month-old rTg4510 male mice using a Leica RM2235 microtome. All housing and experimental procedures were carried out in compliance with the local ethical review panel of the Universities of Oxford and Exeter, respectively, under UK Home Office project licenses held in accordance with the Animals (Scientific Procedures) Act 1986. Animals were housed under 12-hour light/dark cycle with *ad libitum* access to food and water.

### Human brain tissue sections

Anonymized human post-mortem brain tissue (4 µM FFPE brain sections) was obtained from the Oxford Brain Bank. Tissue was obtained from donors with: no known brain pathology (donor ID NP91/2018), diagnosed Alzheimer’s disease (donor ID NP25/2018), a diagnosed 4R tauopathy (globular glial tauopathy, donor ID NP61/2018), and a diagnosed 3R tauopathy (Pick’s disease, donor ID NP39/2016).

### Preparation of protein samples and Western blotting

For preparation of cell lysates, cells were washed twice with PBS at room temperature (RT) and lysed in 1x (radio immunoprecipitation assay buffer (RIPA buffer, Merck Millipore, cat. no. 20-188) containing 0.1% SDS, 1x cOmplete^™^ EDTA-free Protease Inhibitor Cocktail (Roche, cat. no. 11873580001) and 1x PhosSTOP™ phosphatase inhibitors (Roche, cat. no. 4906837001). Lysis was allowed to proceed on ice for 15 minutes before cells were scraped from the cell culture flask, transferred to low-protein binding microcentrifuge tubes and lysis was allowed to proceed for a further 15 minutes on ice, for a total of 30 minutes. Lysates were then spun down at 20,000 x g for 20 minutes to pellet genomic DNA and cellular debris, and the supernatant was aliquoted into fresh low-protein binding tubes. Lysates were stored at -80°C.

To extract proteins from mouse brains, frozen tissues were ground in a mortar on dry ice and grindates were transferred to low-protein binding microcentrifuge tubes. Grindates were then sonicated on ice in lysis buffer (1x RIPA buffer containing 0.1% SDS, cOmplete^™^ EDTA-free Protease Inhibitor Cocktail and PhosSTOP™ phosphatase inhibitors; 16 µL lysis buffer per mg tissue grindate), samples were spun at 20,000 x g for 20 minutes and the supernatant was aliquoted into fresh low-protein binding tubes. Lysates were stored at -80°C.

Protein concentration was quantified using the Pierce™ BCA Protein Assay Kit (Thermo Fisher Scientific, cat. no. 23227) according to the manufacturer’s instructions. To prepare samples for Western blotting, lysates were thawed on ice, mixed 3:1 with 4x Laemmli buffer (Biorad, cat. no. 1610747) containing β-mercaptoethanol and heated at 95°C for 10 minutes. Samples were adjusted to the same protein concentration by adding 1x Laemmli buffer containing β-mercaptoethanol. Sodium dodecyl sulfate-polyacrylamide gel electrophoresis (SDS-PAGE) was performed by loading: 6 µg per lane for HEK293T cell lysates, 5 ng/isoform per lane for Tau ladder (Sigma, cat. no. T7951), 8 µg per lane for mouse brain extracts, 40 µg per lane for SH-SY5Y and HAP1 cell lysates, and 50 ng per lane for recombinant phosphorylated Tau as well as 50 ng/isoform per lane for the Tau ladder used as a control on the recombinant phospho-Tau blots. Proteins were separated on 4-15% gradient Mini-PROTEAN TGX Stain Free Gels (Bio-Rad, cat. no. 4568084 for 10-well gels, cat. no. 4561086 for 15-well gels) and transferred to Immobilon-FL PVDF membranes (Merck Millipore, cat. no. IPFL00010). Blots were blocked in 5% Amersham™ ECL Prime Blocking Reagent (SLS, cat. no. RPN418) diluted in TBS (blocking buffer, no detergent). Primary antibodies were diluted in blocking buffer containing 0.1% Tween-20 and membranes were incubated with primary antibodies for either 2 hours at RT or overnight at 4°C. For primary antibody details, see Supp. Table S1. Secondary antibodies used were IRDye® 680RD Donkey anti-Rabbit IgG (Licor, cat. no. 925-68073), IRDye® 800CW Donkey anti-Mouse IgG (Licor, cat. no. 925-32212), or IRDye® 800CW Goat anti-Rat IgG (Licor, cat. no. 926-32219) as appropriate, and were diluted 1:10,000 in blocking buffer containing 0.1% Tween-20 and 0.01% SDS. Membranes were incubated with secondary antibodies for 1 hour at RT on an orbital shaker. Blots were washed three times with TBS buffer containing 0.1% Tween-20 after the primary and secondary antibody incubations and once with TBS buffer without detergent at the end. Membranes were imaged dry on an Odyssey CLx scanner (LI-COR Biosciences), and blot images were visualised and quantified in the Image Studio software (LI-COR Biosciences).

### Dephosphorylation of protein extracts with lambda phosphatase

To generate protein extracts for protein dephosphorylation, cells were lysed and proteins were extracted as described above, with the exception that PhosSTOP™ phosphatase inhibitors were omitted from samples to be dephosphorylated, but added to control samples to protect protein phosphorylation. To dephosphorylate proteins, samples were treated with lambda phosphatase (NEB, cat. no. 0753; 10 U enzyme per µg protein) according to manufacturer’s instructions at 30°C for 2 hours. Buffer only was added to control samples, which were incubated on ice for 2 hours to protect protein phosphorylation. At the end of the incubation period, the reaction was terminated by adding 4x Laemmli sample buffer to all samples and heating at 95°C for 10 minutes. Samples were either used immediately or stored at -20°C until further analysis.

### Tau immunoprecipitation from HAP1 cells

To immunoprecipitate (IP) endogenous Tau from parental and *MAPT*-edited HAP1 cells, we used cytoplasmic extracts prepared using the NE-PER kit (Thermo Fisher Scientific, cat. no. 7883, 11 µL CERI reagent/1 x 10^6^ cells) according to the manufacturer’s instructions. cOmplete^™^ EDTA-free Protease Inhibitor Cocktail and PhosSTOP™ phosphatase inhibitors were added to the CERI extraction buffer. Protein extracts were transferred to low-protein binding tubes, stored on ice and used immediately to quantify protein concentration using the Pierce™ BCA Protein Assay Kit and set up the IP reactions. Protein lysates (1 mg per IP reaction) were incubated in low-protein binding microcentrifuge tubes with Tau antibodies (pThr231 or SP70 rabbit monoclonal antibodies; 0.8 µg antibody/mg protein extract), or an equal amount of rabbit monoclonal IgG control antibody (Abcam, cat. no. ab172730), overnight at 4°C with end-over-end mixing, to allow the formation of antibody/protein immune complexes. After addition of protein G-coated magnetic dynabeads (5 μl beads/μg antibody, Thermo Fisher Scientific, cat. no. 13424229), the lysate+antibody+bead mixtures were incubated for 1 hour at 4°C with end-over-end mixing, to allow binding of the antibody/protein complexes to the beads. Beads were then washed three times with 1 mL cold wash buffer (25mM Tris, 150mM NaCl, 1mM EDTA, 1% NP-40, 5% glycerol, pH 7.4), transferred to fresh low-protein binding tubes and proteins were eluted from the beads by incubating in 1.1x Laemmli buffer (containing no reducing agent) at RT for 10 minutes. Eluates were snap-frozen on dry ice and stored at -80°C until further processing.

### Sample processing of Tau IP eluates for proteomic analysis

Eluted proteins were subjected to proteomic processing as follows; all sample volumes were adjusted to 200 μL prior to in-solution digestion. Proteins were reduced in dithiothreitol (5 mM final concentration) and incubated for 45 minutes at RT. Samples were alkylated with iodoacetamide (20 mM final concentration) and incubated for 45 minutes at RT in the dark. Contaminating salts and detergents were removed following methanol/chloroform extraction of proteins. Briefly, to a sample volume of ∼200 µL, 600 µL of methanol and 150 µL chloroform were added and vortexed. For protein precipitation, 450 µL of MilliQ H_2_O was added, vortexed, and spun down (max. speed, tabletop centrifuge) at RT for 2 minutes. The upper aqueous phase was removed without disrupting the precipitate at the interface. Another 450 µL of methanol was added, vortexed, and centrifuged for 2 minutes at RT. The supernatant was discarded and precipitated protein discs were allowed to briefly air dry. Proteins were resuspended in 50 μL 6 M urea buffer with vortexing and brief sonication on ice. Prior to digestion, urea concentrations were reduced to a final concentration of <1M by diluting the reaction mixture with 250 μL MilliQ H_2_O. Trypsin (Sequencing Grade Modified Trypsin; Promega) was added in a 1:50 w/w ratio of enzyme:total protein and incubated overnight at 37 °C. The following day, samples were acidified to 1% TFA to stop the digestion reaction. Samples were desalted and concentrated using C-18 Sep-Pak (Waters) cartridges according to the manufacturer’s instructions. Peptides were eluted in 500 µL buffer B (35% ACN, 65% MilliQ H_2_O, 0.1% TFA). Eluted peptides were dried using a vacuum concentrator (Speedvac, Eppendorf) and stored at -20°C until analysis by mass spectrometry (MS). Prior to MS analysis, dried peptides were resuspended in buffer A (98% MilliQ H_2_O, 2% ACN, 0.1% TFA).

### Liquid chromatography-tandem mass spectrometry (LC-MS/MS) and data analysis

LC-MS/MS analysis was performed using a Dionex Ultimate 3000 nano-ultra high-pressure reverse-phase chromatography coupled on-line to a Q Exactive Orbitrap mass spectrometer (Thermo Scientific). Prior to introduction to the MS, peptides were separated on an EASY-Spray PepMap RSLC C18 column (500 mm × 75 μm, 2 μm particle size, Thermo Scientific) over a 60-minute gradient of 2-35% acetonitrile in 5% dimethyl sulfoxide, 0.1% formic acid at 250 nL/minute. The mass spectrometer was operated in data-dependent acquisition mode for automated switching between MS and MS/MS acquisition. Full MS survey scans were acquired from *m/z* 400-2,000 at a resolution of 70,000 at *m/z* 200 and AGC target of 3e^6^ ions for a maximum injection time of 100 ms. The top 15 most abundant precursor ions were selected for high collision dissociation fragmentation after isolation with a mass window of 1.6 Th and at a resolution of 17,500, with a maximum injection time of 128 ms. The normalised collision energy was 28%.

MS raw data were searched against the UniProtKB human sequence database (fused Uniprot/Trembl, 02/2020) and label-free quantitation (LFQ) was performed using MaxQuant Software (v1.6.10.43). Search parameters were set to include carbamidomethyl (C) as a fixed modification, with oxidation (M) and protein N-terminal acetylation included as variable modifications. A maximum of 2 missed cleavages were allowed, with matching between runs. LFQ was performed using unique peptides only. Label-free interaction data analysis was performed using Perseus (v1.6.10.43).

### Immunofluorescence staining of FFPE tissue sections

FFPE tissue sections (4 µM-thick) were dewaxed and rehydrated using a graded ethanol series (100%, 90%, 70%, methanol for 1 minute; final wash in ddH2O for 5 minutes). Before antibody labelling, antigens were unmasked by heat-induced epitope retrieval (HIER) using either 10 mM citrate solution (pH6) and/ or Tris ethylenediaminetetraacetic acid (TE) solution (pH9). Tissue sections were blocked with 5% normal goat serum in phosphate-buffered saline (PBS; blocking buffer) before incubating with primary antibodies either overnight at 4°C or for 2 hours at RT (for primary antibody details, see Supp. Table S1). The resulting antigen-antibody complexes were detected using Alexa Fluor-conjugated secondary antibodies (1/400; AlexaFluor 488, 555, 647 - secondary antibodies are listed in Supp. Table S1). Cell nuclei were stained with DAPI (1 μg/mL; Invitrogen, UK). Primary and secondary antibodies were diluted in DAKO REAL antibody diluent (Agilent, UK, cat. no. S202230-2). Tissue sections were washed three times in PBS to remove excess antibodies after each antibody incubation step. Sections were mounted in fluorescence mounting media (Agilent, UK, cat. no. S302380-2) before imaging. To create autofluorescence control slides, mouse brain sections were stained as above using the relevant Alexa Fluor-conjugated secondary antibodies (AlexaFluor 488, 555, 647) in the absence of primary antibodies.

### Lambda phosphatase pre-treatment of tissue sections

Following HIER, rTg4510 mouse brain sections were incubated in the absence (control) or presence (treated) of 10,000 U/mL λ phosphatase (New England Biolabs, cat. no. P0753) for 24 hours at RT, following manufacturer’s instructions. Tissue sections were then washed in PBS, and a second HIER in Citrate buffer (pH 6.0) was performed to cease phosphatase activity. Sections were then immunostained and mounted in fluorescence mounting media, as described above.

### Thioflavin S staining

Following immunofluorescence staining as described above, tissue sections were incubated with 0.5% Thioflavin S (diluted in water; Sigma Aldrich, cat. no. T1892) for 2 minutes, followed by submerging in 100% ethanol for a total of 10 times. Slides were then washed twice for 5 minutes in ddH_2_O before mounting in fluorescence mounting media, as above.

### Image acquisition, processing and quantifications

Low-magnification images of whole-tissue sections were obtained through scanning slides on the Akoya PhenoImager HT^TM^ Automated Quantitative Pathology Imaging system (CLS143455). Autofluorescence control slides (described above) were imaged to generate spectral libraries of the endogenous autofluorescence signatures detected in each imaging channel, in order to enable the removal of autofluorescence prior to image analysis. Regions of interest (ROIs) were subsequently re-imaged in high definition using the 40x objective and processed using the InForm^TM^ image analysis platform (Akoya Biosciences, US). The software allows for the generation of unbiased, spectrally unmixed images in which autofluorescence has been removed computationally based on the autofluorescence signature detected for mouse brain tissues in each channel. This is achieved using the autofluorescence slides described above and following the manufacturer’s instructions. Figures were created using the HALO® (Indica Labs) image analysis platform.

For all samples, staining was reviewed across the entire brain section with a particular focus on both the cortex and hippocampus. For rTg4510 mouse brains, signal was strongest in cortex (as neuronal loss in the hippocampus is pronounced in rTg4510 mice at this age), whereas images for region CA1 of the hippocampus are provided for wildtype, *Mapt^-/-^* and hTau mouse brains.

Quantification analysis of the immunofluorescent signal was performed using HALO® (Indica Labs), a gold standard image analysis platform for quantitative analysis of IHC data. The Highplex FL module was used to quantify the signal intensity obtained with Tau antibodies in rTg4510 mouse brain sections, that had either been untreated (control) or treated with λPP. For each image, the region of interest (cortex) was manually annotated. Efficiency of nuclear detection was assessed visually by comparison with the DAPI nuclear stain. The cytoplasm radius and cell size were set manually. Cells were categorised as either Tau-positive (low, medium or high) or Tau-negative. Statistical analyses to compare control (untreated) and λPP-treated sections was performed using the two-tailed, independent *t-test*.

## Results

We tested a total of 53 commercially available anti-Tau antibodies (hereafter referred to as Tau antibodies), that can be split into four broad classes: (i) “total” Tau antibodies, (ii) phospho-Tau antibodies, (iii) other PTM-dependent Tau antibodies and (iv) isoform-specific Tau antibodies. The term “total” Tau antibodies is used to denote reagents with the presumed ability to detect all Tau variants, regardless of phosphorylation status, aggregation status, or the presence of most other PTMs, in keeping with the nomenclature commonly encountered in the Tau research literature. These are sometimes also referred to as “pan” Tau antibodies. However, by definition, these antibodies cannot detect Tau species that lack the region containing the antibody epitope, so the “total Tau” denomination does not account for truncation. Unless otherwise specified, residue numbering is based on the 2N4R Tau splice isoform (441 amino acids-long). **Fig. 1C** summarise the antibodies used in this study and **Supp. Table S1** provides all relevant information, including, where known, the isotype, immunogen, target epitope and predicted species reactivity.

### Approaches to validating Tau antibodies by WB

Recombinant proteins as well as biological samples of murine and human origin were employed for the validation of Tau antibodies by WB (**Fig. 1D**). To aid data interpretation, we first present relevant information on the different experimental approaches:

#### 1. Overexpression of wildtype human Tau in HEK293T cells

To determine each antibody’s ability to detect Tau in complex biological samples, we generated cellular lysates where Tau is present at high levels. For this, we overexpressed wildtype, human Tau in HEK293T cells, a widely employed human cellular model for protein expression, owing to their faithful translation and processing of proteins and ability to generate most PTMs, including phosphorylation. To allow for unambiguous identification of the transgenic protein, we overexpressed human Tau (isoform 0N3R) tagged with a fluorescent protein (tdTomato) at either the N-terminus (tdTomato-N-Tau) or the C-terminus (Tau-C-tdTomato). Control cells expressed the respective linker and tdTomato protein alone (N-tdTomato ctrl and C-tdTomato ctrl, respectively). Using an anti-RFP antibody that detects the tdTomato fluorescent protein tag, we confirmed tdTomato expression at similar levels across all four cell lines (**Supp. Fig. S1A**). Bands corresponding to tdTomato alone (predicted MW of 54.2 kDa, observed at ∼60 kDa) were present not only in control cells, but also in Tau-overexpressing cells, suggesting that a proportion of the tdTomato tag is cleaved off (**Supp. Fig. S1A**). In fact, a faint band corresponding to the full-length tdTomato-tagged Tau (predicted MW 90.9 kDa, observed on WB just below 95 kDa) was only observed in tdTomato-N-Tau cells, indicating that the majority of the transgenic Tau protein is proteolytically processed (**Supp. Fig. S1A**).

As overexpression in HEK293T cells leads to high levels of protein expression, we utilised this system as a first-pass test to determine the ability of “total” Tau antibodies, and relevant isoform-specific Tau antibodies, to detect Tau in complex protein extracts (**Figs. 2-5, 10, 11**). Transgenic Tau expressed in mammalian cells, including in HEK293-derived cell lines, has been found to become extensively phosphorylated [77–79]. We therefore anticipated that many of the PTM-dependent Tau antibodies would also detect the transgenic protein in these samples, whilst allowing for the possibility that some Tau PTMs may not be generated in HEK293T cells cultured under standard conditions. In the case of PTM-dependent antibodies, a positive signal can, therefore, demonstrate an antibody’s ability to detect Tau, whereas lack of a signal does not distinguish between an antibody’s failure to detect the target protein or the absence of the relevant PTM(s) (**Figs. 6-9**).

**Figure 2.**
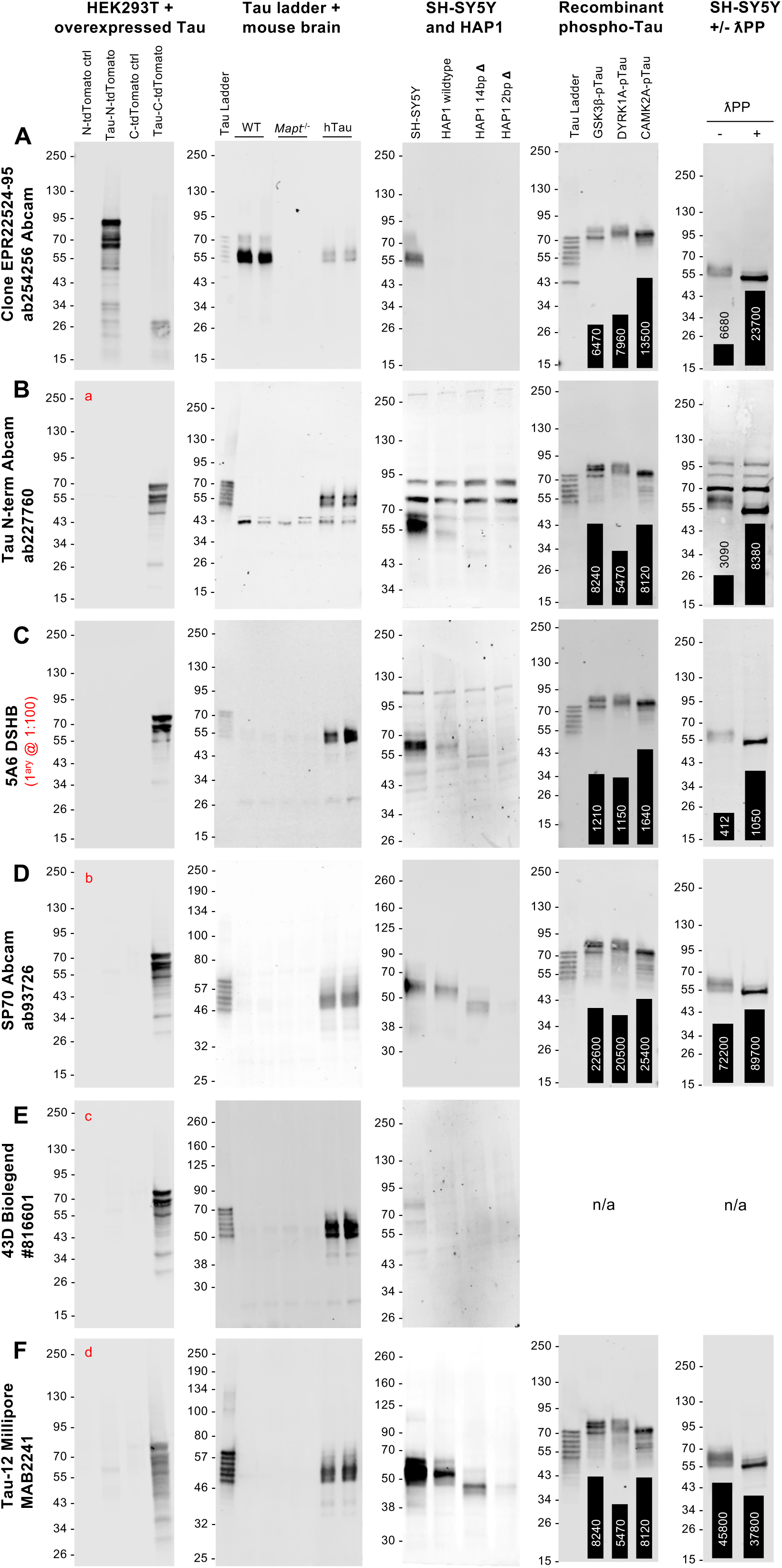
Validation of total Tau antibodies by WB (part 1). **A-F:** WBs of lysates from HEK293T cells overexpressing 0N3R human Tau and corresponding control cells (**first column**); recombinant human Tau ladder (5 ng/isoform/lane), plus adult mouse brain lysates from wildtype, *Mapt^-/-^* and hTau mice (**second column**); lysates from SH-SY5Y neuroblastoma cells, plus HAP1 cells, parental (wildtype) and two cell lines carrying either a 14 bp deletion (14 bp Δ) or a 2-bp deletion (2 bp Δ) in *MAPT* exon 4 (**third column**); recombinant human Tau ladder (50 ng/isoform/lane) plus recombinant 2N4R Tau that has been phosphorylated by one of three known Tau kinases: GSK3ꞵ, DYRK1A or CAMKIIA (**fourth column**); lysates from SH-SY5Y neuroblastoma cells that have been either untreated (-) or treated (+) with λPP (**fifth column**). For WBs in the fourth and fifth columns, quantifications of the Tau signal intensity for each lane are shown superimposed on each WB image as a bar chart, with the respective value [a.u.] printed on or above each bar of the chart. WB membranes shown in each panel (**row**) were probed with a different Tau antibody: clone EPR22524 Abcam #ab254256 (**A**), N-terminal-targeting antibody Abcam #227760 (**B**), clone 5A6 DSHB (**C**), clone SP70 Abcam #ab93726 (**D**), clone 43D BioLegend #816601 (**E**), clone Tau-12 Millipore #MAB2241 (**F**). In this and all subsequent figures, where applicable, lowercase red lettering found in the upper left corner of specific blots indicates that adjusting brightness/contrast display settings revealed additional bands. The corresponding WB image obtained with the altered settings is shown in **Supp. Fig. S3**. In this and all subsequent figures, where applicable, MW markers [kDa] are indicated on the left of each blot.

**Figure 3.**
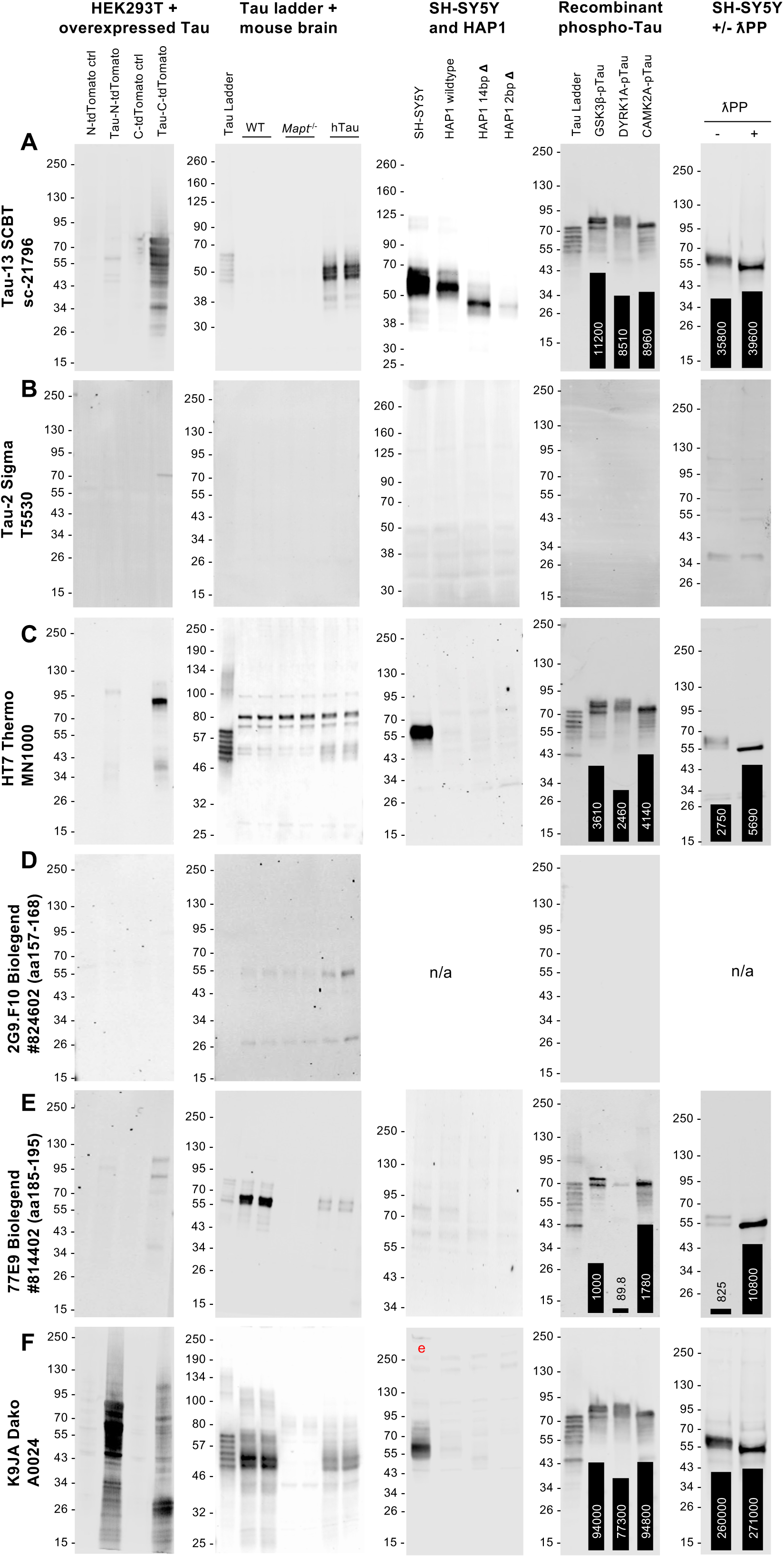
Validation of total Tau antibodies by WB (part 2). **A-F:** WBs of lysates from HEK293T cells overexpressing 0N3R human Tau and corresponding control cells (**first column**); recombinant human Tau ladder (5 ng/isoform/lane), plus adult mouse brain lysates from wildtype, *Mapt^-/-^* and hTau mice (**second column**); lysates from SH-SY5Y neuroblastoma cells, plus HAP1 cells, parental (wildtype) and two cell lines carrying either a 14 bp deletion (14 bp Δ) or a 2-bp deletion (2 bp Δ) in *MAPT* exon 4 (**third column**); recombinant human Tau ladder (50 ng/isoform/lane) plus recombinant 2N4R Tau that has been phosphorylated by one of three known Tau kinases: GSK3ꞵ, DYRK1A or CAMKIIA (**fourth column**); lysates from SH-SY5Y neuroblastoma cells that have been either untreated (-) or treated (+) with λPP (**fifth column**). For WBs in the fourth and fifth columns, quantifications of the Tau signal intensity for each lane are shown superimposed on each WB image as a bar chart, with the respective value [a.u.] printed on or above each bar of the chart. WB membranes shown in each panel (**row**) were probed with a different Tau antibody: clone Tau-13 Santa Cruz Biotechnology #sc-21796 (**A**), clone Tau-2 Sigma #T5530 (**B**), clone HT7 Thermo Fisher Scientific #MN1000 (**C**), clone 2G9.F10 BioLegend #824602 (**D**), clone 77E9 BioLegend #814402 (**E**), clone K9JA Dako #A0024 (**F**).

**Figure 4.**
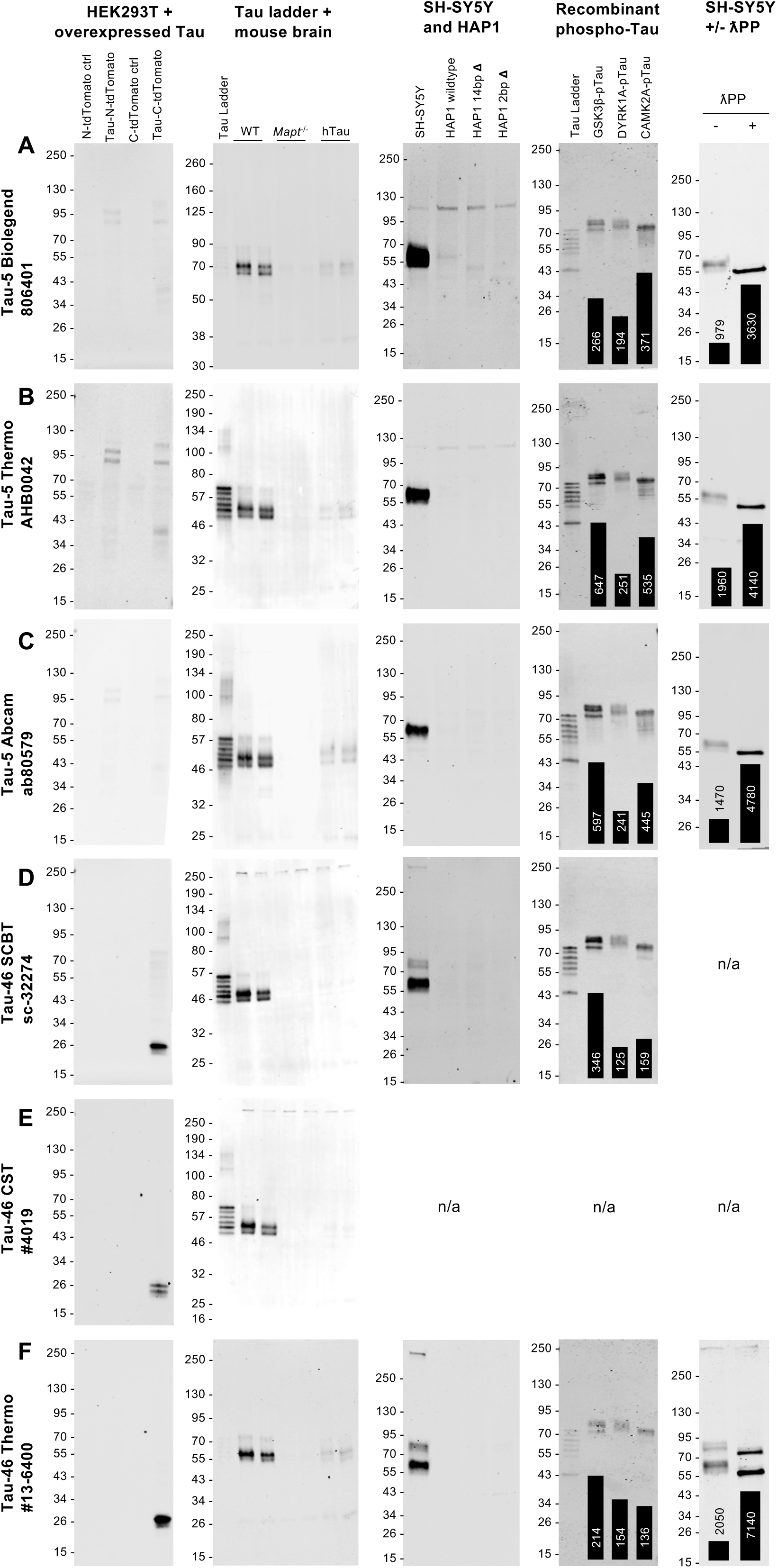
Validation of total Tau antibodies by WB (part 3). **A-F:** WBs of lysates from HEK293T cells overexpressing 0N3R human Tau and corresponding control cells (**first column**); recombinant human Tau ladder (5 ng/isoform/lane), plus adult mouse brain lysates from wildtype, *Mapt^-/-^* and hTau mice (**second column**); lysates from SH-SY5Y neuroblastoma cells, plus HAP1 cells, parental (wildtype) and two cell lines carrying either a 14 bp deletion (14 bp Δ) or a 2-bp deletion (2 bp Δ) in *MAPT* exon 4 (**third column**); recombinant human Tau ladder (50 ng/isoform/lane) plus recombinant 2N4R Tau that has been phosphorylated by one of three known Tau kinases: GSK3ꞵ, DYRK1A or CAMKIIA (**fourth column**); lysates from SH-SY5Y neuroblastoma cells that have been either untreated (-) or treated (+) with λPP (**fifth column**). For WBs in the fourth and fifth columns, quantifications of the Tau signal intensity for each lane are shown superimposed on each WB image as a bar chart, with the respective value [a.u.] printed on or above each bar of the chart. WB membranes shown in each panel (**row**) were probed with a different Tau antibody: clone Tau-5 BioLegend #806401 (**A**), clone Tau-5 Thermo Fisher Scientific #AHB0042 (**B**), clone Tau-5 Abcam #ab80579 (**C**), clone Tau-46 Santa Cruz Biotechnology #sc-32274 (**D**), clone Tau-46 Cell Signalling Technologies #4019 (**E**), clone Tau-46 Thermo Fisher Scientific #13-6400 (**F**).

**Figure 5.**
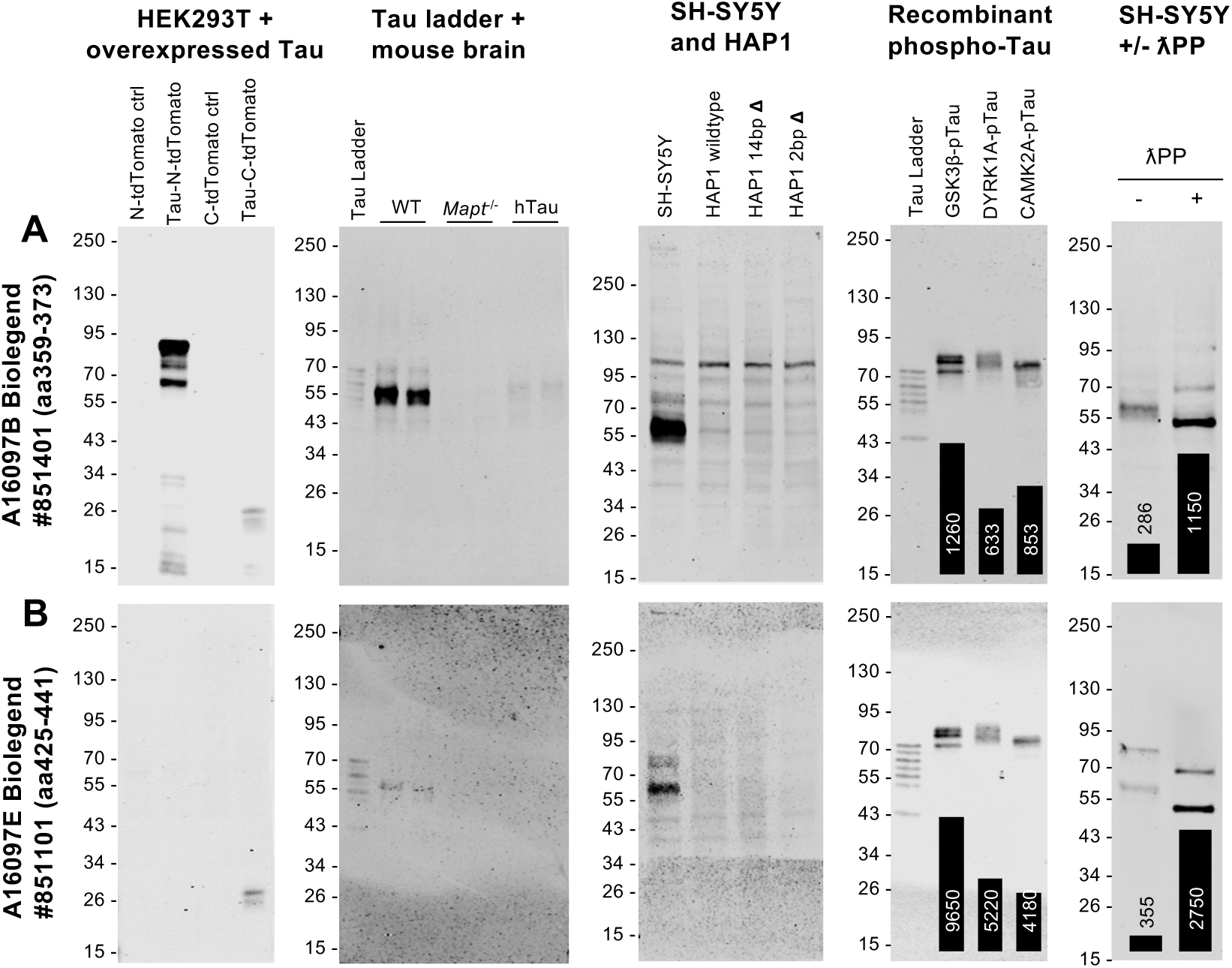
Validation of total Tau antibodies by WB (part 4). **A, B:** WBs of lysates from HEK293T cells overexpressing 0N3R human Tau and corresponding control cells (**first column**); recombinant human Tau ladder (5 ng/isoform/lane), plus adult mouse brain lysates from wildtype, *Mapt^-/-^* and hTau mice (**second column**); lysates from SH-SY5Y neuroblastoma cells, plus HAP1 cells, parental (wildtype) and two cell lines carrying either a 14 bp deletion (14 bp Δ) or a 2-bp deletion (2 bp Δ) in *MAPT* exon 4 (**third column**); recombinant human Tau ladder (50 ng/isoform/lane) plus recombinant 2N4R Tau that has been phosphorylated by one of three known Tau kinases: GSK3ꞵ, DYRK1A or CAMKIIA (**fourth column**); lysates from SH-SY5Y neuroblastoma cells that have been either untreated (-) or treated (+) with λPP (**fifth column**). For WBs in the fourth and fifth columns, quantifications of the Tau signal intensity for each lane are shown superimposed on each WB image as a bar chart, with the respective value [a.u.] printed on or above each bar of the chart. WB membranes shown in each panel (**row**) were probed with a different Tau antibody: A16097B BioLegend #851401(**A**), A16097E BioLegend #851101 (**B**).

While HEK293 and derived cell lines are sometimes assumed not to express Tau [80–83], endogenous Tau expression has been detected in many non-neuronal cell lines, including both HEK293 and HEK293T, at the mRNA and protein levels [84–87]. In line with this, we find that HEK293T cells used in this study express low levels of endogenous Tau, which was detectable when loading 50 µg protein lysate/lane on SDS-PAGE (**Sup. Fig. S1N**). However, we note that endogenous Tau expression remained largely undetectable on WB in lysates from cells overexpressing tdTomato-tagged Tau (**Figs. 2-11**), as only low amounts of protein lysate (6 µg/lane) were required for the detection of transgenic Tau, given that this was expressed at much higher levels than the endogenous protein.

#### 2. Purified recombinant human Tau

A commercial recombinant human Tau ladder, containing equimolar amounts of the six common Tau splice isoforms (0N3R, 0N4R, 1N3R, 1N4R, 2N3R and 2N4R), served as a second positive control for total and isoform-specific Tau antibodies, as well as for the dephosphorylation-dependent Tau-1 antibody clone (**Figs. 2-5, 9-11**). Conversely, as recombinant Tau proteins produced in bacteria are devoid of PTMs, the Tau ladder served as a negative control for the remaining PTM-dependent antibodies (**Figs. 6-9**). As detailed in Methods, the Tau ladder used on some of the WBs contained a seventh Tau isoform of a MW ∼10 kDa lower than 0N3R. In addition, we also included recombinant 2N4R Tau that has been phosphorylated with one of three known Tau kinases: GSK3β, DYRK1A or CAMK2A, respectively (**Figs. 2-11**).

**Figure 6.**
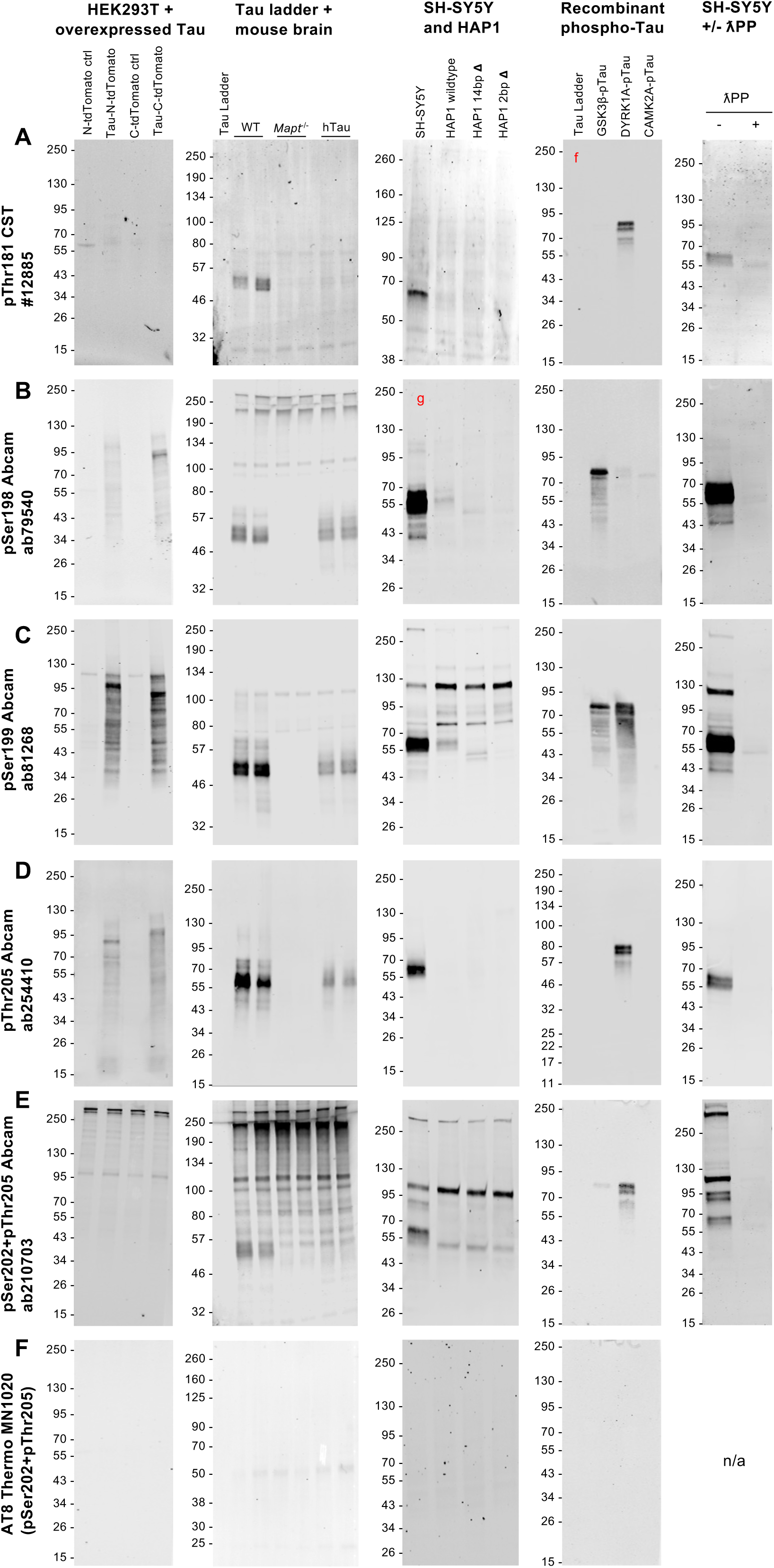
Validation of phospho-Tau antibodies by WB (part 1). **A-F:** WBs of lysates from HEK293T cells overexpressing 0N3R human Tau and corresponding control cells (**first column**); recombinant human Tau ladder (5 ng/isoform/lane), plus adult mouse brain lysates from wildtype, *Mapt^-/-^* and hTau mice (**second column**); lysates from SH-SY5Y neuroblastoma cells, plus HAP1 cells, parental (wildtype) and two cell lines carrying either a 14 bp deletion (14 bp Δ) or a 2-bp deletion (2 bp Δ) in *MAPT* exon 4 (**third column**); recombinant human Tau ladder (50 ng/isoform/lane) plus recombinant 2N4R Tau that has been phosphorylated by one of three known Tau kinases: GSK3ꞵ, DYRK1A or CAMKIIA (**fourth column**); lysates from SH-SY5Y neuroblastoma cells that have been either untreated (-) or treated (+) with λPP (**fifth column**). For WBs in the fourth and fifth columns, quantifications of the Tau signal intensity for each lane are shown superimposed on each WB image as a bar chart, with the respective value [a.u.] printed on or above each bar of the chart. WB membranes shown in each panel (**row**) were probed with a different Tau antibody: pThr181 Cell Signalling Technologies #12885 (**A**), pSer198 Abcam #ab79540 (**B**), pSer199 Abcam #81268 (**C**), pThr205 Abcam #254410 (**D**), pSer202+pThr205 Abcam #210703 (**E**), AT8 Thermo Fisher Scientific #MN1020 (**F**).

**Figure 7.**
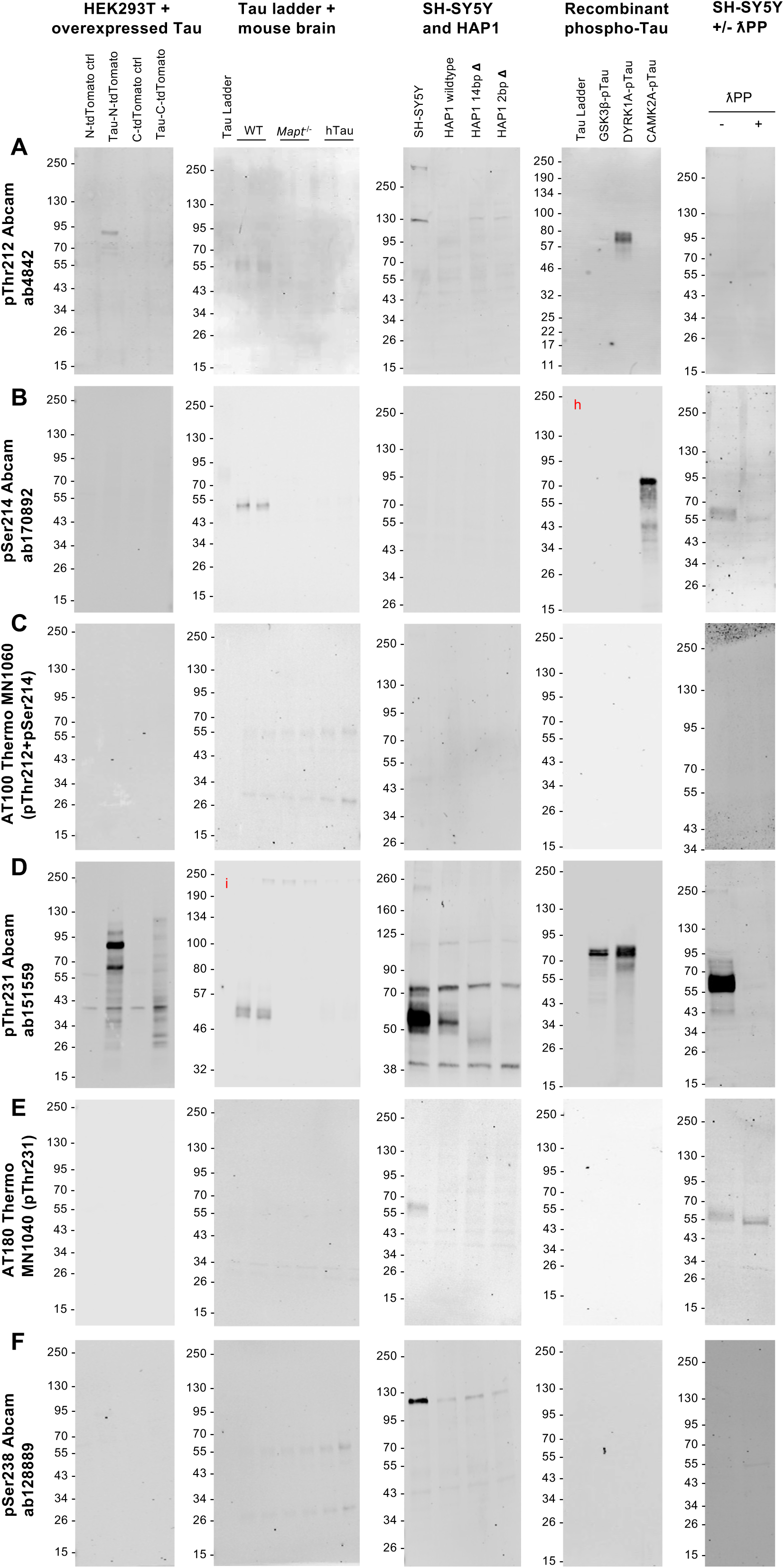
Validation of phospho-Tau antibodies by WB (part 2). **A-F:** WBs of lysates from HEK293T cells overexpressing 0N3R human Tau and corresponding control cells (**first column**); recombinant human Tau ladder (5 ng/isoform/lane), plus adult mouse brain lysates from wildtype, *Mapt^-/-^* and hTau mice (**second column**); lysates from SH-SY5Y neuroblastoma cells, plus HAP1 cells, parental (wildtype) and two cell lines carrying either a 14 bp deletion (14 bp Δ) or a 2-bp deletion (2 bp Δ) in *MAPT* exon 4 (**third column**); recombinant human Tau ladder (50 ng/isoform/lane) plus recombinant 2N4R Tau that has been phosphorylated by one of three known Tau kinases: GSK3ꞵ, DYRK1A or CAMKIIA (**fourth column**); lysates from SH-SY5Y neuroblastoma cells that have been either untreated (-) or treated (+) with λPP (**fifth column**). For WBs in the fourth and fifth columns, quantifications of the Tau signal intensity for each lane are shown superimposed on each WB image as a bar chart, with the respective value [a.u.] printed on or above each bar of the chart. WB membranes shown in each panel (**row**) were probed with a different Tau antibody: pThr212 Abcam #ab4842 (**A**), pSer214 Abcam #ab170892 (**B**), AT100 Thermo Fisher Scientific #MN1060 (**C**), pThr231 Abcam #ab151559 (**D**), AT180 Thermo Fisher Scientific #MN1040 (**E**), pSer238 Abcam #ab128889 (**F**).

**Figure 8.**
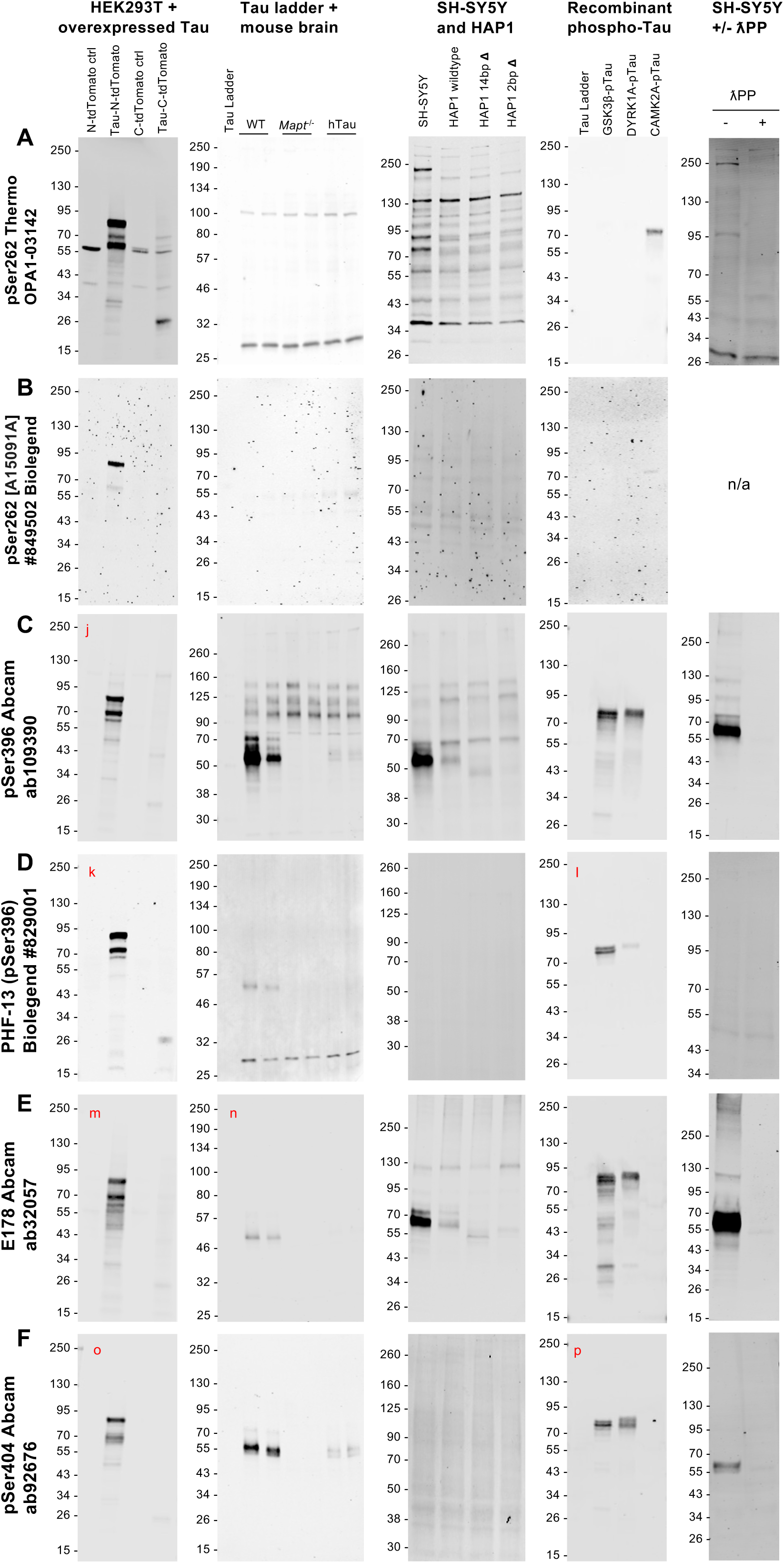
Validation of phospho-Tau antibodies by WB (part 3). **A-F:** WBs of lysates from HEK293T cells overexpressing 0N3R human Tau and corresponding control cells (**first column**); recombinant human Tau ladder (5 ng/isoform/lane), plus adult mouse brain lysates from wildtype, *Mapt^-/-^* and hTau mice (**second column**); lysates from SH-SY5Y neuroblastoma cells, plus HAP1 cells, parental (wildtype) and two cell lines carrying either a 14 bp deletion (14 bp Δ) or a 2-bp deletion (2 bp Δ) in *MAPT* exon 4 (**third column**); recombinant human Tau ladder (50 ng/isoform/lane) plus recombinant 2N4R Tau that has been phosphorylated by one of three known Tau kinases: GSK3ꞵ, DYRK1A or CAMKIIA (**fourth column**); lysates from SH-SY5Y neuroblastoma cells that have been either untreated (-) or treated (+) with λPP (**fifth column**). For WBs in the fourth and fifth columns, quantifications of the Tau signal intensity for each lane are shown superimposed on each WB image as a bar chart, with the respective value [a.u.] printed on or above each bar of the chart. WB membranes shown in each panel (**row**) were probed with a different Tau antibody: pSer262 Thermo Fisher Scientific #OPA1-03142 (**A**), pSer262 (clone A15091A) BioLegend #894502 (**B**), pSer396 Abcam #ab109390 (**C**), clone PHF-13 BioLegend #829001 (**D**), clone E178 Abcam #ab32057 (**E**), pSer404 Abcam #ab92676 (**F**).

**Figure 9.**
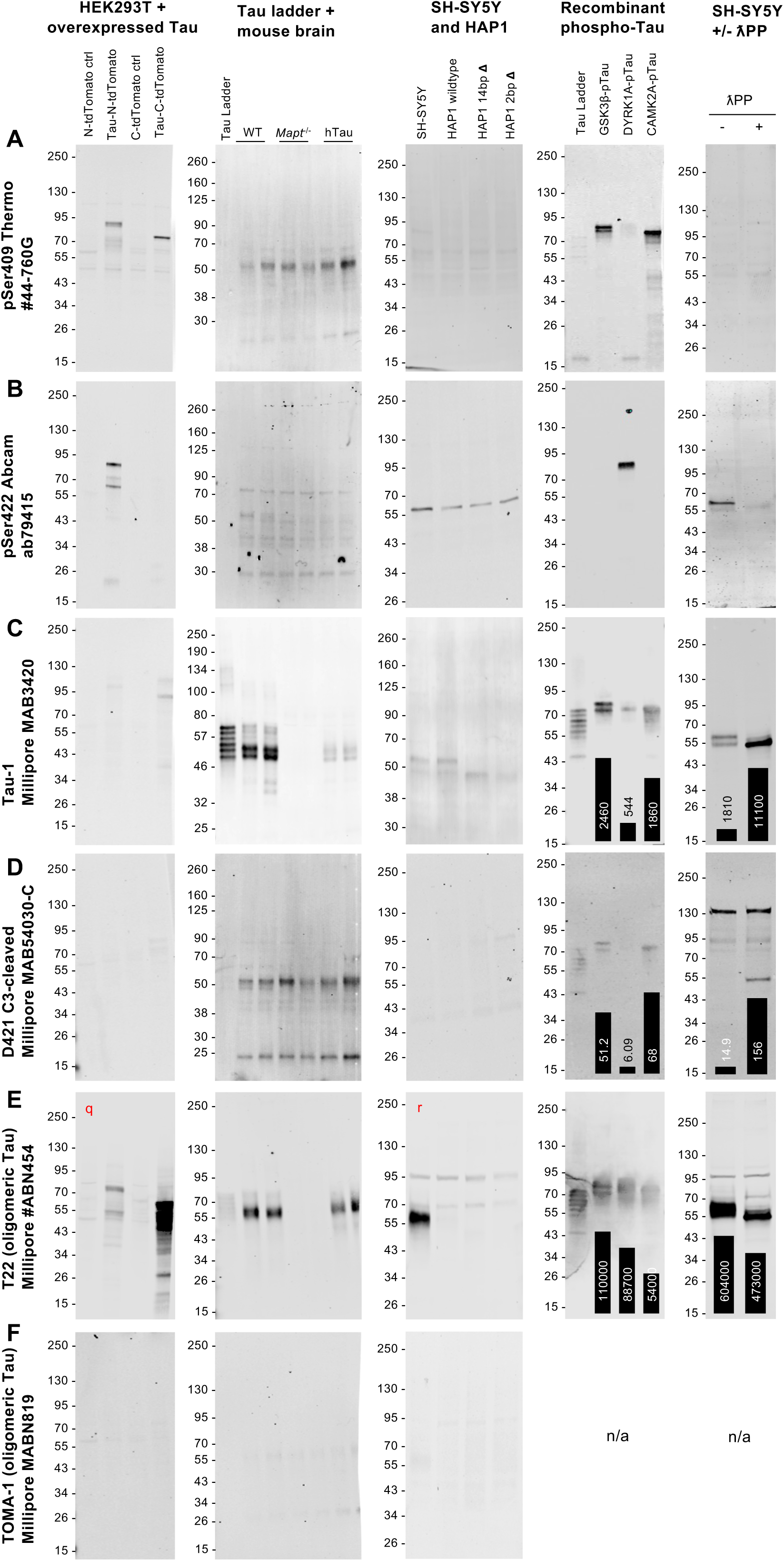
Validation of phospho-Tau antibodies (part 4) and other PTM-dependent antibodies by WB. **A-F:** WBs of lysates from HEK293T cells overexpressing 0N3R human Tau and corresponding control cells (**first column**); recombinant human Tau ladder (5 ng/isoform/lane), plus adult mouse brain lysates from wildtype, *Mapt^-/-^* and hTau mice (**second column**); lysates from SH-SY5Y neuroblastoma cells, plus HAP1 cells, parental (wildtype) and two cell lines carrying either a 14 bp deletion (14 bp Δ) or a 2-bp deletion (2 bp Δ) in *MAPT* exon 4 (**third column**); recombinant human Tau ladder (50 ng/isoform/lane) plus recombinant 2N4R Tau that has been phosphorylated by one of three known Tau kinases: GSK3ꞵ, DYRK1A or CAMKIIA (**fourth column**); lysates from SH-SY5Y neuroblastoma cells that have been either untreated (-) or treated (+) with λPP (**fifth column**). For WBs in the fourth and fifth columns, quantifications of the Tau signal intensity for each lane are shown superimposed on each WB image as a bar chart, with the respective value [a.u.] printed on or above each bar of the chart. WB membranes shown in each panel (**row**) were probed with a different Tau antibody: pSer409 Thermo Fisher Scientific #44-760G (**A**), pSer422 Abcam #ab79415 (**B**), clone Tau-1 Millipore #MAB3420 (**C**), clone D421 Millipore #MAB54030-C (**D**), clone T22 Millipore #ABN454 (**E**), TOMA-1 Millipore #MABN819 (**F**).

#### 3. Genetic manipulation of endogenous Tau

Antibody specificity can vary substantially depending on the species being tested. Thus, fully addressing antibody specificity ideally requires the use of samples originating from the species of interest, but from which the relevant gene has been knocked out such that the target protein is ablated. In the absence of such tissue, samples in which the expression of the target protein is specifically decreased (e.g. by genetic knock-down approaches) can also be employed. As a first test for antibody specificity, we employed mouse brain lysates from wildtype mice, *Mapt^-/-^* mice [73] and humanised Tau mice (hTau; [74]) (**Figs. 2-11**). The latter express human Tau at physiological levels on a mouse *Mapt^-/-^* background.

To test antibody specificity in samples of human origin, we made use of commercially-available HAP1 cells, which have been edited via Clustered Regularly Interspaced Short Palindromic Repeats (CRISPR)/Cas9 technology to generate cell lines that carry either a 2 bp or a 14 bp deletion in *MAPT* exon 4, a constitutively-included exon in all known Tau splice isoforms (**Fig. 1A, Figs. 2-11**). As the deletions introduced in this manner lead to out-of-frame shifts accompanied by the introduction of early stop codons, these two cell lines are predicted to lack expression of the Tau protein. Therefore, only bands that result from non-specific cross-reactivity were expected when analysing lysates from *MAPT*-edited HAP1 cells compared to parental HAP1 cells by WB. However, preliminary experiments revealed that Tau-immunoreactive bands continued to be detected in *MAPT*-edited cells but were shifted to a lower MW compared to parental cells, suggesting that the presumed “knockout” cells continue to produce a lower MW version of Tau (**Figs. 2-3**). To test this possibility, we used antibodies to immunoprecipitate (IP) Tau from parental and 2 bp-deletion cells. In the latter cell line, Tau expression is nearly undetectable by WB. Mass-spec analysis of the IP eluates identified Tau peptides in both cell lines, confirming that wildtype HAP1 cells express Tau and demonstrating that the presumed knockout *MAPT*-edited HAP1 cells continue to express Tau protein, corroborating our WB observations (**Supp. Fig. S2**). Despite this, HAP1 cells proved informative for antibody validation by WB due to the shift in MW displayed by Tau bands following *MAPT* editing, which allowed us to distinguish between specific and non-specific Tau-immunoreactive bands. In consequence, bands that were detected with Tau antibodies in HAP1 cell lysates were deemed non-specific if their MW did not change between *MAPT*-edited compared to parental cell lines. As wildtype, parental HAP1 cells express only relatively low levels of endogenous Tau, this provided the opportunity to identify antibodies that are both specific and sufficiently sensitive to detect Tau in samples where it is present at much lower levels than in neuronal/brain samples.

#### 4. Human neuroblastoma cell line +/- phosphatase treatment

Alongside HAP1 cells, we also analysed whole-cell lysates from SH-SY5Y, a widely-utilised human neuroblastoma cell line whose Tau expression is well documented [88–90]. Endogenous Tau expressed in SH-SY5Y cells is phosphorylated at multiple residues, including Ser199/Ser202, Thr231, Ser396 and Ser404 [91,92], thus making these a useful model for Tau antibody testing. To establish the requirement for phosphorylation and/or the impact of phosphorylation on antibody binding, SH-SY5Y lysates were treated with lambda phosphatase (λPP) to dephosphorylate proteins. λPP-treated lysates were tested alongside untreated, control lysates.

### Presentation of the WB data in main figures

In **Figs. 2-11**, each panel (row) shows WBs probed with a different Tau antibody (antibody name, supplier and catalog number are indicated on the left) with the following WBs being shown from left to right: lysates from HEK293T cells overexpressing Tau and respective control cells (**1^st^ column**); recombinant Tau ladder (5 ng/isoform) plus brain lysates from wildtype, *Mapt^-/-^* and hTau mice (**2^nd^ column**); lysates from SH-SY5Y neuroblastoma cells, plus lysates from parental and *MAPT*-edited HAP1 cells (**3^rd^ column**); recombinant Tau ladder (50 ng/isoform) and phosphorylated recombinant Tau (**4^th^ column**); lysates from SH-SY5Y neuroblastoma cells +/- λPP (**5^th^ column**). All WB images are provided uncropped. Where adjusting image brightness/contrast revealed additional bands, whether specific or non-specific, this is indicated using red lettering in the top left corner of the relevant WB image and the adjusted image is shown in **Supp. Fig. S3**.

Specificity of the secondary antibodies was confirmed by omitting the primary antibody and probing WB membranes with each secondary antibody individually (**Supp. Fig. S1**). Equal protein loading was confirmed by probing blots with an anti-GAPDH antibody (**Supp. Fig. S1 B, G, H, M**). Of note, when tested on mouse brain samples, in most cases, we did not observe prominent cross-reactivity of anti-mouse secondary antibodies with endogenous immunoglobulins (IgGs), the heavy chain of which is expected to overlap with the specific Tau signal, and which have previously been reported to interfere with the detection of Tau by WB [69].

### Analysis of PTM-independent “total” Tau antibodies by WB

Of the 20 “total Tau” antibodies tested by WB (**Fig. 1**; **Supp. Table S1**), only two (clones Tau-2 and 2G9.F10) failed to detect specific Tau bands in any of the samples utilised in this study (**Figs. 2-5**). The remainder of total Tau antibodies all detected the target protein: recombinant version, when overexpressed in HEK293T cells, as well as Tau expressed at physiological levels in mouse brain lysates and in SH-SY5Y cells (**Figs. 2-5**). The inclusion of brain lysates from *Mapt^-/-^* mice revealed that antibodies #ab227760, HT7, K9JA and Tau-46 cross-reacted with prominent non-specific bands in samples of mouse origin, but only those detected by HT7 were of MWs that overlapped with the specific Tau signal (**Figs. 2-4**). Faint non-specific bands were also observed with antibodies 5A6, 43D and 2G9.F10 at ∼55 and ∼26 kDa in mouse brain lysates (**Figs. 2, 3**), and may correspond to the heavy and light chains of endogenous mouse IgGs, respectively [69]. In our testing, these did not interfere with detection of the specific Tau signal, owing to their relatively weak intensity. The remaining total Tau antibodies tested (#ab254256, SP70, Tau-12, Tau-13, 77E9, Tau-5, A16097B and A16097E) detected Tau with high specificity when used to probe mouse brain samples by WB (**Figs. 2-5**). Through comparison with bands detected in lysates from *MAPT*-edited HAP1 cells, we further established that antibodies #ab227760, 5A6, K9JA, Tau-5 BioLegend, Tau-5 Thermo and A16097B reacted with Tau but also with non-specific bands when used to probe human cell lysates on WB (**Figs. 2-5**). In the case of K9JA, non-specific bands were most obvious under settings required to detect the low levels of Tau expressed in HAP1 cells (**Supp. Fig. S3e**), suggesting that these may not interfere with the detection of Tau in human samples or cell types that express Tau at higher levels. As detected with the SP70, Tau-12 and Tau-13 antibodies, Tau protein levels in parental wildtype HAP1 cells were ∼35-50% of those in SH-SY5Y cells (**Supp. Fig. S4**). However, we found that not all of the antibodies that detected Tau in SH-SY5Y cell lysates were able to detect the lower levels of Tau expression in HAP1 cells. Specifically, #ab254256, 43D, HT7, Tau-5, Tau-46, A16097E and A16097B all failed to detect Tau in HAP1 cell lysates (**Figs. 2-5**), highlighting the importance of antibody choice when analysing endogenous Tau in non-brain/non-neuronal samples. Taken together, our data indicate that the #ab254256, SP70, 43D, Tau-12, Tau-13, HT7, 77E9 and Tau-5 Abcam total Tau antibodies detect the target protein with high specificity on WB when used to probe lysates of human origin (**Figs. 2-5**).

In the case of the Tau-5 and Tau-46 antibody clones, aliquots purchased from three different suppliers were tested in parallel (**Fig. 4**). While all six reagents detected Tau in wildtype mouse brain lysates, two of the six reacted with the recombinant Tau ladder weakly (Tau-46 Thermo Fisher Scientific #13-6400, and Tau-5 BioLegend #806401). Of note, among the three Tau-46 antibodies tested, two (SCBT and CST) reacted strongly with the recombinant human Tau ladder but failed to detect human Tau in extracts from hTau mouse brains. In contrast, the third Tau-46 antibody (Thermo) detected the recombinant Tau ladder only weakly but gave a clear signal for hTau mouse brain samples, Similarly, among the three Tau-5 reagents tested, the ones purchased from Thermo and Abcam detected the Tau ladder strongly, while only a weak signal was obtained with the one purchased from Biolegend. These observations caution that differences may exist even when “the same” antibody clone is sourced from different manufacturers.

By definition, the binding of total Tau antibodies should be unaffected by PTMs. As such, total Tau antibodies are expected to produce comparable immunoreactive signals to all three recombinant phospho-Tau variants described above. In contrast to this prediction, quantifications of the signal intensity detected for each lane revealed that many antibodies displayed differential reactivity with recombinant Tau that has been phosphorylated by different kinases (**Figs. 2-5, fourth column**). Furthermore, quantifying the intensity of the specific Tau bands detected with total Tau antibodies in untreated versus λPP-treated SH-SY5Y lysates, revealed that, for many of the antibodies, the signal intensity increased following dephosphorylation of the samples (**Figs. 2-5, fifth column**), with this increase being as large as 7.7-fold for clone A16097E (**Fig. 5B, fifth column**) and ∼13-fold for clone 77E9 (**Fig. 3E, fifth column**). Immunoreactivity of the #ab254256, #ab227760, 5A6, HT7, Tau-5, Tau-46 and A16097B antibodies was also enhanced following dephosphorylation of SH-SY5Y protein extracts with λPP (**Figs. 2-5, fifth column**). Conversely, our data suggest that the antibodies least affected by Tau phosphorylation status were the N-terminal antibodies SP70, Tau-12 and Tau-13, as well as the polyclonal K9JA raised against the protein’s C-terminal half (**Figs. 2-3, fourth and fifth columns**). Of note, none of the antibodies tested showed higher immunoreactivity in untreated compared to λPP-treated lysates (**Figs. 2-5, fifth column**), suggesting that none of these reagents has higher affinity for phosphorylated Tau. Taken together, our results demonstrate that binding of many “total” Tau antibodies is partially inhibited by phosphorylation, despite the assumption that these reagents bind Tau with equal affinity, irrespective of PTM occupancy.

### Analysis of phosphorylation-dependent Tau antibodies by WB

During development and in tauopathies, Tau is hyperphosphorylated at many different sites [17,93,94], which has spurred the development of a plethora of putative phospho-Tau antibodies. We tested 20 commercial phospho-Tau antibodies (**Fig. 1; Supp. Table S1**), of which 17 were developed to detect Tau phosphorylated at single residues, while three are expected to detect double phosphorylation events. This category includes the E178 antibody clone which was marketed as a total Tau antibody until 2021 and has been used as such in several studies to date (E178 has been cited in at least 72 published studies [95]), further highlighting the importance of detailed antibody validation.

It has previously been reported that several phospho-Tau antibodies can also react with the unphosphorylated peptide when tested on peptide arrays [70], thus raising concerns over their specificity for phosphorylated Tau. Of the 20 phospho-Tau antibodies tested here, only the pSer409 antibody reacted weakly with the unphosphorylated Tau ladder, with cross-reactivity only observed when high levels of the Tau ladder were loaded on SDS-PAGE (50 ng/isoform/lane) (**Fig. 9A, fourth column**). These data confirm that the remaining 19 antibodies do not detect unmodified Tau by WB (**Figs. 6-9**).

Thirteen of the 20 phospho-Tau antibodies tested (pSer198, pSer199, pThr205, pThr212, pThr231, 2 x pSer262, pSer396, PHF-13, E178, pSer404, pSer409 and pSer422) detected the protein in Tau-overexpressing HEK293T cell lysates (**Figs. 6-9, first column**), demonstrating that the transgenic protein is extensively phosphorylated, in agreement with previous reports [96–100]. The remaining seven of 20 phospho-Tau antibodies tested (pThr181, pSer202+pThr205, AT8, AT100, pSer214, AT180 and pSer238) did not detect transgenic Tau overexpressed in HEK293T cells (**Figs. 6-9, first column**), potentially due to the protein not being phosphorylated at the relevant epitopes in this experimental system, under the routine cell culture conditions employed in this study. Some phospho-Tau antibodies (pSer199, pSer202+pThr205, pThr231, pSer262, pSer409) also reacted with proteins present in control HEK293T cells (**Figs. 6-9, first column**), but it was not possible to ascertain whether this was indicative of non-specific antibody binding or due to the detection of endogenous Tau. Endogenous Tau expression has previously been reported in both HEK293 and HEK293T cells, at the mRNA and protein levels [78,101–103], but the levels of the endogenous protein are likely to be much lower than those of the overexpressed protein.

In mouse brain lysates, five antibodies (pThr181, pSer202+pThr205, pSer214, pThr231 and PHF-13) immunoreacted with specific Tau bands in wildtype but not hTau brain lysates, six antibodies (pSer198, pSer199, pThr205, pSer396, E178 and pSer404) detected specific Tau bands in both wildtype and hTau brain lysates, while the remaining nine (AT8, pThr212, AT100, AT180, pSer238, 2 x pSer262, pSer409, pSer422) did not produce a specific signal in either (**Figs. 6-9**). These data suggest that human Tau expressed in hTau mouse brains is phosphorylated at some but not all of the same sites as endogenous murine Tau. Prominent non-specific bands that migrated in the 46-70 kDa MW range, where specific Tau bands are expected, were observed only with the pSer202+pTh205 and pSer409 antibodies, while faint non-specific bands were also present in this region on WBs probed with the AT8, AT100, pSer238 and pSer422 antibodies. Outside of the expected Tau MW range, prominent non-specific bands were observed with the pSer198, pSer202+pThr205, pSer262 (ThermoFisher), pSer396 and PHF-13 antibodies, with faint bands also observed when probing with the pSer199, pSer409 and pSer422 antibodies. Comparison with lysates from *Mapt^-/-^* mice suggested that antibodies pThr205, AT8, pThr212, pSer214, AT100, pThr231, AT180, pSer262 (BioLegend), E178 and pSer404 are highly specific for use in combination with samples of mouse origin on WB.

A pattern of Tau phosphorylation similar to that of wildtype mouse brain was also observed in SH-SY5Y human neuroblastoma cells, where Tau-immunoreactive bands were detected with nine antibodies (pThr181, pSer198, pSer199, pThr205, pSer202+pThr205, pThr231, AT180, pSer396 and E178) (**Figs. 6-9, third column**). By comparison with HAP1 cells, specificity of Tau-immunoreactive band(s) in the 50-70 kDa MW range was confirmed for all nine antibodies. However, we note that two of these antibodies (pSer202+pThr205 and pSer396) also reacted with additional non-specific bands that migrate in the 50-70 kDa MW range, and that five of these antibodies (pSer199, pSer202+pThr205, pThr231, E178 and pSer396) reacted with prominent non-specific bands outside the 50-70 kDa MW range, thus suggesting that further testing may be required if using these reagents in assays where the MW of the proteins being detected in not known. For antibodies AT8, pThr212, pSer214, AT100, PHF-13, pSer404 and pSer409, no immunoreactive bands were detected in any of the SH-SY5Y or HAP1 cell lysates, supporting the specificity of these reagents. In contrast, pSer262 detected multiple bands that were also present in lysates from *MAPT*-edited HAP1 cells (most obvious for the Thermo Fisher Scientific reagent; **Fig. 8A-B, third column**), suggesting that antibodies targeting this phospho-epitope display extensive non-specific cross-reactivity.

The requirement for phosphorylation was tested by treating SH-SY5Y protein extracts with λPP (**Figs. 6-9, fifth column**). This abrogated the signal detected with eight out of the nine antibodies that detected a signal in untreated SH-SY5Y lysates, thus confirming the phosphorylation-dependent nature of these antibodies’ epitopes. In contrast, the signal detected with the AT180 antibody did not decrease in λPP-treated protein extracts (**Fig. 7E, fifth column**), while the band displayed a downward shift in apparent MW, a characteristic of phosphorylation-independent antibodies. This may be interpreted as evidence that AT180 cross-reacts with the unphosphorylated protein. However, taken together with the fact that this antibody did not react with the recombinant Tau ladder (Fig. 7), an alternative explanation could be that it detects a λPP-resistant epitope (potentially dependent on conformation or other PTMs), which becomes better accessible following removal of inhibitory phosphorylation on nearby residues.

To further characterise phospho-Tau antibodies, we used recombinant 2N4R Tau that has been phosphorylated with one of three kinases (GSK3β, DYRK1A or CAMK2A). Sixteen of the 20 phospho-Tau antibodies reacted with at least one of the recombinant phosphorylated Tau versions tested but, with the exception of pSer198, none detected all three types of phosphorylated recombinant Tau (**Figs. 6-9, fourth column**). This is in agreement with the prediction that each kinase phosphorylates Tau at a specific pattern of residues. Antibody pThr212, which did not detect a clear, specific signal in any of the biological samples tested here, immunoreacted with DYRK1A-phosphorylated Tau, thus demonstrating its ability to detect phospho-Tau (**Fig. 7A, fourth column**). Four antibodies did not react with any of the three phosphorylated recombinant Tau versions: AT8, AT100, AT180 and pSer238. As AT8, AT100 and pSer238 also did not react with Tau in any of the biological samples tested, it was not possible to establish whether absence of signal was a consequence of these antibodies failing to bind their target epitope or indicative of Tau not being phosphorylated at the relevant residues in any of the samples tested here. For AT8 and AT100, failure to bind their respective epitopes is an unlikely explanation, given their widespread use to detect pathological Tau. However, as none of the samples tested here are expected to contain pathological Tau, lack of AT8 and AT100 immunoreactivity is not unexpected.

### Analysis of other PTM-dependent Tau antibodies by WB

Tau-1 is a dephosphorylation-dependent antibody [104,105], widely employed to detect Tau that is not phosphorylated at residues Ser195 to Thr205, located in the mid-domain of the Tau protein, a region that includes the AT8 epitope (**Fig. 1, Supp. Table S1**). Because of this, one common practice is to dephosphorylate samples in order to detect “total” Tau with the Tau-1 clone. Tau-1 reacted strongly with the unphosphorylated, recombinant Tau ladder, as well as with endogenous Tau from wildtype mouse brains, but, importantly, did not display detectable immunoreactivity with proteins in *Mapt^-/-^* mouse brain lysates, demonstrating its specificity when used in combination with murine protein extracts (**Fig. 9C, second column**). In lysates from Tau-overexpressing HEK293T cells (**Fig. 9C, first column**), SH-SY5Y cells and HAP1 cells (**Fig. 9C, third column**), this antibody detected only faint but specific bands, suggesting that the majority of Tau is phosphorylated at the Tau-1 epitope in these samples. Treating SH-SY5Y lysates with λPP resulted in a ∼6-fold increase in the intensity of the immunoreactive signal detected on WB, in agreement with the dephosphorylation-dependent nature of its epitope (**Fig. 9C, fifth column**). Taken together, our data support that Tau-1 detects the dephosphorylated versions of both murine and human Tau with high specificity.

Early stages of AD are characterised by the proteolytical cleavage of Tau at Asp421 by caspases, giving rise to ΔTau, which can adopt the MC1 pathological conformation and has the ability to “seed” aggregation [106]. Whether this event contributes to pathology or is a protective response to early pathological changes remains a matter of debate [106–108]. Given the particular biological relevance of this PTM, we tested the D421 antibody clone, which was raised against Tau cleaved at Asp421 [109] and has been reported to be highly specific. No specific D421-immunoreactivity was observed in lysates from HEK293T Tau-overexpressing cells, mouse brain, SH-SY5Y or HAP1 cells (**Fig. 9D**), suggesting that caspase-cleaved Tau is not present in the experimental models employed in this study. In mouse brain lysates, conspicuous non-specific bands were present at ∼55 and ∼25 kDa, presumed to represent endogenous IgGs (**Fig. 9D, second column**). However, we found that the D421 antibody detected a specific Tau band in SH-SY5Y lysates following λPP treatment (**Fig. 9D, fifth column**), an unexpected finding that suggests that phosphorylation inhibits its binding to Tau. While these data may be interpreted as evidence for the presence of caspase-cleaved Tau in SH-SY5Y cells, further testing revealed that D421 also immunoreacts weakly with full-length Tau (**Fig. 9D, fourth column**), a feature that only became apparent when higher amounts of recombinant Tau were loaded on SDS-PAGE (50 ng/isoform/lane on recombinant phospho-Tau WB, fourth column, compared to 5 ng/isoform/lane on mouse brain WB, second column). Furthermore, phosphorylation of recombinant Tau with DYRK1A strongly inhibited detection with the D421 antibody (**Fig. 9D, fourth column**). Taken together, our data demonstrate that binding of the D421 antibody to Tau is impaired by phosphorylation and further suggest that this antibody is not specific to Tau truncated at Asp421 as it also displayed immunoreactivity towards full-length, recombinant Tau.

Also targeting Tau species of pathological and diagnostic relevance, the T22 and the Tau oligomer-specific monoclonal antibody 1 (TOMA-1) antibody clones were raised by using *in vitro*-generated recombinant 2N4R Tau oligomers as immunogens [110,111]. No TOMA-1-immunoreactive bands were detected in any of the samples tested, thus supporting this antibody’s specificity, but offering no information with regards to its ability to detect Tau oligomers (**Fig. 9F**). In contrast, the T22 antibody reacted with the Tau ladder and detected monomeric Tau in lysates from HEK293T Tau-overexpressing cells, wildtype and hTau mouse brains, and SH-SY5Y cells (**Fig. 9E**). Immunoreactivity was maintained in SH-SY5Y lysates following λPP treatment (**Fig. 9E, fifth column**). Absence of T22-immunorecative bands in *Mapt^-/-^* mouse brain lysates supports this reagent’s specificity when used to detect Tau in mouse samples (**Fig. 9E, second column**), but non-specific bands observed at ∼70 and 95 kDa in SH-SY5Y and HAP1 cell lysates indicated non-specific cross-reactivity with proteins present in samples of human origin (**Fig. 9E, third column**). Taken together, our data demonstrate that T22 can detect monomeric Tau in a PTM-independent manner, thus demonstrating that it is not specific to Tau oligomers.

### Analysis of isoform-specific Tau antibodies by WB

The expression of Tau splice isoforms is regulated in a developmental-, tissue- and disease-dependent manner [112–119]. In the adult human brain, distortion of the ratios between different isoforms has been reported to correlate with and even to cause neurodegeneration [120–127]. There is, therefore, strong research interest in identifying antibodies that can be used to reliably detect and quantify the various Tau splice isoforms. In this category, we include antibodies purposefully developed to recognise specific Tau splice isoforms as well as antibodies that were identified as targeting certain splice isoforms in the course of this work. The former category comprises of antibodies against: 0N Tau, 2N Tau, 3R Tau (clone RD3) and four different antibodies raised against 4R Tau: Millipore clone 7D12.1, Millipore clone 1E1A6 (also known as RD4), BioLegend clone 5F9, and Abcam clone EPR21725 (Fig. 1; Supp. Table S1). For simplicity, antibodies in the latter category will hereafter be referred to as: Millipore-7D12, Millipore-RD4, BioLegend and Abcam 4R Tau antibodies, respectively. An antibody targeting 1N Tau (BioLegend #823902, clone 4H5.B9) was discontinued during the course of this work and was therefore excluded from this report. No alternative antibody against human 1N Tau isoforms is currently available commercially, to the best of our knowledge. A murine-specific 1N Tau antibody has been generated by an academic laboratory [128] but, to date, no antibodies have been described that target the human 1N Tau isoform.

The 0N Tau antibody detected a single band for 0N3R Tau-overexpressing HEK293T cells but only did so in cells expressing Tau-C-tdTomato, suggesting that placement of the tdTomato tag at the N terminus, closer to its epitope, may inhibit antibody binding and/or may promote proteolytic processing of the protein such that the relevant epitope is destroyed (**Fig. 10A, first column**). However, this antibody failed to react with its target isoforms in the recombinant Tau ladder (**Fig. 10A, second column**), even when 50 ng/isoform were loaded on SDS-PAGE (**Fig. 10A, fourth column**), nor did it detect Tau in mouse brain samples (**Fig. 10A, second column**) or in SH-SY5Y human cell lysates (**Fig. 10A, third column**), despite evidence that 0N Tau is the main Tau isoform expressed in both [89,129]. Taken together, our data suggest that the 0N antibody can detect its target protein, but only when this is overexpressed/present at very high levels.

**Figure 10.**
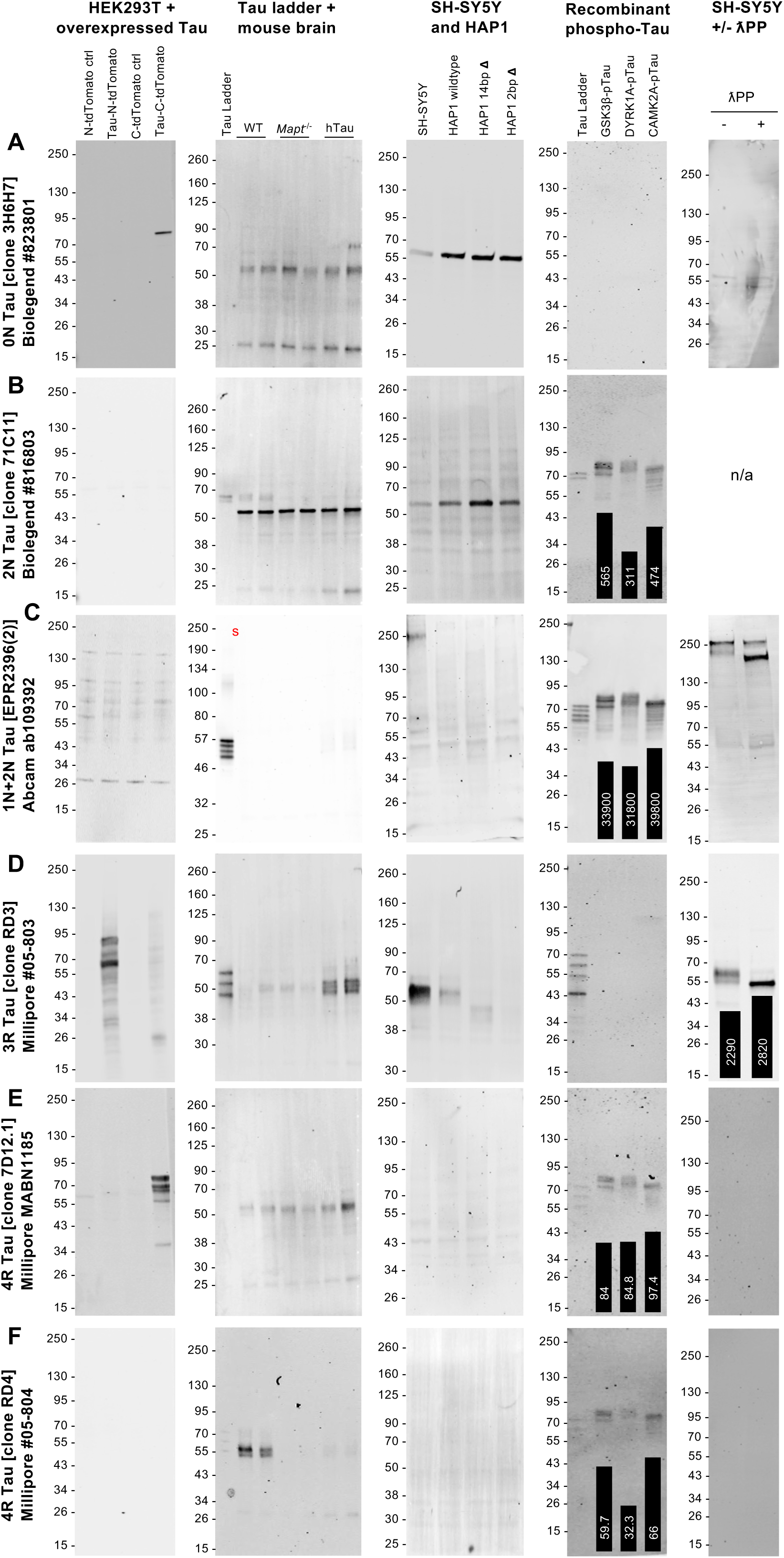
Validation of isoform-specific Tau antibodies by WB (part 1). **A-F:** WBs of lysates from HEK293T cells overexpressing 0N3R human Tau and corresponding control cells (**first column**); recombinant human Tau ladder (5 ng/isoform/lane), plus adult mouse brain lysates from wildtype, *Mapt^-/-^* and hTau mice (**second column**); lysates from SH-SY5Y neuroblastoma cells, plus HAP1 cells, parental (wildtype) and two cell lines carrying either a 14 bp deletion (14 bp Δ) or a 2-bp deletion (2 bp Δ) in *MAPT* exon 4 (**third column**); recombinant human Tau ladder (50 ng/isoform/lane) plus recombinant 2N4R Tau that has been phosphorylated by one of three known Tau kinases: GSK3ꞵ, DYRK1A or CAMKIIA (**fourth column**); lysates from SH-SY5Y neuroblastoma cells that have been either untreated (-) or treated (+) with λPP (**fifth column**). For WBs in the fourth and fifth columns, quantifications of the Tau signal intensity for each lane are shown superimposed on each WB image as a bar chart, with the respective value [a.u.] printed on or above each bar of the chart. WB membranes shown in each panel (**row**) were probed with a different Tau antibody: clone 3H6H7 (0N Tau) BioLegend #823801 (**A**), clone 71C11 (2N Tau) BioLegend #816803 (**B**), clone EPR2396(2) (1N and 2N Tau isoforms) Abcam #ab109392 (**C**), clone RD3 (3R Tau) Millipore #05-803 (**D**), clone 7D12.1 (4R Tau) Millipore #MABN1185 (**E**), clone RD4 (4R Tau) Millipore #05-804 (**F**).

In contrast, the 2N antibody reacted with two of the six isoforms of the recombinant Tau ladder, as expected, albeit weakly, and detected a faint but specific signal in wildtype mouse brain lysates (**Fig. 10B, second column**), in agreement with 2N Tau making up only a small proportion of the total Tau pool expressed in adult mouse brains [129]. This antibody did not detect specific bands in SH-SY5Y or HAP1 cells (**Fig. 10B, third column**), in line with published evidence that 2N Tau splice isoforms are not expressed in undifferentiated SH-SY5Y cells [89]. In both mouse brain and human cell lysates, however, the 2N antibody reacted with a prominent, non-specific band at ∼55 kDa. Our data, therefore, suggest that the 2N Tau antibody binds both murine and human Tau but reacts non-specifically with at least one other protein.

Despite this reagent being marketed as a total Tau antibody, we found that antibody Abcam #ab109392 (clone name: EPR2396(2)) detected only four of the six splice isoforms present in the recombinant Tau ladder (**Fig. 10C, second and fourth columns**). By using fluorescent WB detection to combine it with the Tau-12 antibody, we established that #ab109392 detects 1N and 2N, but not 0N, isoforms (**Supp. Fig. S5A**), suggesting that its epitope lies within the first N-terminal insert encoded by exon 2 (**Fig. 1**). In mouse brain samples, this antibody displayed a weak, but detectable, specific Tau signal in hTau mice (**Fig. 10C, second column; Supp. Fig. S3s**), supporting its specificity to human Tau, in agreement with the supplier’s product data sheet. No specific bands were detected in HEK293T cell lysates (**Fig. 10C, first column**) nor in SH-SY5Y or HAP1 cell lysates (**Fig. 10C, third column**), in line with HEK293T cells overexpressing 0N Tau and with previous literature that only 0N Tau isoforms are detectable in undifferentiated SH-SY5Y cells [89]. However, faint non-specific bands were detected in lysates of all human cell lines tested, offering an example of antibody specificity being dependent upon the origin of the samples being tested (**Fig. 10C, first and third columns**). We note that the same antibody clone, EPR2396(2), is marketed by other suppliers (e.g. Origene cat. no. TA307184 and GeneTex cat. no. 62576) as a phospho-Tau antibody that detects the protein when it is phosphorylated at Thr50, a residue encoded by *MAPT* exon 2. This is in line with our conclusion that its epitope lies within the first N-terminal insert. However, our findings that this antibody clone detects 1N and 2N isoforms of the recombinant Tau ladder demonstrates that its binding is not phosphorylation-dependent. In agreement with this conclusion, #ab109392 reacted equally with all three recombinant phosphorylated Tau versions (**Fig. 10C, fourth column**).

The RD3 antibody clone, raised against 3R Tau, detected 0N3R Tau when overexpressed in HEK293T cells (**Fig. 10D, first column**), reacted with its three target isoforms in the human Tau recombinant ladder (**Fig. 10D, second and fourth columns**), and immunoreacted with human Tau expressed in hTau brains (**Fig. 10D, second column**), as well as in SH-SY5Y and HAP1 cells (**Fig. 10D, third column**). No immunoreactivity was observed with wildtype murine Tau (**Fig. 10D, second column**), in agreement with published data that adult wildtype mouse brains express 4R Tau exclusively [129]. When tested on mouse brain lysates, RD3 detected very faint non-specific bands of ∼50 kDa, presumed to represent the heavy chain of endogenous mouse IgGs, although these did not interfere with Tau detection in hTau brain lysates (**Fig. 10D, second column**), whereas its signal appeared specific to Tau in human cell lysates (**Fig. 10D, third column**).

Of the 4R Tau antibodies tested, all four reacted with recombinant Tau, although the BioLegend and Millipore-7D12 reagents did so only weakly (**Figs. 10E-F, 11A-B, second and fourth columns**), suggesting that all of these antibodies can detect the target protein. However, Millipore-7D12 also reacted with 0N3R Tau overexpressed in HEK293T cells, indicating that it is not specific to 4R Tau (**Fig. 10E, first column**). Of the four 4R Tau antibodies, only the Millipore-RD4 and Abcam reagents detected Tau in mouse brain lysates, where both displayed high specificity based on comparison with *Mapt^-/-^* samples (**Figs. 10F, 11B, second column**). Although undifferentiated SH-SY5Y cells are often thought to express only the 0N3R isoform, expression of 4R Tau has also reported in these cells, albeit at much lower levels than 3R Tau [89,130,131]. In the case of HAP1 cells, our mass-spec analyses revealed the presence of 4R-specific Tau peptides in wildtype cells (**Supp. Fig. S2**, see peptide spanning the junction of exons 10-11). Despite this, only the Abcam reagent reacted with proteins present in SH-SY5Y and HAP1 human cell lysates (**Fig. 11B, third column**). However, the appearance of the band detected in HAP1 cells (sharp, rather than diffuse) and the fact that its apparent MW remained unchanged between wildtype and *MAPT*-edited cells calls its identity into question. In contrast, the two bands detected with the Abcam 4R Tau antibody in the 60-70 kDa MW range in SH-SY5Y lysates displayed the diffuse appearance typical of Tau bands observed in samples that have not been dephosphorylated (**Fig. 11B, third column**) and overlapped with the bands detected by the highly-specific total Tau, Tau-13 antibody (**Supp. Fig. S5B**). To complement the observations of previous studies, we obtained further evidence of low level 4R Tau expression in SH-SY5Y cells by combining RD3 with the highly specific #ab254256 total Tau antibody (**Supp. Fig. S5C**). Following dephosphorylation of SH-SY5Y lysates, #ab254256 detected two closely-spaced bands: (i) a prominent lower band that was also detected with the RD3 antibody, establishing this band’s identity as 3R Tau, and (ii) a weaker, upper band, that did not react with RD3 and whose MW is in agreement with that of the 0N4R Tau isoform (**Supp. Fig. S5B**; the two bands are indicated with red arrows). While the proximity of the putative 3R and 4R bands precludes precise quantifications, the relative difference in signal intensity between these two bands indicates that 4R isoforms account for <=10% of the total Tau signal detected in SH-SY5Y cells. Other specific total Tau antibodies, e.g. SP70 and Tau-12 (**Fig. 2D, F**), also detected a similar pattern of two closely-spaced bands in λPP-treated SH-SY5Y lysates. Combining RD3 with the Abcam 4R Tau antibody to probe λPP-treated SH-SY5Y lysates further demonstrated that the two antibodies detect closely-spaced but non-overlapping bands whose MW difference is in agreement with that of 0N3R and 0N4R Tau isoforms, respectively (**Supp. Fig. S5D**). These data suggest that undifferentiated SH-SY5Y cells express low levels of 4R Tau, which can be detected by the Abcam 4R Tau antibody. However, our data also suggest that the Abcam 4R Tau antibody may display non-specific cross-reactivity with proteins present at high levels in certain cell types, e.g. HAP1 cells. Taken together, our data indicate that, of the four antibodies that target 4R Tau tested here, only the Millipore-RD4 reagent detects both murine and human 4R Tau with high specificity, although the signal intensity obtained with this antibody was relatively faint.

**Figure 11.**
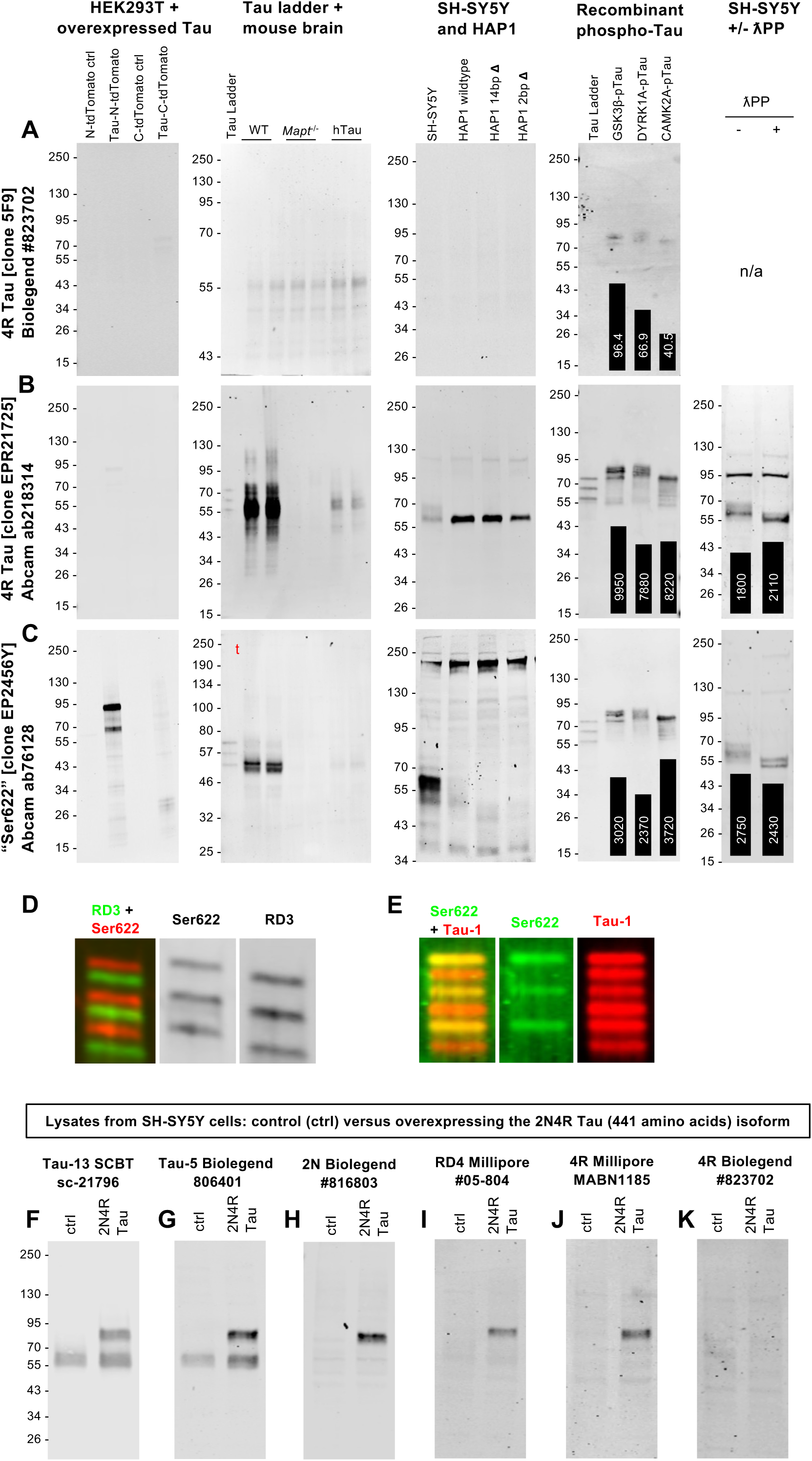
Validation of isoform-specific Tau antibodies by WB (part 2). **A-C:** WBs of lysates from HEK293T cells overexpressing 0N3R human Tau and corresponding control cells (**first column**); recombinant human Tau ladder (5 ng/isoform/lane), plus adult mouse brain lysates from wildtype, *Mapt^-/-^* and hTau mice (**second column**); lysates from SH-SY5Y neuroblastoma cells, plus HAP1 cells, parental (wildtype) and two cell lines carrying either a 14 bp deletion (14 bp Δ) or a 2-bp deletion (2 bp Δ) in *MAPT* exon 4 (**third column**); recombinant human Tau ladder (50 ng/isoform/lane) plus recombinant 2N4R Tau that has been phosphorylated by one of three known Tau kinases: GSK3ꞵ, DYRK1A or CAMKIIA (**fourth column**); lysates from SH-SY5Y neuroblastoma cells that have been either untreated (-) or treated (+) with λPP (**fifth column**). For WBs in the fourth and fifth columns, quantifications of the Tau signal intensity for each lane are shown superimposed on each WB image as a bar chart, with the respective value [a.u.] printed on or above each bar of the chart. WB membranes shown in each panel (**row**) were probed with a different Tau antibody: clone 5F9 (4R Tau) BioLegend #823702 (**A**), clone EPR21725 (4R Tau) Abcam #ab218314 (**B**), clone EP2456Y (“Ser622”) Abcam #ab76128 (**C**). **D, E**: WB of recombinant human Tau ladder was co-probed with the Ser622 antibody (red in **D**, green in **E**) and either the RD3 3R Tau antibody clone (**D**, green) or the Tau-1 antibody clone (**E**, red). **F-K**: WBs of lysates from SH-SY5Y cells overexpressing 2N4R human Tau or control (ctrl) cells were probed with different Tau antibodies: clone Tau-13 Santa Cruz Biotechnology #sc-21796 (**F**), clone Tau-5 BioLegend #806401 (**G**), clone 71C11 (2N Tau) BioLegend #816803 (**H**), clone RD4 Millipore #05-804 (**I**), clone 7D12.1 (4R Tau) Millipore #MABN1185 (**J**), clone 5F9 (4R Tau) BioLegend #823702 (**K**).

One antibody that warrants closer inspection is the Abcam #ab76128 reagent. This is marketed as an antibody that “detects Tau, whether phosphorylated or unphosphorylated on Ser622” (<u>link</u> to Abcam product page last checked on 5^th^ September 2022), hereafter referred to as the “Ser622 antibody”, and has been used in at least 21 published studies to date (<u>link</u> to Citeab page, last accessed 23^rd^ March 2023). Using residue numbering based on the 758 amino acids-long PNS-Tau isoform (Uniprot ID P10636-1, canonical Tau isoform on Uniprot) revealed that the Ser622 residue corresponds to Ser305 in the 2N4R Tau isoform, the last residue encoded by the alternatively-spliced exon 10, whose inclusion gives rise to 4R Tau, thus calling into question this reagent’s ability to detect all Tau splice isoforms. In line with this, our testing established that Ser622 detected the 4R, but not 3R, isoforms of the Tau ladder (**Fig. 11C-E**). Ser622 also detected Tau in wildtype mouse brain lysates with high specificity (**Fig. 11C, second column**). Similarly to the Abcam 4R Tau antibody, Ser622 immunoreacted with endogenous Tau in SH-SY5Y cell lysates (**Fig. 11C, third column**). Combining the RD3 and Ser622 antibodies to probe λPP-treated SH-SY5Y cell lysates established that the prominent Tau bands detected with each antibody do not overlap (**Supp. Fig. S5E**), although closer inspection shows that Ser622 also reacted, albeit weakly, with the RD3-immunoreactive band (**Supp. Fig. S5E**, red arrow), suggesting that Ser622 also reacts with 3R Tau. Confirming this possibility, Ser622 detected 0N3R Tau when overexpressed in HEK293T cells (**Fig. 11C, first column**). Taken together with the evidence that 4R isoforms represent a small fraction of the total Tau pool in SH-SY5Y cells, our data suggest that Ser622 has a much higher affinity for 4R Tau, but can also detect 3R isoforms, if the latter are present at high levels. Therefore, results obtained through using this antibody should be interpreted with caution.

Several of the isoform-specific antibodies tested did not react with Tau in any of the cell/tissue lysates (**Figs. 10-11**). In order to establish whether these can detect the target protein in complex biological samples, we overexpressed 2N4R Tau in SH-SY5Y cells. Total Tau antibodies Tau-13 and Tau-5 detected both the endogenous and overexpressed proteins, whereas the 2N, Millipore-RD4 and Millipore-7D12 detected only the latter (**Fig. 11F-J**). These data demonstrate that these three antibodies are able to detect their target protein when it is expressed at sufficiently high levels. In contrast, the BioLegend 4R antibody failed to detect 4R Tau even when the target protein was overexpressed (**Fig. 11K**).

Phosphorylatable residues are found within, and in proximity to, the regions targeted by all isoform-specific antibodies. Nevertheless, these antibodies are assumed to detect their target isoforms regardless of Tau phosphorylation status. By comparing antibody reactivity to recombinant Tau that has been phosphorylated by different kinases or to SH-SY5Y cell lysates prior or after λPP treatment, our data suggest that the binding of the following antibodies to Tau is partially hindered by phosphorylation: 2N, Millipore-RD4, BioLegend 4R and Ser622 (**Figs. 10-11, fourth and fifth columns**).

### Several Tau antibodies cross-react with MAP2 giving rise to mid-MW bands that may interfere with detection of some Tau isoforms

The C-terminal half of Tau shares extensive sequence homology with related microtubule-associated proteins (MAPs), in particular MAP2 and MAP4 [132]. This raises the possibility that antibodies whose epitopes lie in the C-terminal half of the protein, may cross-react with other MAPs. Indeed, the Tau-46 clone, whose epitope lies close to the Tau C-terminus, has been repeatedly proposed to cross-react with MAP2 [19,133–135], a feature that is also noted on the product data sheet by several commercial suppliers (e.g. Thermo Fisher Scientific, Santa Cruz Biotechnology, Cell Signalling Technology). In line with this, we observed that all three Tau-46 antibody clones tested, detected presumed non-specific bands: high MW bands >250 kDa in protein extracts from mouse brains and SH-SY5Y cells, and mid-MW bands (75-90 kDa MW range) in SH-SY5Y cell lysates (**Fig. 4D-F**). The MWs of these non-specific bands are in agreement with those reported for the MAP2a/MAP2b and MAP2c isoforms, respectively, while their absence from HAP1 cell lysates is in agreement with the characteristics of MAP2 as a neuronal-enriched protein. While cross-reactivity with the MAP2a/2b isoforms does not interfere with the detection of monomeric Tau on SDS-PAGE, owing to the large differences in MW, high-MW MAP2 isoforms might overlap with oligomeric/aggregated Tau species. More likely to be problematic, however, is the fact that mid-MW MAP2c isoforms migrate to an apparent MW similar to that of 2N Tau isoforms. As undifferentiated SH-SY5Y cells only express detectable levels of 0N Tau isoforms, this enables the clear distinction between common Tau and mid-MW MAP2c bands. However, caution should be exercised when analysing lysates from cells or tissues that express 2N Tau isoforms, as non-specific MAP2 signals might overlap the Tau bands.

Aligning the amino acid sequences of the 2N4R Tau and MAP2c isoforms, highlighted the extent of sequence similarity between the C-terminal halves of the two proteins (**Supp. Fig. S6A**) and raised the possibility that other C-terminal total Tau antibodies, as well as some phospho-Tau antibodies, may also cross-react with MAP2. Using fluorescent WB detection to combine pairs of Tau antibodies, we established that the K9JA, pSer199 and pSer202+pThr205 antibodies also react with mid-MW bands that overlap those detected by Tau-46 in SH-SY5Y cell lysates (**Supp. Fig. S7A-D**), raising the possibility that all of these antibodies cross-react with MAP2c. To test this possibility directly, we employed MAP2 antibodies, raised in either mouse or rabbit. As these were developed against an N-terminal fragment of MAP2, they do not cross-react with Tau, as evidenced by the fact that MAP2 antibodies did not detect the “canonical” Tau bands in the 55-70 kDa region in SH-SY5Y cell lysates (**Supp. Fig. S7E-N**), nor did they react with the recombinant Tau ladder (**Supp. Fig. S7O**). While the rabbit MAP2 antibody displayed non-specific cross-reactivity with a band of approximately 43 kDa (**Supp. Fig. S7E, M**; non-specific band is indicated with a red asterisk), this did not affect our analyses. Utilising fluorescent WB detection to combine MAP2 antibodies with different Tau antibodies, we found that the Tau-46, K9JA, A16097B, pSer199, pSer202+pThr205, and pSer396 Tau antibodies all reacted with mid-MW bands that were also immunoreactive with MAP2 antibodies (**Supp. Fig. S7E-J’**). Of note, sequence similarity raises the possibility that other phospho-Tau antibodies (e.g.: pThr205, pThr231, pSer262, pSer409, pSer422) may also cross-react with MAP2 (**Supp. Fig. S6A**). However, as most of these antibodies did not provide a positive signal in SH-SY5Y cell lysates, it was not possible to test this prediction experimentally. Another important consideration is that, whether phospho-Tau antibodies cross-react with MAP2 and, if so, which isoforms they detect, will depend on whether the respective MAP2 isoforms are phosphorylated at the corresponding residues in the samples being analysed.

The N-terminal halves of the Tau and MAP2 proteins are largely dissimilar (**Supp. Fig. S6A**), leading to the prediction that N-terminal antibodies (e.g. Tau-12, Tau-13, SP70, 43D, 5A6) and some mid-Tau antibodies (e.g. HT7 and Tau-5) should not cross-react with MAP2. Indeed, none of the N-term or mid-domain total Tau antibodies tested here detected non-specific bands in mouse brain lysates of MWs >250 kDa, where MAP2a and MAP2b isoforms are expected (**Figs. 2-3**). In line with our predictions, combining the Tau-13 or SP70 total Tau antibodies with MAP2 antibodies confirmed that neither detected MAP2-immunoreactive bands (**Supp. Fig. S7M, N**). Furthermore, most of the sequence encoded by the alternatively-spliced exon 10 and the first eight amino acids encoded by exon 11 are not present in MAP2 (**Supp. Fig. S6A**), indicating that 3R and 4R isoform-specific Tau antibodies are also unlikely to cross-react with MAP2. In agreement with this, we note that 3R and 4R Tau antibodies did not detect high-MW bands in mouse brain lysates nor mid-MW bands in SH-SY5Y lysates (**Figs. 10-11**). Finally, utilising fluorescent WB to combine the pSer198 or E178 phospho-Tau antibodies with MAP2 antibodies revealed that these two phospho-Tau antibodies also do not cross-react with MAP2 bands (**Supp. Fig. S7K-L**), a finding that is in agreement with predictions made based on alignment of the amino acid sequences (**Supp. Fig. S6A**). The differential results obtained with the pSer396 (**Supp. Fig. S7J, J’**) and E178 (**Supp. Fig. S7K, K’**) antibodies, both thought to target Tau phosphorylated at Ser396, suggest that differences may exist between the epitopes of these two antibodies.

To assess directly whether Tau antibodies cross-react with MAP2 on WB, we used recombinant MAP2c, carrying a ∼26 kDa GST tag and which, therefore, migrates on SDS-PAGE to an apparent MW of ∼100 kDa. This confirmed that clones K9JA, A16097B and Tau-46 all detected the purified, recombinant MAP2 protein on WB (**Supp. Fig. S7P**). In contrast and as expected, Tau-12 did not detect recombinant MAP2 (**Supp. Fig. S7P**). Taken together, these data demonstrate that several mid-domain and C-term total and phospho-Tau antibodies cross-react with MAP2, whereas antibodies whose epitopes lie in the N-terminal half of Tau as well as 3R- and 4R-targeting antibodies do not.

Tau also shares sequence similarities with MAP4, albeit to a lesser extent than with MAP2 [132]. Alignment of the Tau and MAP4 amino acid sequences revealed extensive similarities between the two proteins in the region encoded by Tau residues 251-358 (**Supp. Fig. S6B**). This suggested that, of the total and phospho-Tau antibodies tested, only the polyclonal K9JA total Tau antibody and the antibodies targeting pSer262 may potentially cross-react with MAP4. Comparison of the two protein sequences also revealed that, unlike MAP2, MAP4 shares with Tau most of the sequence encoded by the 4R-defining exon 10, including a stretch of 17 identical residues (**Supp. Fig. S6B**; Gln288-Gly304 in the 2N4R Tau sequence). This raises the possibility that some 4R isoform-specific Tau antibodies may cross-react with MAP4, depending on the exact location of their epitope. Indeed, in SH-SY5Y and HAP1 cell lysates, Ser622 cross-reacted strongly with a band just below the 250 kDa MW marker (**Fig. 11C**), a MW that does not correspond to any of the MAP2 isoforms, but which is consistent with the MW of the related MAP4 protein. Both MAP2 and MAP4 are expressed in SH-SY5Y cells at the mRNA [102] and protein levels [136]. Cross-reactivity of 3R Tau antibodies with MAP4, however, is not expected given that the MAP4 sequence contains a 38 amino acids-long, non-conserved stretch intercalated in between the regions of sequence homology with Tau exons 9 and 10 (**Supp. Fig. S6B**).

### High-MW Tau-immunoreactive bands are detected with several total Tau and phospho-Tau antibodies

Several total and phospho-Tau antibodies (e.g.: 5A6, Tau-12, Tau-13, Tau-5, K9JA, pSer198, pSer199, pThr231, pSer396, E178) detected high-MW Tau-immunoreactive bands of approximately 110-120 kDa in SH-SY5Y cell lysates (**Figs. 2, 3**, **6-8**). Detection with different antibodies that recognise a variety of epitopes distributed along the length of the protein – including the highly-specific antibodies Tau-12, Tau-13 and Tau-5, which target epitopes that are not shared with any other MAPs –, raised the possibility that these high-MW bands may represent “true” Tau signal. To test whether the different antibodies react with the same high-MW band, we used fluorescent WB detection to combine pairs of Tau antibodies raised in mouse and rabbit on the same membrane (**Supp. Fig. S8**). These experiments revealed overlap between the high-MW signals detected with the different antibodies (**Supp. Fig. S8**). Taken together, these data offer evidence that the high-MW Tau-immunoreactive bands detected with different antibodies may indeed represent specific Tau signals, although further work is needed to establish their exact identity.

### Analogous approaches to validating Tau antibodies by IHC-IF

To test Tau antibody performance in IHC, FFPE tissue sections were chosen in preference to frozen samples as most tissues available for research are routinely processed by formalin fixation. Since proteins are denatured during most WB protocols, this may expose antibody binding sites that are not accessible *in situ* and are not, therefore, available for detection via IHC. Conversely, antibody binding sites present *in situ* may be destroyed when denaturing proteins for WB. For FFPE samples, target binding sites are further modified through the formalin fixation and antigen retrieval steps. Therefore, determination of antibody performance and specificity by WB does not guarantee suitability for IHC applications. To determine their ability to bind Tau, we first tested antibodies by IHC-IF using FFPE mouse brain sections from 9-month old rTg4510 mice. This mouse model overexpresses the human Tau 0N4R isoform, carrying the P301L pathological mutation, under the control of the calcium calmodulin kinase II (CaMKII) promoter system, resulting in high expression in the hippocampus and neocortex, and the subsequent accumulation of hyperphosphorylated Tau and NFTs in these regions [76]. To assess the effect of phosphorylation on antibody binding and, in the case of phospho-specific antibodies, to establish the requirement for protein phosphorylation, we compared the stainings obtained with each antibody in control, untreated tissue sections versus λPP-treated serial sections (**Figs. 12, 14** and **16**).

**Figure 12.**
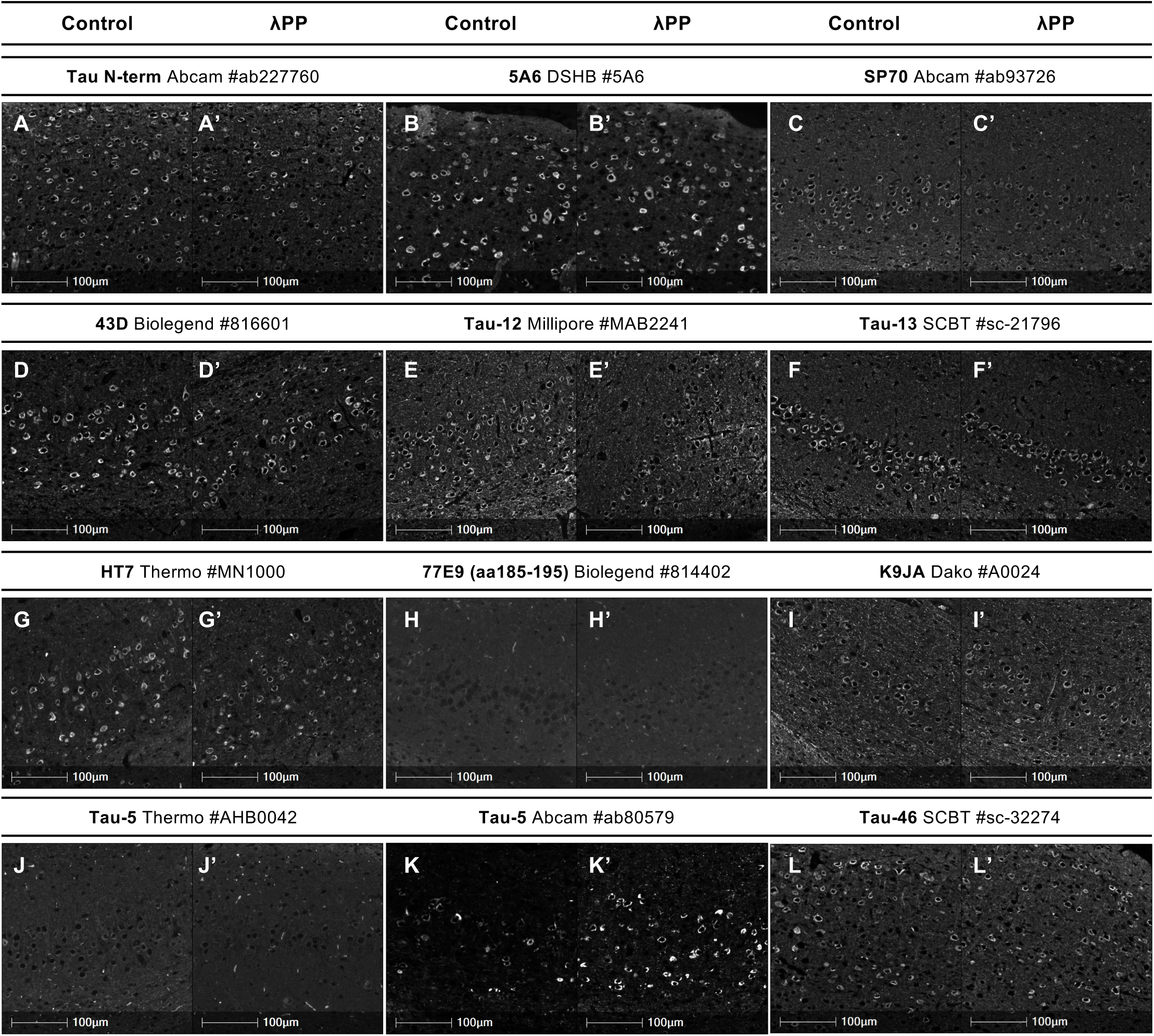
Validation of total Tau antibodies by IHC-IF (part 1). **A-L’:** Fluorescence micrographs of serial FFPE brain sections from 9-month old rTg4510 mice, either untreated (control; **A, B, C, D, E, F, G, H, I, J, K and L**) or treated (λPP; **A’, B’, C’, D’, E’, F’, G’, H’, I’, J’, K’ and L’**) with λPP, immunostained with Tau antibodies: N-terminal-targeting antibody Abcam #227760 (**A, A’**), clone 5A6 DSHB (**B, B’**), clone SP70 Abcam #ab93726 (**C, C’**), clone 43D BioLegend #816601 (**D, D’**), clone Tau-12 Millipore #MAB2241 (**E, E’**), clone Tau-13 Santa Cruz Biotechnology #sc-21796 (**F, F’**), clone HT7 Thermo Fisher Scientific #MN1000 (**G, G’**), clone 77E9 BioLegend #814402 (**H, H’**), clone K9JA Dako #A0024 (I, I’), clone Tau-5 Thermo Fisher Scientific #AHB0042 (**J, J’**), clone Tau-5 Abcam #ab80579 (**K, K’**), clone Tau-46 Santa Cruz Biotechnology #sc-32274 (**L, L’**). Brain region: cortex. Tau staining is shown in grayscale. Scale bars = 100µm.

Most studies published to date have employed overexpression models that express transgenic Tau at levels much higher than endogenous Tau. However, an antibody’s potential non-specific cross-reactivity with other proteins is difficult to assess in these samples owing to the disproportionately high levels of expression of the target protein compared to any potential sources of non-specific signal. To characterise antibody performance when Tau is expressed at physiological levels, we used FFPE brain tissue sections from wildtype, *Mapt*^-/-^ and hTau mice, as described above. As the predominant Tau isoform present in wildtype mice is 4R [129], whereas in the hTau mice it is 3R [74], this allows for additional characterisation of isoform-specific antibodies. Antibodies tested at this stage represented a selected subset of those reagents that provided a positive signal in rTg4510 brain sections. Note that different humanised transgenic mouse models are referred to as hTau or similar in the literature. Some of these overexpress the human Tau protein and are expected to react more strongly with Tau antibodies than the hTau mouse samples used in this study, where Tau expression is driven solely by the human *MAPT* promoter sequences captured within the 9 kb upstream of the *MAPT* transcription start site [75].

Micrographs of control brain sections that were probed with secondary antibodies only, omitting the primary antibody, confirmed absence of signal following the computational removal of autofluorescence, thus demonstrating that any immunofluorescent signal detected in the final images resulted from binding of the primary antibodies (**Supp. Fig. S9B**).

### Validation of PTM-independent “total” Tau antibodies by IHC-IF

Of the 12 total Tau antibodies tested (N-term #ab227760, 5A6, SP70, 43D, Tau-12, Tau-13, HT7, K9JA, 2 x Tau-5, 77E9 and Tau-46), all detected a positive signal in rTg4510 mouse brain sections, except for clone 77E9 (**Fig. 12**), thus demonstrating their ability to detect human Tau in FFPE tissue sections when the target protein is present at high levels. Similar to our WB observations, Tau-5 purchased from two different manufacturers performed differently: immunostaining with the Thermo Fisher Scientific reagent resulted in a weak signal that was hardly detectable above background, when used at the manufacturer’s recommended concentration (1:200) (**Fig. 12J-J’**). By contrast, staining with the Abcam reagent provided a clear signal against a low background (**Fig. 12K-K’**). Dephosphorylation of proteins in tissue sections by treatment with λPP did not noticeably alter the pattern or intensity of staining obtained with most total Tau antibodies, except for the Tau-5 clone purchased from Abcam (**Fig. 12**). For this antibody, increased staining intensity was observed following λPP treatment (**Fig. 12K-K’, Supp. Fig. S10A, C**), suggesting that its binding to Tau in FFPE tissue sections is partially inhibited by the presence of phosphorylated residues. In contrast, λPP treatment did not affect the staining intensity or percentage of Tau-positive cells in sections probed with the Tau-13 antibody (**Supp. Fig. S10B, D**). These findings are in agreement with our WB observations reported above.

Upon testing of mouse brain sections from wildtype, *Mapt^-/-^* and hTau mice, we found that the immunofluorescence signal for Tau was weak in wildtype as well as in hTau mouse brains, both of which express Tau at physiological levels (**Fig. 13**). Relatively few studies have performed Tau immunostainings on wildtype mouse brains, but evidence from the literature supports weak immunoreactivity of endogenous murine Tau [39,137–142]. Human Tau-specific antibodies SP70, Tau-12 and Tau-13 did not detect a signal in wildtype or *Mapt^-/-^* mouse brains but did so in hTau mice, demonstrating their specificity for human Tau (**Fig. 13A-C”**). No clear signal was detected with the human-specific HT7 antibody in either wildtype, hTau or *Mapt^-/-^* mice, suggesting that, while this antibody is specific, it may not be suitable for detecting low levels of Tau expression (**Fig. 13D-D”**). K9JA displayed specific Tau immunoreactivity in both wildtype and, more weakly, in hTau mice (**Fig. 13E-E”**), while Tau-5 detected Tau in wildtype mouse brain sections, albeit weakly, but failed to produce a clear signal in hTau brains, suggesting it is better suited for the detection of murine Tau (**Fig. 13F-F”**). Unexpectedly, Tau-46 immunostaining was observed primarily in the apical dendrites of hippocampal pyramidal neurons, but this was present in mouse brain sections from all three genotypes, indicating non-specific cross-reactivity (**Fig. 13G-G”**). We note that this staining pattern is reminiscent of that obtained in wildtype mouse brain sections with anti-MAP2 antibodies [39]. Based on knowledge of its epitope, together with our WB data above (**Supp. Figs. S6-S7**), we propose that the Tau-46 immunoreactivity observed in wildtype, *Mapt^-/-^* and hTau mouse brain sections is a result of the antibody’s cross-reactivity with MAP2, suggesting that even in wildtype brains, the endogenous Tau signal detected with this antibody is weak by comparison. These findings are largely in agreement with our WB data presented above. One notable difference between our WB and IHC-FFPE observations is that K9JA, which displays non-specific cross-reactivity in WB applications (**Fig. 3F; Supp. Fig. S3e**), appears to detect Tau with high specificity in IHC-FFPE staining (**Fig. 13E-E”**), suggesting that K9JA cross-reacts with additional epitopes that are exposed by denaturing proteins during SDS-PAGE sample preparation.

**Figure 13.**
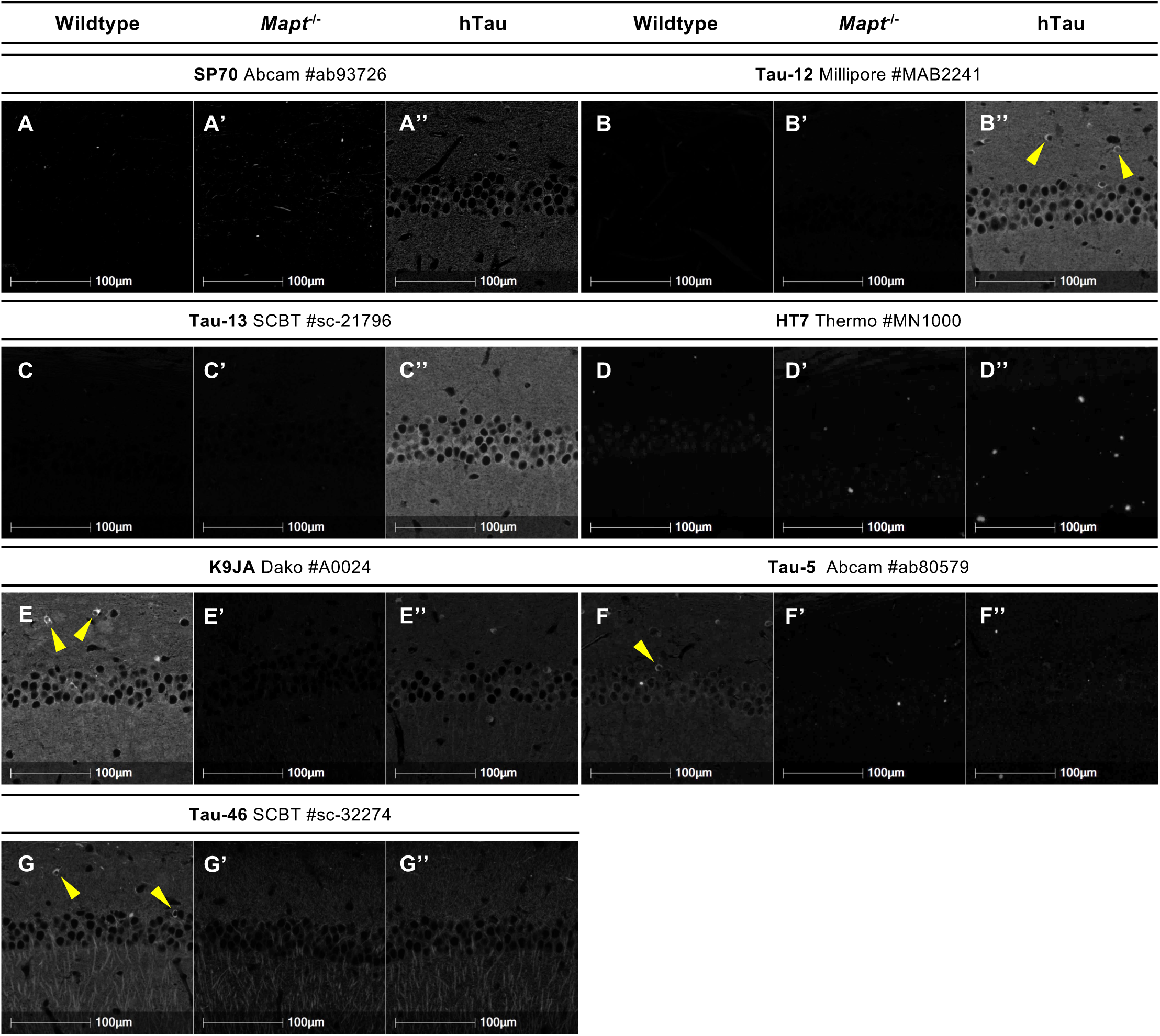
Validation of total Tau antibodies by IHC-IF (part 2). **A-G”:** Fluorescence micrographs of FFPE brain sections from 5-month old wildtype (**A, B, C, D, E, F and G**), *Mapt^-/-^*(**A’, B’, C’, D’, E’, F’ and G’**) and hTau (**A’’, B’’, C’’, D’’, E’’, F’’ and G’’**) mice immunostained with Tau antibodies: SP70 Abcam #ab93726 (**A-A’’**), clone Tau-12 Millipore #MAB2241 (**B-B’’**), clone Tau-13 Santa Cruz Biotechnology #sc-21796 (**C-C’’**), clone HT7 Thermo Fisher Scientific #MN1000 (**D-D’’**), clone K9JA Dako #A0024 (**E-E’’**), clone Tau-5 Abcam #ab80579 (**F-F’’**), clone Tau-46 Santa Cruz Biotechnology #sc-32274 (**G-G’’**). Yellow arrowheads indicate Tau-positive cell bodies located in *stratum oriens*. Brain region: CA1 region of the hippocampus. Tau staining is shown in grayscale. Scale bars = 100µm.

In wildtype and/or hTau mouse brain sections, a small number of cells located in the *stratum oriens* immunoreacted strongly with the Tau-12, K9JA, Tau-5 and Tau-46 total Tau antibodies (**Fig. 13B’, E, F, G**; indicated with yellow arrowheads). Specificity of this signal was demonstrated by its absence in brain sections from *Mapt^-/-^* mice (**Fig. 13B’, E’, F’, G’**). These data suggest that Tau is expressed in a subset of *stratum oriens* interneurons in adult mice.

### Validation of phosphorylation-dependent Tau antibodies by IHC-IF

As a consequence of the overexpression of human tau carrying the P301L mutation, rTg4510 mice display accumulation of hyperphosphorylated Tau and develop robust NFT-like pathology at 4-5 months of age [76]. Many of the phosphorylation-dependent antibodies as well as the oligomeric-specific antibodies are, therefore, predicted to provide a positive signal in rTg4510 mouse brain sections. In line with this, all phospho-Tau antibodies tested (pThr181, pSer198, pSer199, pSer202+pTh205, AT8, AT100, pSer214, pThr231, AT180, pSer238, pSer262, pSer396, PHF-13, E178, pSer404 and pSer409) yielded a clear positive immunoreactivity in brain sections from rTg4510 mice (Fig. 14), demonstrating that these antibodies have the ability to react with phosphorylated Tau in FFPE tissue sections when the target epitope is present at high levels.

**Figure 14.**
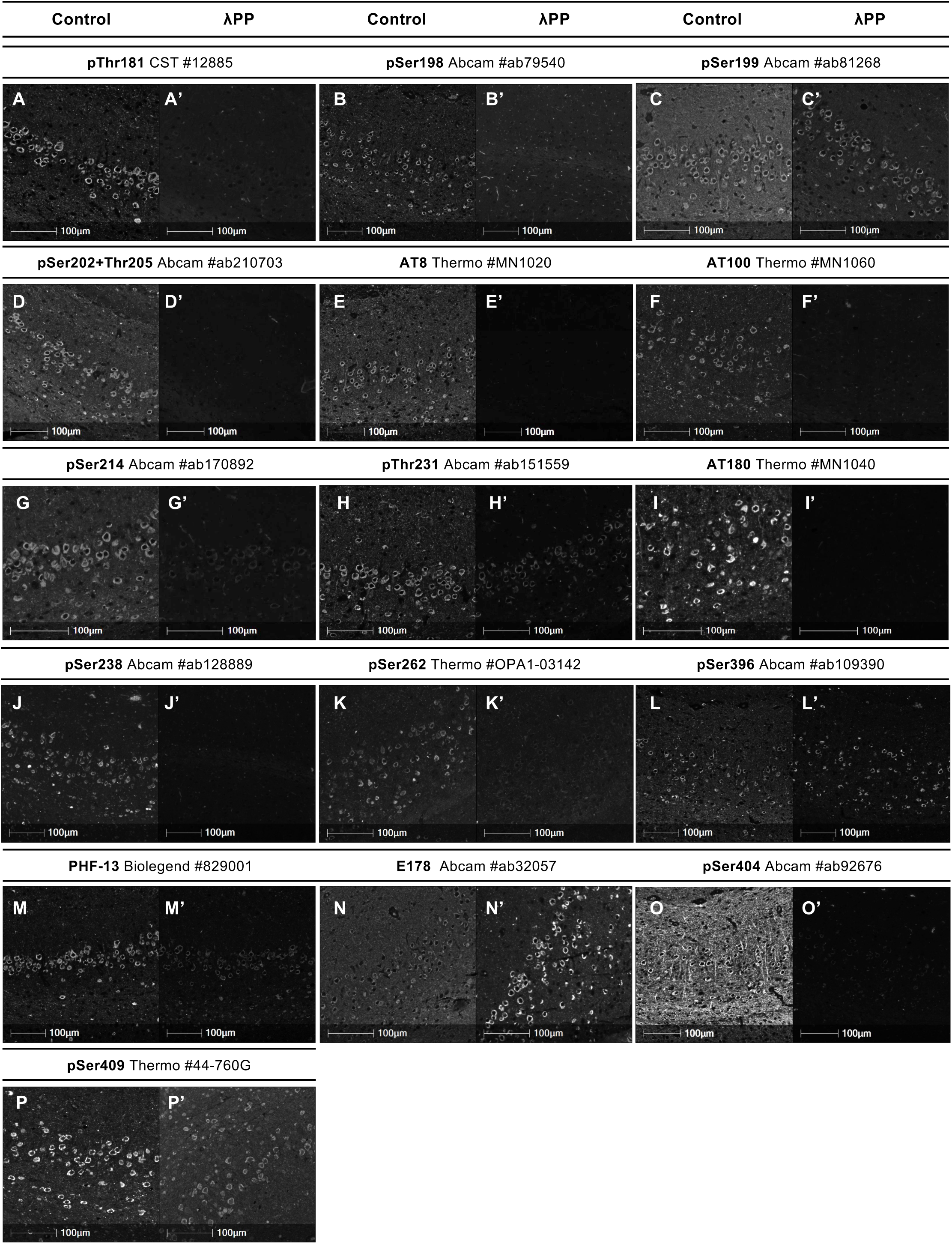
Validation of phospho-Tau antibodies by IHC-IF (part 1). **A-P’:** Fluorescence micrographs of serial FFPE brain sections from 9-month old rTg4510 mice, either untreated (control; **A, B, C, D, E, F, G, H, I, J, K, L, M, N, O and P**) or treated (λPP; **A’, B’, C’, D’, E’, F’, G’, H’, I’, J’, K’, L’, M’, N’, O’ and P’**) with λPP, immunostained with Tau antibodies: pThr181 Cell Signalling Technologies #12885 (**A, A’**), pSer198 Abcam #ab79540 (**B, B’**), pSer199 Abcam #81268 (**C, C’**), pSer202+pThr205 Abcam #210703 (**D, D’**), AT8 Thermo Fisher Scientific #MN1020 (**E, E’**), AT100 Thermo Fisher Scientific #MN1060 (**F, F’**), pSer214 Abcam #ab170892 (**G, G’**), pThr231 Abcam #ab151559 (**H, H’**), AT180 Thermo Fisher Scientific #MN1040 (**I, I’**), pSer238 Abcam #ab128889 (**J, J’**), pSer262 Thermo Fisher Scientific #OPA1-03142 (**K, K’**), pSer396 Abcam #ab109390 (**L, L’**), clone PHF-13 BioLegend #829001 (**M, M’**), clone E178 Abcam #ab32057 (**N, N’**), pSer404 Abcam #ab92676 (**O, O’**), Ser409 Thermo Fisher Scientific #44-760G (**P, P’**). Brain region: cortex. Tau staining is shown in grayscale. Scale bars = 100 µm.

Following treatment of the samples with λPP, immunoreactivity of the pThr181, pSer198, pSer202+pTh205, AT8, AT100, AT180, pSer238 and pSer404 antibodies was abrogated, demonstrating the phosphorylation-dependence of their epitopes (**Fig. 14**). For pSer214, pThr231, pSer262, PHF-13 and pSer409, λPP treatment resulted in decreased immunoreactivity but the signal was not completely abrogated (**Fig. 14**), supporting the phosphorylation-dependence of these epitopes while also implying that several of the phospho-epitopes present in rTg4510 mouse brains were incompletely dephosphorylated under the conditions employed here. Unexpectedly, λPP treatment did not alter the intensity of the immunoreactive signal observed in the cell bodies of rTg4510 cortical neurons with the pSer199 antibody (**Fig. 14C-C’**). This contrasts with our WB data in which pSer199 immunoreactivity was abrogated by dephosphorylation with λPP of SH-SY5Y cell lysates, which do not contain Tau aggregates (**Fig. 6C, fifth column**). Furthermore, in the case of antibodies pSer396 and E178, which are thought to detect Tau phosphorylated at Ser396, staining intensity increased in λPP-treated rTg4510 brain sections compared to control sections (**Fig. 14L-L’, N-N’**). Close inspection of the images revealed two types of signal in untreated/control sections: a weak, diffuse signal was observed in a subset of cell bodies as well as projections, while intense staining was present in the majority of cortical neuron cell bodies (**Fig. 14C-C’, N-N’**). The latter appeared fibrillar, reminiscent of Tau bundles/PHF aggregates. In the case of the pSer199, pSer396 and E178 antibodies, λPP treatment abrogated the weak, diffuse staining but not the intense/fibrillar staining, suggesting that the respective phospho-epitopes are readily dephosphorylated only when Tau is present in non-aggregated/prefibrillar states. We therefore hypothesised that these phospho-epitopes lie in regions of the protein that are not accessible to phosphatases when Tau is present in PHF aggregates, thus rendering them resistant to λPP treatment. Co-staining with Thioflavin S (ThS), a fluorescent dye with ꞵ-sheet binding properties, confirmed that the intense phospho-Tau staining observed in rTg4510 cortical neurons co-localised with ThS-positive protein aggregates (**Supp. Fig. S11**). While λPP has been used successfully *in vitro* to dephosphorylate PHF-Tau extracted from AD human brains [143], to the best of our knowledge, it has not previously been used to dephosphorylate Tau *in situ* and, thus, its efficiency towards PHF-Tau present in FFPE tissue sections is yet to be characterised. Staining FFPE human brain sections from control individuals and patients with AD, Pick’s Disease (PiD) or globular glial tauopathy (GGT) confirmed a lack of detectable signal with the AT8 and E178 antibodies in control brain sections, while positive immunostaining was observed with both antibodies in AD, PiD and GGT brains (**Supp. Fig. S12A**). Similarly to our observations in rTg4510 mouse brain sections, treatment with λPP abrogated the AT8 staining observed in AD, PiD and GGT brains, but increased the staining intensity obtained with E178 in AD and PiD brains, while E178 staining in GGT brains was unaffected (**Supp. Fig. S12A**). Co-staining with ThS confirmed that AT8 and E178 signals co-localised with ThS-positive protein aggregates present in human AD brains, which remained true for the enhanced E178 immunostaining observed in λPP-treated brain sections (**Supp. Fig. S12B**). Taken together with the fact that neither antibody immunoreacted with proteins present in control human brain (**Supp. Fig. S12A**), this suggests that both antibodies detect pathological PHF-Tau. Based on the observations presented here, we hypothesise that, when Tau is aggregated in PHFs, phosphorylation at the pSer396/E178 epitope is resistant to λPP treatment. We further hypothesise that binding of the E178 and pSer396 antibodies can be inhibited by phosphorylation at a nearby site, thus explaining the improved signal seen when this inhibitory phosphorylation event is removed by λPP treatment.

Ten (pThr181, pSer198, pSer199, AT8, AT100, pSer214, AT180, pSer396, E178 and pSer404) of the 16 phospho-antibodies tested in rTg4510 mouse brains were further tested in brain sections from wildtype, *Mapt*^-/-^ and hTau mice to assess specificity (**Fig. 15**). No specific immunoreactivity was detected with antibodies pThr181, AT8, AT100, pSer396 and E178, in either wildtype or hTau brains, in agreement with the much lower phosphorylation levels expected for wildtype Tau expressed at endogenous levels (**Fig. 15**). Lack of AT8 immunoreactivity in wildtype rodent brains is in line with published literature [19,144]. For all these antibodies, lack of immunoreactivity in *Mapt^-/-^* brains, taken together with IHC data from rTg4510 brains, demonstrate their specificity for phosphorylated Tau. Weak but detectable immunoreactivity was observed for antibodies pSer214, AT180 and pSer404 in wildtype and hTau brains, with no signal detected in *Mapt*^-/-^ brains, confirming the specificity of these reagents (**Fig. 15**). In addition, a small number of positive cells were also detected in the *stratum oriens* of wildtype, but not *Mapt*^-/-^, mouse hippocampi with all three antibodies (**Fig. 15F, G, J**; yellow arrowheads). These data suggest that Tau phosphorylated at the pSer214, AT180 and pSer404 epitopes is present in *stratum oriens* interneurons in wildtype adult mice. Finally, clear immunoreactive signals were detected in the soma of CA1 pyramidal neurons with the pSer198 and pSer199 antibodies in both wildtype and hTau mouse hippocampi (**Fig. 15B, B”, C, C”**). Specificity of the cytoplasmic staining observed with these two antibodies was confirmed by comparison with *Mapt*^-/-^ brains (**Fig. 15B’, C’**). In addition, pSer199 also displayed weak nuclear immunoreactivity in wildtype mouse CA1 pyramidal neurons. However, the intensity of this signal increased in *Mapt*^-/-^ and hTau brains, suggesting that pSer199 cross-reacts with a non-specific nuclear phospho-epitope, the levels of which increase in neurons that lack mouse Tau (**Fig. 15C-C”**). As our WB data suggested that pSer199 cross-reacts with phosphorylated MAP2 (**Supp. Fig. S7**) and given that MAP2 levels have been reported to increase in *Mapt*^-/-^ mouse brains [145,146] and that some MAP2 isoforms can localise to the nucleus [147,148], one possibility is that the nuclear pSer199 signal in *Mapt^-/-^* and hTau pyramidal neurons represents phosphorylated MAP2. Due to the potential large difference in abundance between cytoplasmic and nuclear MAP2 pools, the latter may not be easily detectable by IHC when probing with total MAP2 antibodies. To test the hypothesis that this difference results from an increase in nuclear MAP2 phosphorylation in *Mapt^-/-^* brains, assessment of phospho-MAP2 with antibodies targeting specific phosphorylation events will be required.

**Figure 15.**
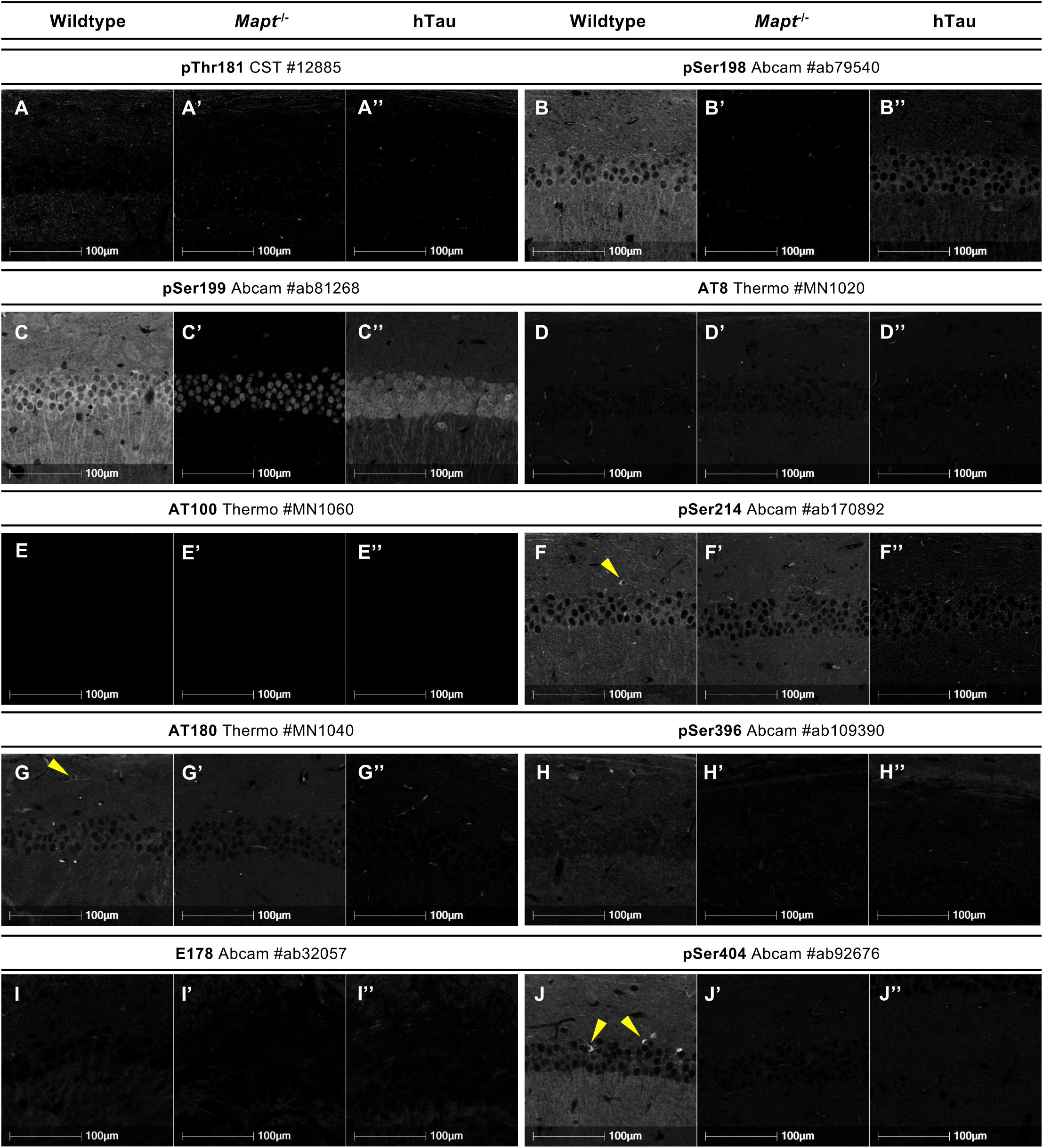
Validation of phospho-Tau antibodies by IHC-IF (part 2). **A-J”:** Fluorescence micrographs of FFPE brain sections from 5-month old wildtype (**A, B, C, D, E, F, G, H, I and J**), *Mapt^-/-^* (**A’, B’, C’, D’, E’, F’, G’, H’, I’ and J’**) and hTau (**A’’, B’’, C’’, D’’, E’’, F’’, G’’, H’’, I’’ and J’’**) mice immunostained with Tau antibodies: pThr181 Cell Signalling Technologies #12885 (**A-A’’**), pSer198 Abcam #ab79540 (**B-B’’**), pSer199 Abcam #81268 (C-C’’), AT8 Thermo Fisher Scientific #MN1020 (**D-D’’**), AT100 Thermo Fisher Scientific #MN1060 (**E-E’’**), pSer214 Abcam #ab170892 (**F-F’’**), AT180 Thermo Fisher Scientific #MN1040 (**G-G’’**), pSer396 Abcam #ab109390 (**H-H’’**), clone E178 Abcam #ab32057 (**I-I’’**), pSer404 Abcam #ab92676 (**J-J’’**). Yellow arrowheads indicate Tau-positive cell bodies located in *stratum oriens*. Brain region: CA1 region of the hippocampus. Tau staining is shown in grayscale. Scale bars = 100 µm.

### Validation of other PTM-dependent Tau antibodies by IHC-IF

Given the accumulation of phosphorylated Tau and overt NFT-like pathology in rTg4510 mouse brains by 9 months (the age of animals employed in this study), the Tau-1 clone was expected to provide weak signal prior to dephosphorylation. Conversely, epitopes of the antibody targeting caspase-cleaved D421 Tau, an event linked with Tau pathology, were anticipated to be present at high levels. As predicted, Tau-1 provided no detectable signal in neuronal cell bodies in samples that had not been dephosphorylated (**Fig. 16A**). This is consistent with the conclusion that Tau is mainly hyperphosphorylated in these tissues. Phosphatase treatment enhanced Tau-1 immunoreactivity (**Fig. 16A’**). A weak Tau-1 signal was detected in wildtype and hTau brains, which was abrogated in *Mapt^-/-^* brains, demonstrating the specificity of the Tau-1 antibody (**Fig. 16H-H”**). Staining rTg4510 mouse brain sections with the D421 antibody produced only weak immunoreactivity (**Fig. 16B**), but the IF signal intensity doubled following λPP treatment (**Fig. 16B’**, **Supp. Fig. S13**). Taken together with our WB findings (**Fig. 9D**), these data suggest that phosphorylation has an inhibitory effect on the binding of the D421 antibody clone to Tau. However, given our observations that D421 also detects full-length Tau by WB (**Fig. 9D**; fourth column), we cannot determine whether D421-immunoreactivity in rTg4510 mouse brain sections indicates the presence of Tau cleaved at Asp421 or whether it is a consequence of the high levels of Tau expression in these samples.

**Figure 16.**
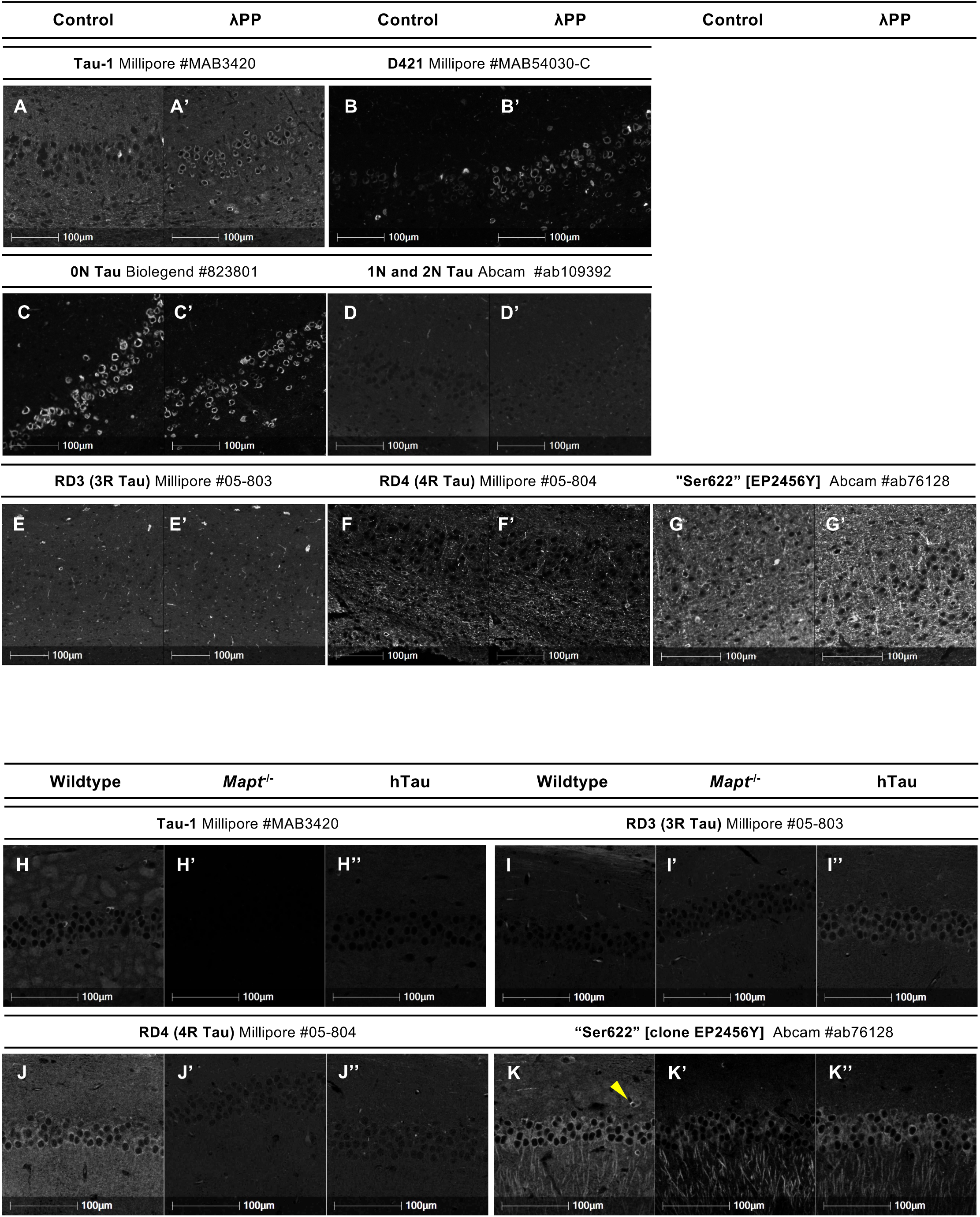
Validation of isoform-specific Tau antibodies and other PTM-dependent Tau antibodies by IHC. **A-G’:** Fluorescence micrographs of serial FFPE brain sections from 9-month old rTg4510 mice, either untreated (control; **A, B, C, D, E, F and G**) or treated (λPP; **A’, B’, C’, D’, E’, F’ and G’**) with λPP, immunostained with Tau antibodies: 0N Tau [clone 3H6H7] BioLegend #823801 (**A, A’**), 1N and 2N Tau [clone EPR2396(2)] Abcam #ab109392 (**B, B’**), clone RD3 (3R Tau) Millipore #05-803 (**C, C’**), clone RD4 (4R Tau) Millipore #05-804 (D, D’), “Ser622” [clone EP2456Y] Abcam #ab76128 (**E, E’**), clone Tau-1 Millipore #MAB3420 (**F, F’**), clone D421 Millipore #MAB54030-C (**G, G’**). Brain region: cortex. Tau staining is shown in grayscale. Scale bars = 100 µm. **H-K”:** Fluorescence micrographs of FFPE brain sections from 5-month old wildtype (**H, I, J and K**), *Mapt^-/-^* (**H’, I’, J’ and K’**) and hTau (**H’’, I’’, J’’ and K’’**) mice immunostained with Tau antibodies: clone RD3 (3R Tau) Millipore #05-803 (**H-H’’**), clone RD4 (4R Tau) Millipore #05-804 (**I-I’’**), “Ser622” [clone EP2456Y] Abcam #ab76128 (**J-J’’**), clone Tau-1 Millipore #MAB3420 (**K-K’’**). Yellow arrowheads indicate Tau-positive cell bodies located in *stratum oriens*. Brain region imaged: CA1 region of the hippocampus. Tau staining is shown in grayscale. Scale bars = 100 µm.

### Validation of isoform-specific Tau antibodies by IHC-IF

As rTg4510 mice overexpress the 0N4R human Tau isoform and 0N4R is also the predominant endogenous Tau isoform expressed in adult mouse brains [129], only 0N and 4R antibodies were expected to provide a positive signal in brain sections from these animals. In line with this, no signal was detected with the 1N+2N Tau (#ab109392) antibody nor with the RD3 3R Tau antibody, supporting these antibodies’ specificity (**Fig. 16D-D’, E-E’**). In the rTg4510 cortex, Tau-overexpressing cells immunoreacted strongly with the 0N antibody (**Fig. 16C-C’**) but, unexpectedly, not with the Millipore-RD4 4R Tau antibody (**Fig. 16F-F’**). However, the latter immunostained neuronal projections and a small number of cell bodies in the rTg4510 cortex (**Fig. 16F-F’**). Given that the MTBRs are central to PHF formation [149–152] and that the majority of Tau-overexpressing cells in rTg4510 brains contain ThS-positive PHFs (**Supp. Fig. S11**) [153–155], we hypothesise that the pattern of staining obtained with RD4 in these samples implies the fact that its epitope is inaccessible in aggregated Tau. If so, RD4 immunoreactivity in cortical cell bodies and neuronal projections would be indicative of the presence of 4R Tau in unaggregated/pre-tangle states. The staining patterns and signal intensities obtained with the 0N and RD4 antibodies were not noticeably altered by λPP treatment, in agreement with their status as PTM-independent antibodies (**Fig. 16C’, F’**). In line with conclusions from our WB data that Ser622 detects 4R Tau with much higher affinity than 3R Tau (**Fig. 11C-E**; **Supp. Fig. S5E**), staining rTg4510 mouse brain sections with the Ser622 antibody revealed an immunoreactivity pattern similar to that obtained with the RD4 antibody, with the highest immunoreactivity observed in neuronal projections (**Fig. 16G-G’**).

The RD3 and RD4 antibodies were tested further by using wildtype, hTau and *Mapt^-/-^* brain sections. In contrast to adult wildtype mouse brains, brains of hTau animals express both 3R and 4R Tau isoforms, with 3R Tau being predominant [156]. In line with this, RD3 immunoreactivity was observed in hTau mice, but not in wildtype or *Mapt^-/-^* (**Fig. 16I-I”**), while RD4 immunoreacted with hippocampal neurons of wildtype but not hTau or *Mapt^-/-^* mice (**Fig. 16J-J”**). These data support the specificity of the RD3 and RD4 antibodies to 3R and 4R Tau isoforms, respectively. Probing of wildtype, *Mapt^-/-^* and hTau mouse brain sections with the Ser622 antibody, resulted in striking staining of the cell bodies and apical dendrites of CA1 pyramidal neurons (**Fig. 16K-K”**). Although the Ser622-immunoreactive signal observed in the soma of pyramidal neurons diminished in *Mapt^-/-^* brains, immunoreactivity in the cells’ apical dendrites was maintained in all three genotypes, demonstrating lack of specificity to Tau (**Fig. 16K-K”**). Indeed, this staining pattern is reminiscent of the staining observed with anti-MAP2 antibodies [39]. As the sequence surrounding the Ser622 epitope is not present in MAP2, but the Tau sequence in this region is highly similar to that of MAP4 (**Supp. Fig. S6B**) and our WB data revealed that Ser622 cross-reacts with a protein of a MW similar to that of MAP4 (**Fig. 11C, third column**), it seems likely that the Ser622 antibody cross-reacts with MAP4. Although MAP4 was originally described as non-neuronal [132], its expression was later reported in all regions of the mouse brain, including in CA1 pyramidal neurons, where MAP4 localises to the cell bodies and apical dendrites [157]. While it is not possible to conclude with certainty whether Ser622 cross-reacts with MAP4 or another protein, our data demonstrate that this antibody detects Tau, when the target protein is present at sufficiently high levels, but also binds non-specifically to other neuronal proteins (possibly MAP4).

## Discussion

Here we have used WB and IHC-IF approaches to test and validate a broad panel of commercially-available total, PTM-dependent and isoform-specific Tau antibodies. A total of 53 antibodies were validated for use in WB and a subset of these, 35 antibodies, were also validated for use in IHC-IF on FFPE tissue sections. As a result, we provide a comprehensive analysis of antibodies suitable for the immunodetection of murine and/or human Tau via WB, IHC-IF or both. The overall performance in WB and IHC-FFPE applications of the reagents tested in this study is summarised in Figs. 17 and 18.

**Figure 17.**
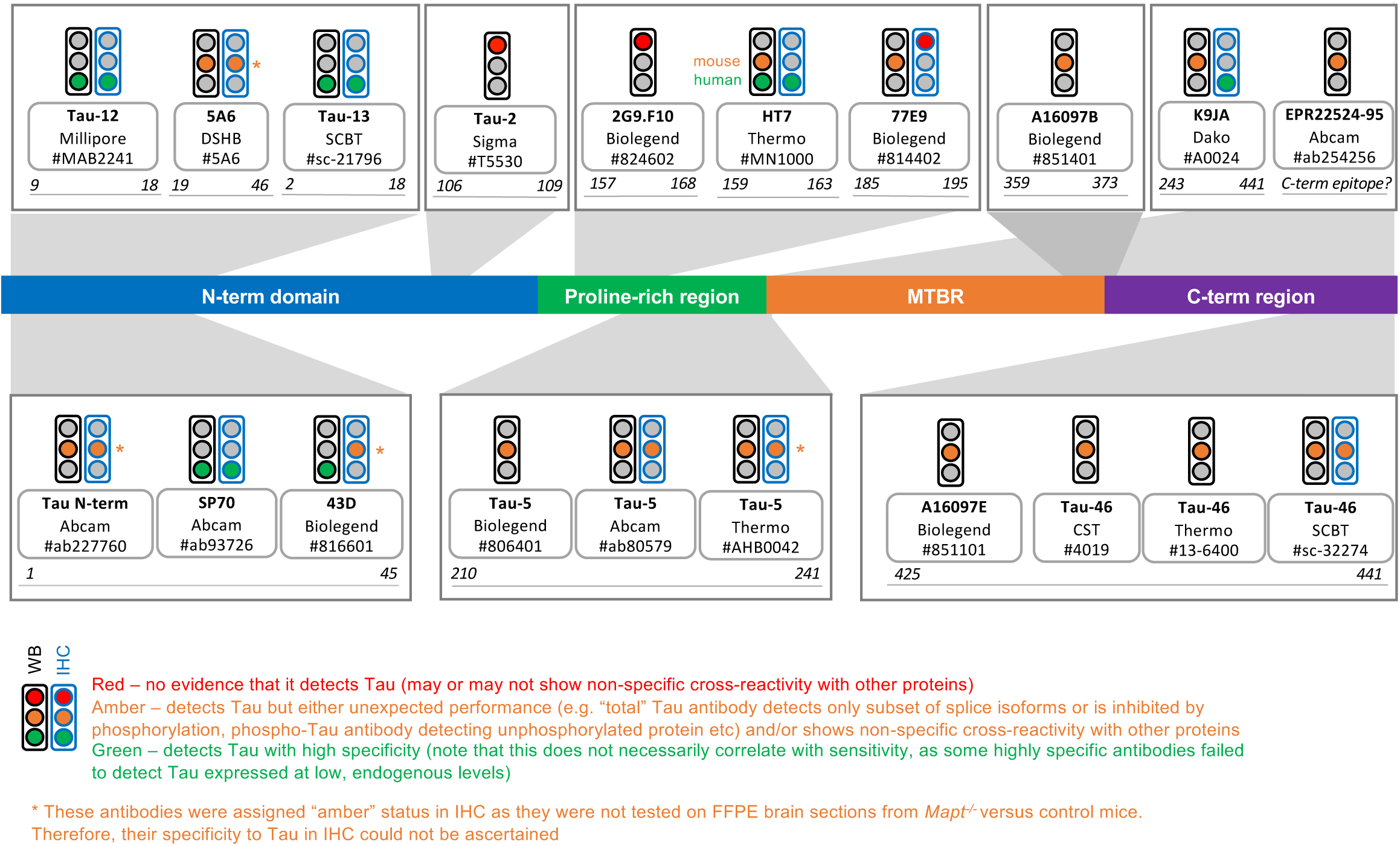
Summary of validation results for total Tau antibodies. Diagram of the Tau protein showing the locations of the epitopes of total Tau antibodies. Residue numbering is based on 2N4R Tau (441 amino acids; Uniprot ID 10636-8). Traffic lights summarise the performance of each antibody in WB (**black outline**) and IHC (**blue outline**). Based on data presented in this study, antibody performance was classified as either: **Green** = detects Tau with high specificity; **Amber** = detects Tau but either unexpected performance (e.g. “total” Tau antibody detects only subset of splice isoforms or is inhibited by phosphorylation, phospho-Tau antibody detecting unphosphorylated protein etc) and/or shows non-specific cross-reactivity with other proteins; or **Red** = no evidence that it detects Tau (may or may not show non-specific cross-reactivity with other proteins). Asterisks indicate antibodies whose performance in IHC was assigned “amber” status on the basis that these reagents were not tested in *Mapt^-/-^*, and corresponding control, FFPE mouse brain sections, such that their specificity in IHC could not be ascertained. Antibody clone HT7 was assigned two colours for performance on WB (“amber” for mouse samples, “green” for human samples), reflecting our findings that its specificity to Tau was species-dependent.

**Figure 18.**
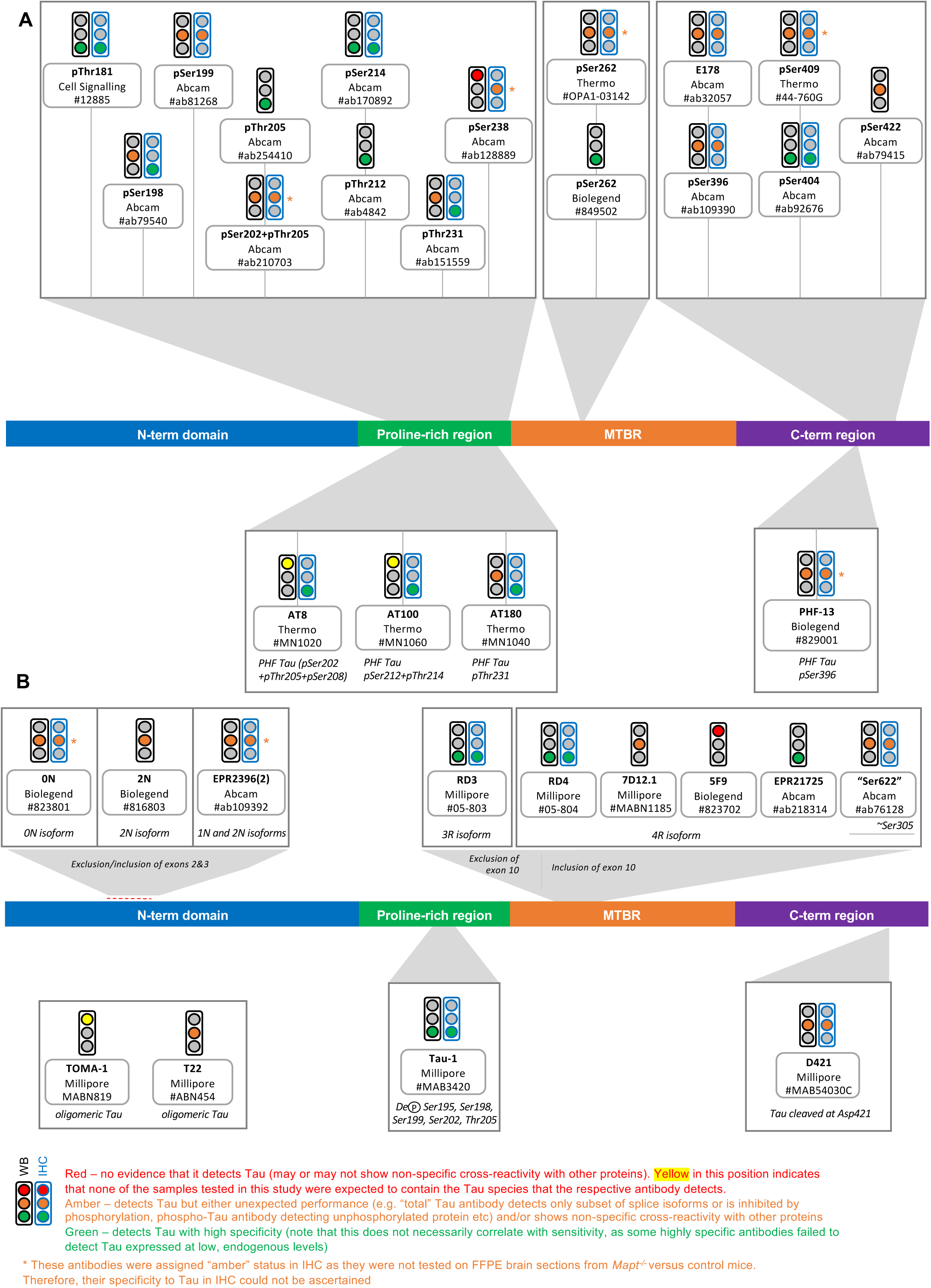
Summary of validation results for phosphorylation-dependent, other PTM-dependent and isoform-specific Tau antibodies. Diagram of the Tau protein showing the locations of the epitopes of phospho-Tau antibodies (**A**) and other-PTM dependent antibodies plus isoform-specific Tau antibodies (**B**). Residue numbering is based on 2N4R Tau (441 amino acids; Uniprot ID 10636-8). Traffic lights summarise the performance of each antibody in WB (**black outline**) and IHC (**blue outline**). Based on data presented in this study, antibody performance was classified as either: **Green** = detects Tau with high specificity; **Amber** = detects Tau but either unexpected performance (e.g. “total” Tau antibody detects only subset of splice isoforms or is inhibited by phosphorylation, phospho-Tau antibody detecting unphosphorylated protein etc) and/or shows non-specific cross-reactivity with other proteins; or **Red** = no evidence that it detects Tau (may or may not show non-specific cross-reactivity with other proteins).

### Antibody performance: ability to detect the target protein and specificity

Experimental validation of any antibody can be addressed by answering two deceptively simple key questions: (1) does the antibody bind its target protein? and (2) does it do so with a high degree of specificity?

The first question is the more straightforward and can be answered by using positive control samples known to contain the target epitope. Using WB and IHC-IF, we confirm that most of the antibodies tested do react with Tau, at least when the target is present at sufficiently high levels. Among these, however, several total Tau antibodies (notably Tau-5, HT7, Tau-46) failed to detect the protein in samples where the target protein is expressed at relatively low levels (wildtype HAP1 cells by WB and hTau mouse brains by IHC-IF), despite yielding positive signals in samples that express Tau at higher but still physiological levels (SH-SY5Y lysates by WB and wildtype mouse brains by IHC-IF). This illustrates clearly that the choice of primary antibody is critical for the reliable detection of Tau, especially when testing samples in which it is present at low levels. It is possible that some of these antibodies (e.g. Tau-5, Tau-46) may have higher affinity for mouse Tau than human Tau, in line with the findings of previous studies that some antibodies originally raised against bovine Tau can display up to 45-fold stronger immunoreactivity with the bovine protein compared to the human protein, despite high sequence homology in the respective epitope regions [158]. A subset of antibodies (clones Tau-2, 2G9.F10, pSer238, TOMA-1) failed to yield a detectable signal by WB, IHC or both, in any of the samples employed in this study. This may suggest that these antibodies do not detect Tau, that the target Tau species (e.g. phosphorylated, aggregated) are not present in the samples used in this study, or that such antibodies are sensitive to specific experimental conditions and may require further testing using alternative protocols to establish their utility. For example, it has been reported that the presence of detergents can influence the immunoreactivity of the Tau-2 clone [159,160].

Another important finding is that immunodetection with several reagents, whose binding to Tau is presumed to be PTM-independent, was affected by the state of Tau phosphorylation. This was true for several total Tau antibodies, as well as for the D421 clone, and was most noticeable by WB (where the most affected “total” Tau reagents were: #ab254256, 5A6, HT7, 77E9, Tau-5, Tau-46, A16097B and A16097E). Strikingly, detection of Tau with the 77E9 clone was impacted by phosphorylation to a similar extent as that observed for the Tau-1 clone, the latter being known to recognise Tau only when dephosphorylated at its target epitope, suggesting that 77E9 also preferentially detects a dephosphorylated version of Tau. These data have far-reaching implications with respect to the validity of assessments of the ratio of phospho-Tau to “total” Tau in tissues and biological fluids. They also imply that, for certain antibodies, detection and quantification of total Tau or caspase-cleaved Tau is likely to be achieved most accurately when performed on phosphatase-treated samples. As such, the choice of “total” Tau antibodies should be made with reference to experimental conditions.

Addressing the specificity of each antibody is a more challenging task that requires the use of appropriate negative control samples where expression of the target epitope is abolished or specifically reduced. Despite its straightforward definition, antibody specificity is not an absolute feature: instead, it must be defined relative to the sample type and to the experimental staining protocol, particularly with respect to any blocking steps and the choice of detection reagents. For example, Tau-12 and AT8 have previously been reported to display non-specific cross-reactivity in mouse brain samples [69]. However, both reagents detected Tau with high specificity under the experimental conditions employed here. One important, but often overlooked consideration is that the specificity of any given antibody may vary depending on the species of origin of the samples being analysed. This is illustrated by our WB data where different non-specific bands were detected by the HT7, K9JA, N-term #ab227760 and #ab109392 antibodies in mouse brain compared to human cell lysates.

The C-terminal half of Tau shares extensive sequence homology with that of MAP2 and, to a lesser extent, MAP4 [132], raising the possibility that some Tau antibodies may cross-react with either or both of these proteins. Here, we confirm cross-reactivity with MAP2 for several total (Tau-46, K9JA, A16097E and A16097B) and, unexpectedly phospho-Tau antibodies (pSer199, pSer202+pThr205 and pSer396). This is of particular concern for any applications where the MW of the protein being detected cannot be ascertained (e.g. IHC, ELISA, co-IP) but also for WB analyses on tissues or cell types that express 2N Tau isoforms, which migrate on SDS-PAGE to a similar apparent MW as MAP2c, thus precluding the clear discrimination between the MAP2 and Tau signals.

Another obvious criterion that can influence the specificity of target epitope detection is the proportion of that epitope relative to other proteins that are potential sources of non-specific binding (i.e. the signal-to-noise ratio). This is particularly relevant when the target protein is present in only small amounts. In previous work, antibodies directed against low-abundance epitopes, such as AT8 in the adult mouse brain, were found to be most prone to non-specific cross-reactivity [69]. In the present study, this principle is exemplified by the complementary observation that almost all antibodies appeared highly specific for Tau when tested on samples where the protein is expressed at high levels (e.g. lysates from Tau-overexpressing HEK293T cells on WB, or brain sections from rTg4510 mice by IHC). This also highlights the potential pitfalls associated with the use of overexpression systems to monitor antibody specificity [71,161]. Similarly, we found that cross-reactivity of the Tau-46 and Ser622 antibodies with other proteins only became apparent when they were used to stain brain sections from wildtype mice, but not from rTg4510 mice.

For PTM-dependent Tau antibodies, non-specific signals may also arise from cross-reactivity with the unmodified protein or with proteins carrying PTMs other than those targeted by the respective antibody. For example, our data highlight that immunoreactivity of the widely-used T22 antibody clone is not restricted solely to oligomeric Tau, since it also detected monomeric Tau on WBs. This feature was evident in several previous studies but had not been emphasised ([162], Fig. 4; [163], Fig. 7; [164], Fig. 3; [165], Fig. 3d - see “Tau 0 hrs” on dot blot; [166], Fig. 4A;[167], Fig. 2; [168], Fig. S15; [169], Fig. 4A-C). Using phospho-peptide arrays, previous work reported that several polyclonal phospho-Tau antibodies can also detect the unphosphorylated form of the protein [70]. Of the 20 phospho-Tau antibodies tested here, only pSer409, a polyclonal reagent, cross-reacted weakly with the recombinant Tau ladder on WB. For the remaining 19 phospho-Tau antibodies tested (one polyclonal, 18 monoclonals), our WB data support the conclusion that they are indeed phosphorylation-specific. However, our IHC findings with certain phospho-Tau antibodies may appear to contradict this statement, as immunostaining of pathological Tau aggregates appeared to remain unchanged (pSer199 antibody) or even improve (E178 and pSer396 antibodies) following λPP treatment. To explain this perplexing observation, we advance the hypothesis that this phenomenon is a consequence of: (1) the fact that phosphorylation at the relevant epitopes (e.g. pSer199 and pSer396) is resistant to λPP in aggregated/PHF-Tau, combined with (2) the fact that removal of potential inhibitory phosphorylation events at nearby residues may improve epitope accessibility. In support of this hypothesis, a previous study found that the binding of a different pSer396 antibody to a peptide carrying its target epitope was hindered by modifications at nearby residues [70].

It is widely understood that, when analysing mouse samples using primary antibodies that have been raised in mice (known as “mouse-on-mouse” immunostaining), endogenous IgG detection by anti-mouse secondary antibodies can contribute to non-specific signals. For WB applications, this can be particularly troublesome as IgG heavy chains migrate on SDS-PAGE at a similar apparent MW as monomeric Tau. In the present study, non-specific bands at ∼26 kDa and/or ∼55 kDa, consistent with the MW of the IgG light and heavy chains, respectively, were observed on WB for some but not all monoclonal mouse antibodies (exceptions include: Tau-12, Tau-13 and 77E9, presumably due to high signal-to-noise ratios). In contrast with previous reports [69] and with the exception of the D421, 0N and 2N antibody clones, we find that, even when present, these bands were only faintly detectable. For IHC, the protocols employed in this study did not yield detectable non-specific signals in brain sections incubated with the secondary antibodies alone, potentially due to any such signals being removed during the autofluorescence removal step of image processing (see Methods), thereby confirming that immunopositivity had resulted from the binding of the primary antibody.

### Non-canonical Tau-immunoreactive bands: low-MW and high-MW Tau isoforms?

In the present work, testing a large panel of antibodies allowed the identification of “non-canonical” bands that may otherwise have been dismissed as non-specific signals. Several of the antibodies tested (most notably total Tau antibodies SP70, 43D, Tau-12, Tau-13, 77E9 and K9JA, as well as PTM-dependent antibodies pSer198, pSer199, pThr205 and Tau-1) detected specific bands migrating at apparent MWs lower than expected for the shortest common brain Tau isoform, full-length 0N3R Tau. These may represent truncated Tau variants, whose presence has been reported extensively in experimental systems and biological samples, where they continue to attract interest as biomarkers as well as for their potential role in spreading Tau pathology between neuroanatomically-connected areas of the brain [170–175].

Tau isoforms of a MW lower than full-length Tau may also arise through unconventional mechanisms, e.g. exon skipping. Our work provides evidence that presumed knockout, *MAPT*-edited HAP1 cell lines express novel low-MW Tau isoforms that incorporate sequences encoded both upstream and downstream of the deletion sites, thus suggesting they have resulted from exon skipping. This finding has broad relevance for attempts to genomically engineer the *MAPT* locus in human cells, as we have recently found that neuronal progenitor cells derived from presumed “Tau knockout” human induced pluripotent stem cells (hiPSCs), that carry CRISPR-Cas9-induced deletions in *MAPT* exons 1 or 4, also continue to express low levels of Tau protein, similarly to our findings in HAP1 edited cells [176]. A 26-30 kDa small Tau isoform that may arise via exon skipping was previously described in SH-SY5Y cells [9], and experimental evidence has shown that “constitutively included” *MAPT* exons can be skipped to generate a gene product that integrates both termini [177]. Two previous independent studies on different panels of presumed “knockout” CRISPR-Cas9-edited HAP1 cell lines found that ∼30% of these continue to express the target protein, often as novel variants induced by skipping of the targeted exon [178,179]. Indeed, gene plasticity, evidenced as skipping of otherwise constitutively-included exons, is a widespread phenomenon induced by CRISPR-Cas9 genomic editing reported to occur in a wide variety of cell lines as well as mammalian, insect and plant models [180–184]. Skipping of *MAPT* exon 4, the exon that carries the CRISPR-Cas9-induced deletions in *MAPT*-edited HAP1 cells, would neither introduce an early stop codon nor alter the downstream amino acid sequence, but would decrease the theoretical MW of the protein, in line with our experimental observations.

Several phospho-Tau antibodies (e.g. pSer199, pThr231, pSer396, E178) detected prominent high-MW Tau-immunoreactive bands of approximately 110-120 kDa in SH-SY5Y and HAP1 cell lysates, a region of the WB membrane that is rarely shown in publications. When presented, signals in this region have variably been attributed to Tau dimers/oligomers, to non-specific cross-reactivity, less often to the Big/PNS-Tau isoform, or have remained unacknowledged. Defined Tau-immunoreactive bands – as opposed to a smear – in the region of 100-125 kDa have been detected in previous studies with various Tau antibodies in samples ranging from neuroblastoma cell lines [9,88,133,185–189], to rodent and human peripheral organs [19,24,190,191], PNS [192–194]and even brain [128,189,195–200]. As these bands were also detected with total Tau antibodies (e.g. Tau-12, Tau-13, K9JA), our data support the possibility that 110-120 kDa high-MW Tau-immunoreactive bands represent specific Tau signals, but further work is needed to establish the identity of these Tau species.

## Limitations

Due to the number of reagents tested and to enable side-by-side comparisons, we used a standardised immunodetection protocol for all antibodies. It is possible that further optimisation of factors, such as antigen retrieval, blocking, detergent concentration and antibody incubation conditions, may improve the apparent performance of some antibodies. Moreover, we did not seek to confirm the identity of each epitope by unequivocal empirical analysis. To mitigate this, we used monoclonal antibodies wherever possible, many of which were raised against well-defined synthetic peptides. For IHC applications, we focused on FFPE tissues, owing to their widespread use and availability. Similar approaches could be employed to validate Tau antibodies for use in IHC on frozen tissue sections.

## Conclusions and recommendations

Here, we have established a resource designed to guide informed antibody choice for the study of Tau in protein extracts by WB or *in situ* by IHC. Our work highlights the importance of carrying out detailed and explicit antibody validation by using complex biological samples representative of the samples of interest in terms of target protein expression levels and species of origin, two factors that can profoundly affect antibody performance. We recommend testing antibodies on a small set of relevant samples, where possible, to establish performance with the specific protocol and reagents available. This is especially pertinent given our observations that the outcomes obtained with the “same” antibody clone can vary depending on the commercial supplier.

On SDS-PAGE, we detected the presence of non-canonical Tau species of MWs both lower and higher than those of “common”/brain Tau splice isoforms. While many publications show only the ∼50-70 kDa region of the WB membrane when analysing Tau proteins, our data highlight the need for the entire membrane to be shown. We further demonstrate that presumed *MAPT* knockout cells express novel low-MW Tau isoforms that likely arise via exon skipping. We propose that future attempts to knockout *MAPT* in human cells should employ strategies that remove the transcription start site and generate either multiple deletions or deletions that span multiple exons, in order to ensure that production of all Tau protein variants is eliminated.

To the best of our knowledge, no truly “total Tau” antibody has emerged to date, since no single antibody can detect all products of proteolytic cleavage. A polyclonal antibody raised against Big/PNS-Tau may achieve this but its utility would likely be compromised by extensive non-specific cross-reactivity. Currently, if the goal is to detect the majority of Tau species, including cleaved variants, our data suggest that an optimal strategy would seek to employ a combined panel of N-terminal, mid-domain and C-terminal antibodies, taking into consideration the limitation that antibodies targeting the C-terminal region of Tau may cross-react with MAP2 and, potentially, also MAP4. Moreover, in light of our findings that phosphorylation can inhibit the binding of some total Tau antibodies, dephosphorylation of the samples prior to analysis may be desirable, although we cannot rule out the existence of further PTMs that may also influence binding.

It is remarkable that the efforts of previous validation studies have identified reliable commercial monoclonal phospho-Tau antibodies targeting only six phospho-epitopes: pSer198, pSer199, AT8, AT180, pSer404 and pSer422 [201]. Our work confirms the specificity of the pSer198, AT8, AT180 and pSer404 antibodies, and adds antibodies targeting pThr181, pThr205, pSer214 and pSer396 to this list. The pSer199 antibody clone tested here is different from that validated previously, while our data revealed that the pSer422 antibody, which was rated highly previously [70,201], shows non-specific cross-reactivity with a human protein but this is only apparent when it is tested on human cell lysates that express Tau at physiological levels. We also find that 19 of the 20 phospho-Tau antibodies tested did not react with unphosphorylated Tau proteins. Through comparison with antibodies raised against recombinant peptides carrying the relevant phospho-epitopes (pSer202+pThr205, pThr231 and pSer396), our data further suggest that the epitopes of some phospho-Tau antibodies raised against AD brain PHF-Tau (AT8, AT180 and PHF-13) remain incompletely characterised and merit further scrutiny.

Despite the biological and pathological relevance of Tau splice isoforms, our work highlights the unmet need for antibodies that detect the different isoforms with high specificity and sensitivity. In particular, antibodies against 0N or 2N Tau demonstrated poor performance, while no currently-available antibodies target human 1N Tau isoforms alone. Antibodies targeting 4R Tau also performed poorly, either in terms of specificity or sensitivity.

The overarching aim of this work was to identify antibodies that can be used reliably to detect Tau with high specificity in tissues or cell types where the protein is expressed at endogenous (often very low) levels. Towards this, we identify a panel of high fidelity “total”, isoform-specific and phospho-Tau antibodies (including “PTM-independent” antibodies SP70, Tau-12, Tau-13 and RD3). In doing so, this work opens the door to studying this complex protein in peripheral tissues where its expression and functions remains largely unexplored. We anticipate that improved understanding of Tau antibody reagents will enable better data interpretation and enhance reproducibility in Tau research studies.

## List of abbreviations

λPP: lambda phosphatase
AD: Alzheimer’s disease
CNS: central nervous system
CRISPR: Clustered Regularly Interspaced Short Palindromic Repeats
FFPE: formalin-fixed, parrafin-embedded
GGT: globular glial tauopathy
hTau: humanised Tau mice (transgenic for the entire human *MAPT* locus on a mouse *Mapt*^-/-^ background)
IF: immunofluorescence
IgG: immunoglobulin G
IHC: immunohistochemistry
MW: molecular weight
PHF: paired helical filaments
PiD: Pick’s disease
PNS: peripheral nervous system
PTM: post-translational modification
SDS-PAGE: sodium dodecyl sulphate-polyacrylamide gel electrophoresis
ThS: Thioflavin S
WB: Western blot

## Declarations

### Ethics approval and consent to participate

Anonymised human brain tissue was obtained from the Oxford Brain Bank. This material was obtained by Oxford Brain Bank from humans with appropriate consent as required by the Human Tissue Act 2004 and with ethical approval (REC name: South Central – Oxford C; REC ref: 15/SC/0639; IRAS ID: 103696).

### Consent for publication

not applicable

### Availability of data and materials

All antibodies mentioned in this manuscript are commercially available. The mass spectrometry proteomics data have been deposited to the ProteomeXchange Consortium via the PRoteomics IDEntification Database (PRIDE) [202] partner repository with the dataset identifier PXD041199. All other data on which conclusions are based are included within the manuscript and its additional files, and raw data files are available from the corresponding authors upon reasonable request.

### Authors’ contributions

MJE performed the majority of the laboratory experiments for the validation of antibodies by WB; CL performed laboratory experiments for the validation of antibodies by IHC-IF, quantified IHC-IF microscopy data and prepared figures for publication; HT performed the immunoprecipitation experiments on HAP1 cells and performed laboratory experiments for the validation of antibodies by WB; SD performed laboratory experiments for the validation of antibodies by IHC-IF; MLZ performed laboratory experiments for the validation of antibodies by IHC-IF; DPOB processed immunoprecipitation eluates for proteomic analysis, analysed these samples on LC/MS-MS and analysed the raw proteomics data; JGK dissected the brains of wildtype, *Mapt^-/-^* and hTau mice; BMK supervised DPOB; NGM gave critical feedback on experimental design, helped secure funding and co-supervised CL; JAT gave critical feedback on experimental design and helped secure funding; SJR supervised CL, designed experiments, interpreted data, helped secure funding and contributed to manuscript writing; MIS designed the study, supervised MJE, helped secure funding, performed laboratory experiments for the validation of antibodies by WB, quantified WB data, interpreted data, prepared figures for publication and wrote the first draft of the manuscript. All authors commented on the manuscript.

## Acknowledgements

Elly Lilly for providing rTg4510 mice to John Brown (University of Exeter). Milena Cioroch and Richard Wade-Martins for providing wildtype, *Mapt^-/-^* and hTau mice. Pia Leete and Christine Flaxman, University of Exeter, for providing technical assistance with mouse brain tissue sectioning and IHC protocols. For providing access to FFPE human brain sections, we acknowledge the Oxford Brain Bank, supported by the Medical Research Council (MRC), the NIHR Oxford Biomedical research Centre and the Brains for Dementia research programme, jointly funded by Alzheimer’s Research UK and Alzheimer’s Society.

## Additional files

**Supplementary Table S1. List of all antibodies used in this study**. Categories shown for all antibodies are as follows: **A**, antibody name and clone name (where applicable, shown in square brackets after the antibody name); **B**, host species; **C**, clonality; **D**, isotype; **E**, commercial supplier; **F**, catalog number. In addition, where applicable and where this information is available, either from the manufacturer or in previous studies, the following information was also included: **G**, antibody concentration as reported by the supplier; **H**, number of studies citing each antibody according to Citeab; **I**, dilution used for WB applications; **J**, dilution used for IHC-IF applications; **K**, immunogen; **L**, targeted epitope; **M**, species reactivity.

**Supplementary Figure S1.**
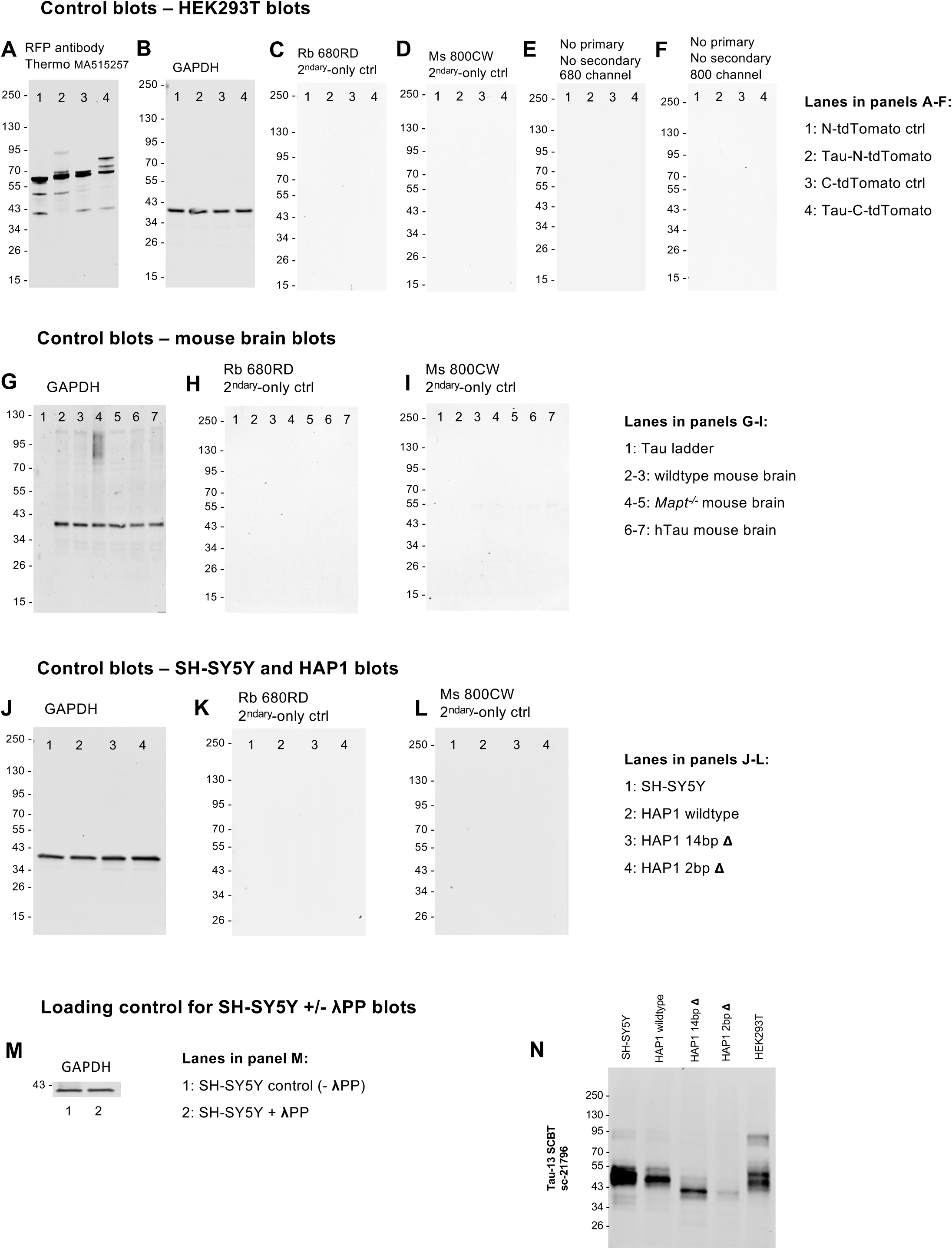
Control WB membranes. **A, G, J, M:** WB membranes probed with an anti-GAPDH antibody as a loading control. **B**: WB membrane of HEK293T cell lysates probed with an anti-RFP antibody to detect tdTomato expression. **C, H, K**: WB membranes probed with the anti-rabbit secondary antibody only, in the absence of a primary antibody, to identify non-specific signals that may arise from binding of the secondary antibody. **D, I, L**: WB membranes probed with the anti-mouse secondary antibody only, in the absence of a primary antibody, to identify non-specific signals that may arise from binding of the secondary antibody. **E, F**: WB membranes of HEK293T cell lysates were subjected to the same protocol as all other WB membranes shown in this study, with the exception that the primary and secondary antibodies were omitted, in order to assess background autofluorescence in the 700 nm (**E**) and 800 nm (**F**) detection channels, respectively. Order of samples for HEK293T blots (**A-F**), mouse brain blots (**G-I**), SH-SY5Y and HAP1 blots (**J-L**) and SH-SY5Y+/-λPP (**M**) is the same as that shown in Figs. 2-11, in columns one, two, three and five, respectively. **N:** To detect endogenous Tau expression in HEK293T cells, protein extracts (50 µg protein/lane) from SH-SY5Y, HAP1 and HEK293T cell lines were loaded on SDS-PAGE. WB membrane was probed with the Tau-13 total Tau antibody.

**Supplementary Figure S2.**
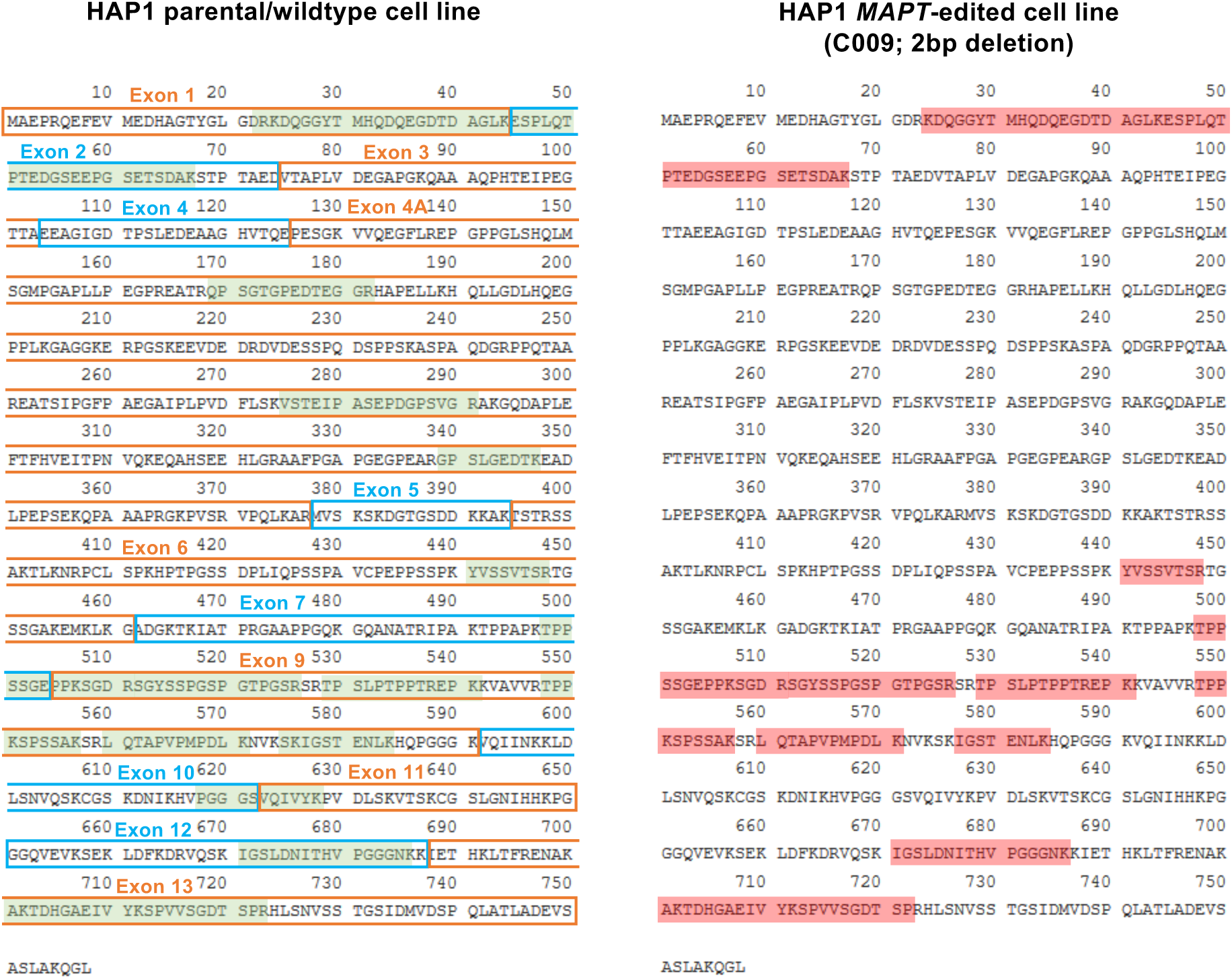
Tau peptides detected by mass-spec in cytoplasmic fractions from wildtype and *MAPT*-edited HAP1 cells following immunoprecipitation of Tau. Peptides detected in the wildtype/parental cell line (left, detected peptides highlighted in green) and in the 2-bp deletion *MAPT*-edited HAP1 cell line (right, detected peptides highlighted in red) are shown mapped to the amino acid sequence of the canonical Uniprot human Tau isoform (Uniprot ID P10636-1; 758 amino acids). Sequences encoded by the different *MAPT* exons are outlined and labelled on the left in alternating orange/blue colours.

**Supplementary Figure S3.**
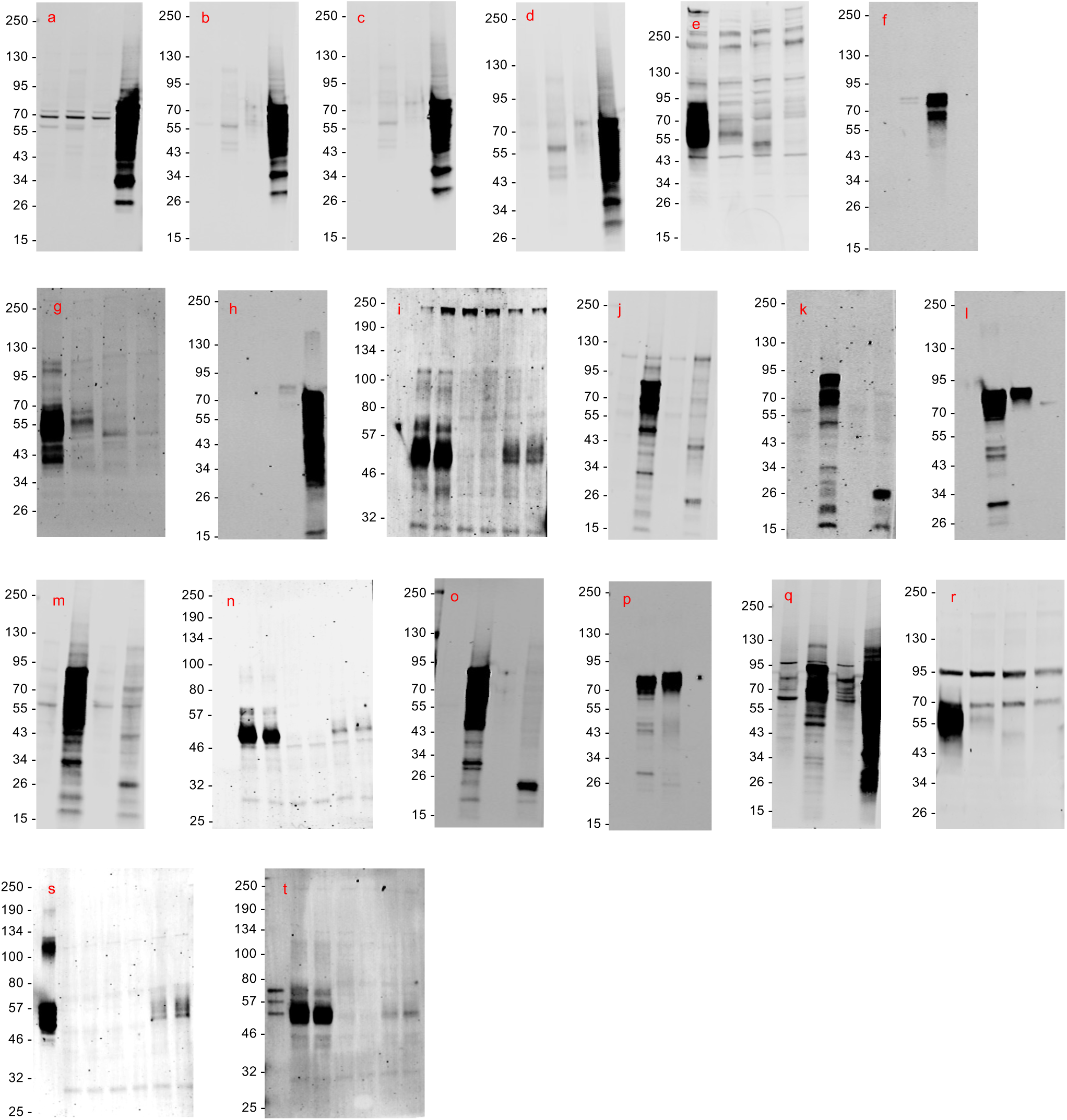
Related to Figs. 2-11: Images of WB membranes probed with different Tau antibodies where adjusting the brightness/contrast display settings led to the oversaturation of the “main” Tau signal but revealed additional bands. Brightness/contrast-adjusted blot images are shown in this figure. Red lettering shown in the upper left corner links each blot to the corresponding one shown in Figs. 2-11.

**Supplementary Figure S4.**
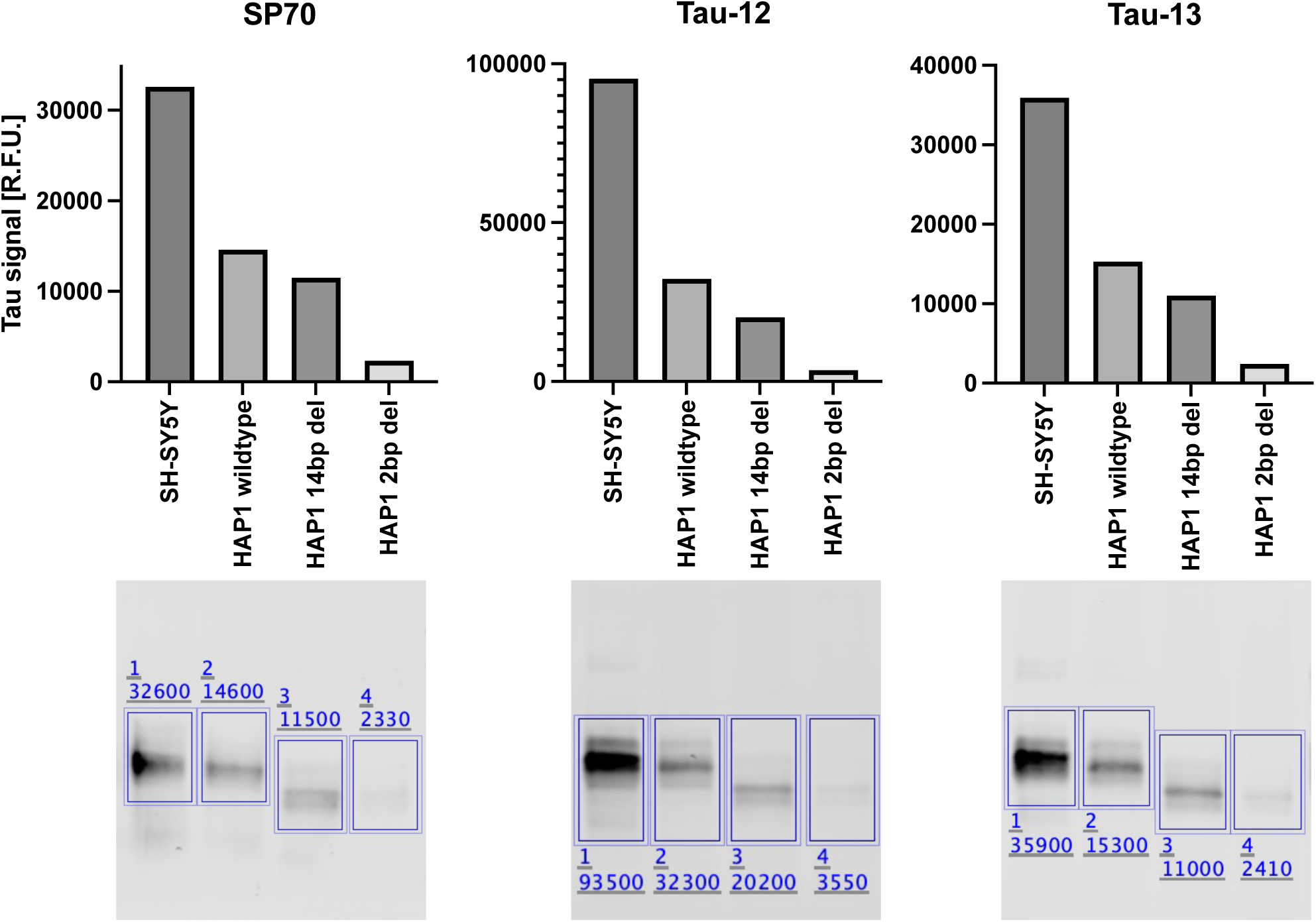
Tau protein expression levels in SH-SY5Y and HAP1 cell lines. Quantifications of the Tau band intensities detected by WB in lysates from SH-SY5Y, HAP1 wildtype, HAP1 14 bp Δ and HAP1 2 bp Δ cells as detected with the SP70 (**A**), Tau-12 (**B**) and Tau-13 (**C**) antibodies. Shown below each graph is the respective WB membrane with the areas used for quantifications marked by rectangles and the detected signal intensity value (R.F.U.) shown above each rectangle. The WB membranes that were used for quantifications and are shown in this figure are the same ones as those shown in the third column in Figs. 2D, 2F and 3A. Raw signal intensity values are reported as relative fluorescence units (R.F.U.).

**Supplementary Figure S5.**
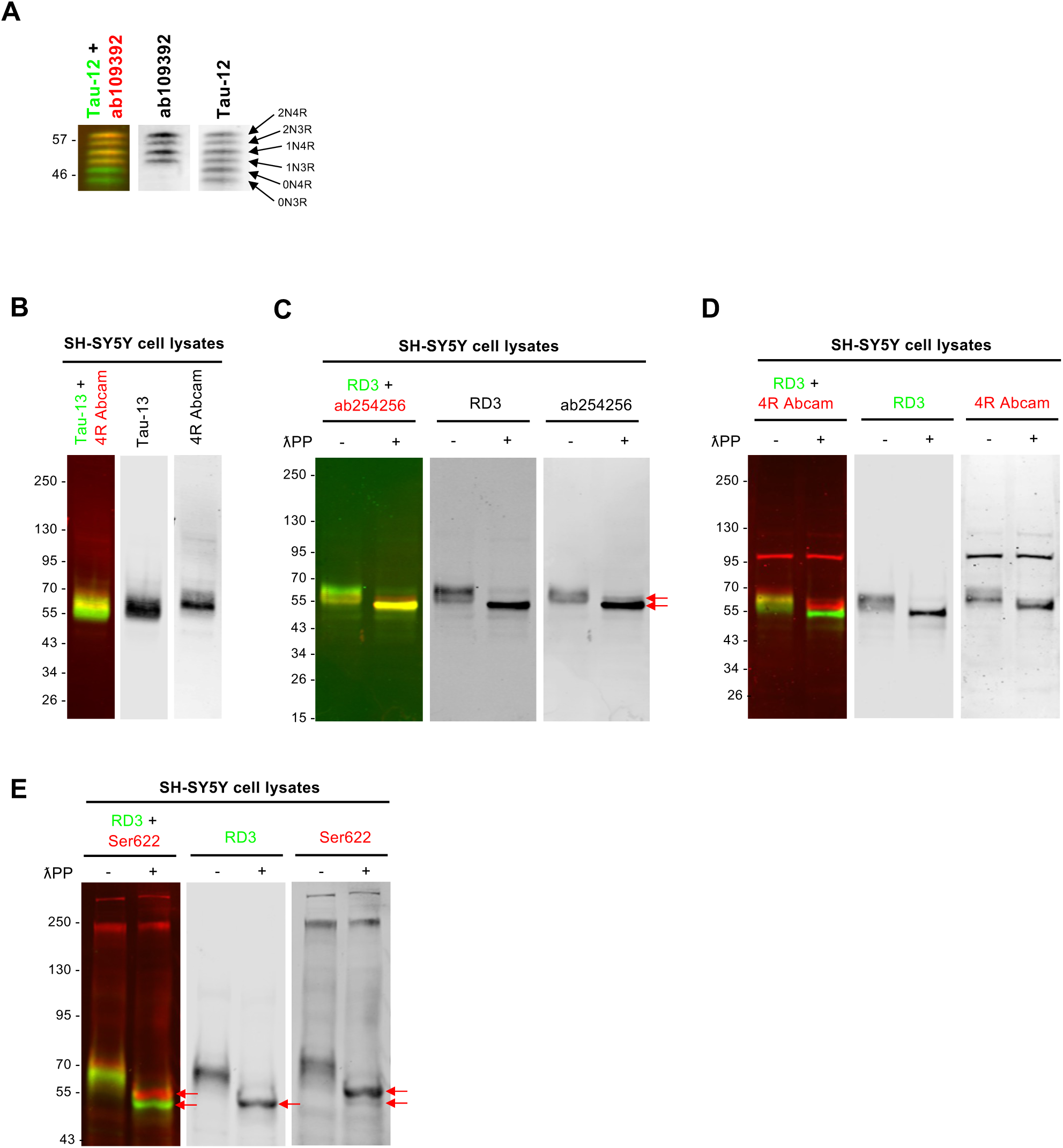
Additional data for the validation of isoform-specific Tau antibodies by WB (related to Figs. 10 and 11). **A**: Recombinant Tau ladder was detected with the Tau-12 total Tau antibody and the #ab109392 antibody, which only detects four of the six Tau isoforms that make up the ladder. The bands detected by #ab109392 correspond to 1N and 2N splice isoforms. **B-E**: Fluorescent WB membranes of SH-SY5Y cell lysates, that were either untreated (panel **B**, marked with “-” in panels **C-E**) or treated (marked with “+” in panels **C-E**) with λPP, were co-probed with different combinations of mouse and rabbit monoclonal Tau antibodies: Tau-13 total Tau antibody + 4R Abcam Tau antibody (**B**), RD3 3R Tau antibody + #ab254256 total Tau antibody (**C**), RD3 3R Tau antibody + 4R Abcam Tau antibody (**D**), RD3 3R Tau antibody + “Ser622” Tau antibody (**E**). Red arrows indicate the Tau bands detected in λPP-treated SH-SY5Y lysates that correspond to 3R (bottom band) and 4R (top band) Tau isoforms, respectively (**C**, **E**).

**Supplementary Figure S6.**
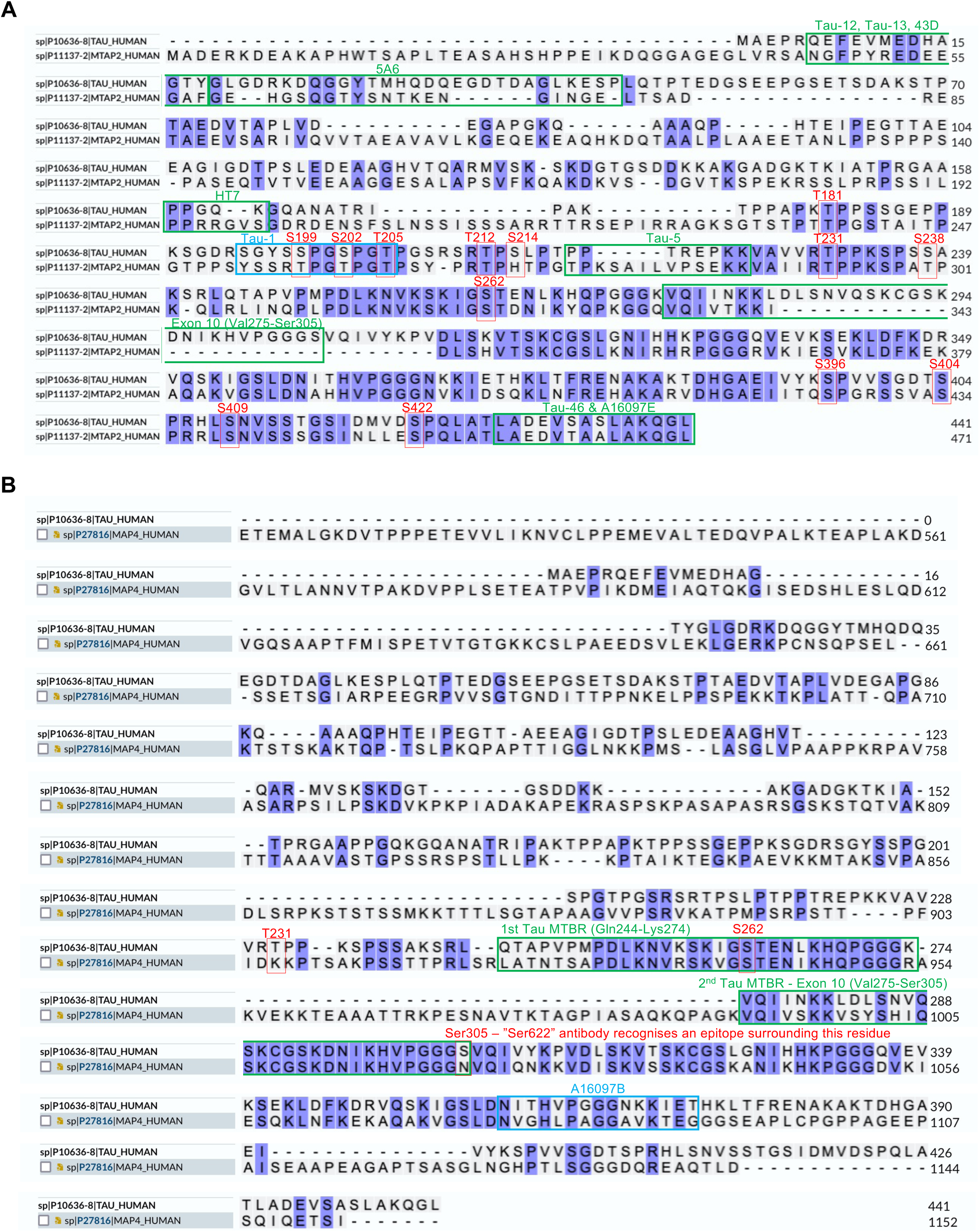
Alignment of protein sequences highlights similarities between Tau and the related microtubule-associated proteins, MAP2 and MAP4. **A**: CLUSTAL Omega alignment of the 2N4R human Tau protein sequence (441 amino acids, Uniprot ID P10636-8) with the human MAP2c protein isoform sequence (471 amino acids, Uniprot ID P11137-2). Phosphorylateable residues targeted by different phospho-Tau antibodies are highlighted in red. The epitope regions of different total Tau antibodies are highlighted in green and that of Tau-1 in blue. **B**: CLUSTAL Omega alignment of the 2N4R human Tau protein sequence (441 amino acids, Uniprot ID P10636-8) with the human MAP4 protein sequence (1,152 amino acids, Uniprot ID P27816). Human Tau residues Thr231, Ser262 and Ser305 are highlighted in red. The sequences of the 1st and 2nd Tau MTBRs (the latter is encoded by the alternatively-spliced exon 10 and is only present in 4R Tau isoforms) are highlighted in green. Epitope region of the A16079B total Tau antibody clone is highlighted in blue.

**Supplementary Figure S7.**
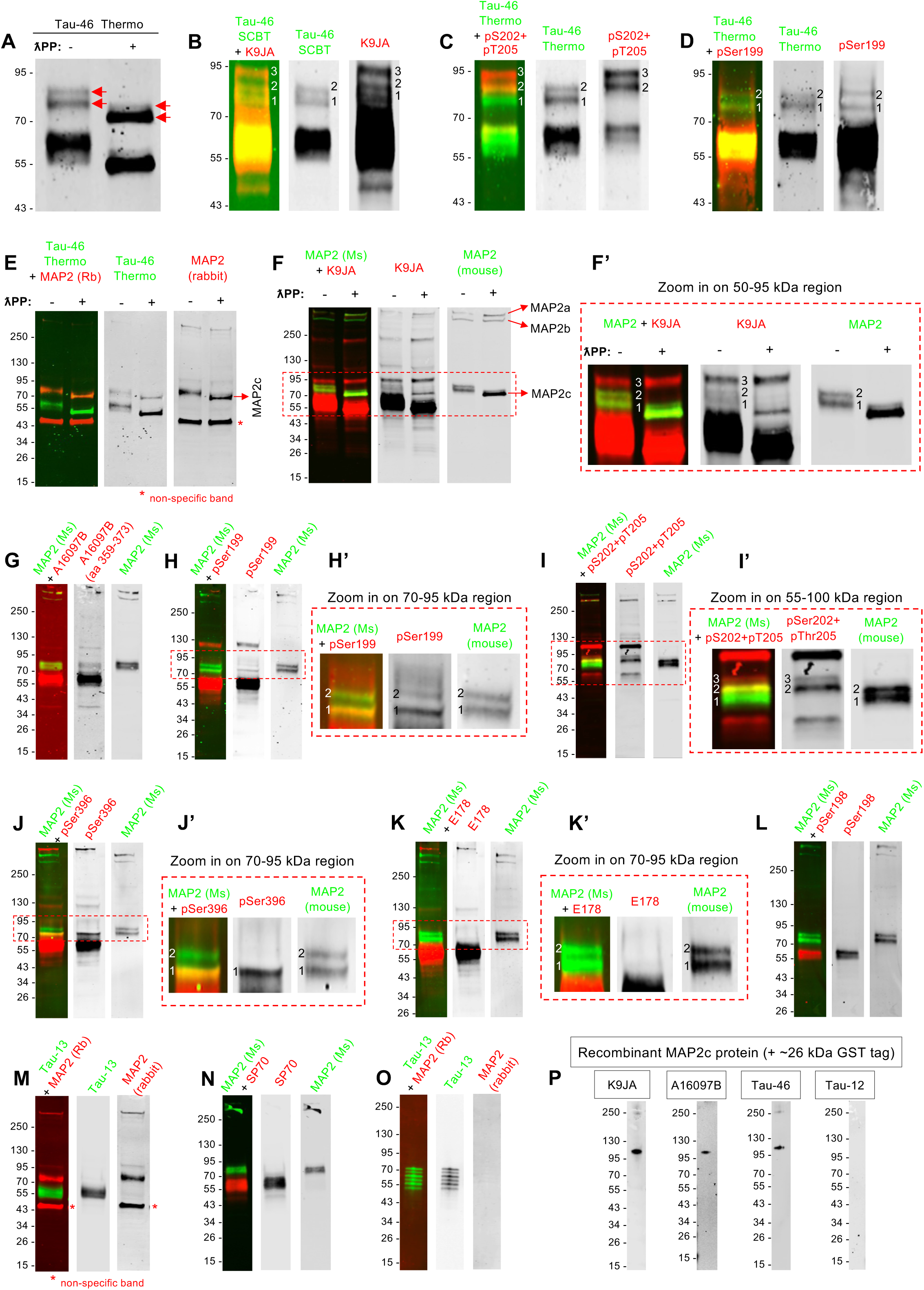
Some Tau antibodies cross-react with MAP2-immunoreactive bands on WB. Fluorescent WB membranes probed with different Tau and MAP2 antibodies. SH-SY5Y protein extracts were employed for all WBs shown in panels **A-N**. For panels **O** and **P**, recombinant proteins were used: recombinant Tau ladder (**O**) and recombinant GST-tagged MAP2c protein (**P**). **A**: Magnified view of the 43-100 kDa region of the SH-SY5Y +/-λPP WB membrane shown in Fig. 4, to highlight mid-MW Tau-46-immunoreactive bands (red arrows). **B-E**: WB membranes co-probed with Tau-46 and either: K9JA (**B**), pSer202+pThr205 (**C**), pSer199 (**D**), or MAP2 (**E**) antibodies. **F-N**: WB membranes co-probed with a MAP2 antibody and one of several Tau antibodies: K9JA (**F, F’**), A16097B (**G**), pSer199 (**H, H’**), pSer202+pThr205 (**I, I’**), pSer396 (**J, J’**), E178 (**K, K’**), pSer198 (**L**), Tau-13 (**M**), or SP70 (**N**). **O**: WB membrane of recombinant Tau ladder co-probed with the Tau-13 and MAP2 antibodies, to demonstrate lack of reactivity of the MAP2 antibody with Tau. **P:** WB membranes of recombinant MAP2c (carrying a ∼26 kDa GST tag) probed with one of four different Tau antibodies, as indicated (K9JA, A16097B, Tau-46 or Tau-12) to directly test for cross-reactivity of Tau antibodies with the MAP2 protein. In panels A, E, F and F’, protein lysates were either untreated (-) or treated (+) with λPP. Bands marked with an asterisk in panels E and M represent non-specific cross-reactivity of the rabbit MAP2 antibody. Magnified views of the regions outlined with red dashed line in panels F, H, I, J and K are provided in panels F’, H’, I’, J’ and K’, respectively. Bands corresponding to different MAP2 protein isoforms are indicated with red arrows in panels E and F.

**Supplementary Figure S8.**
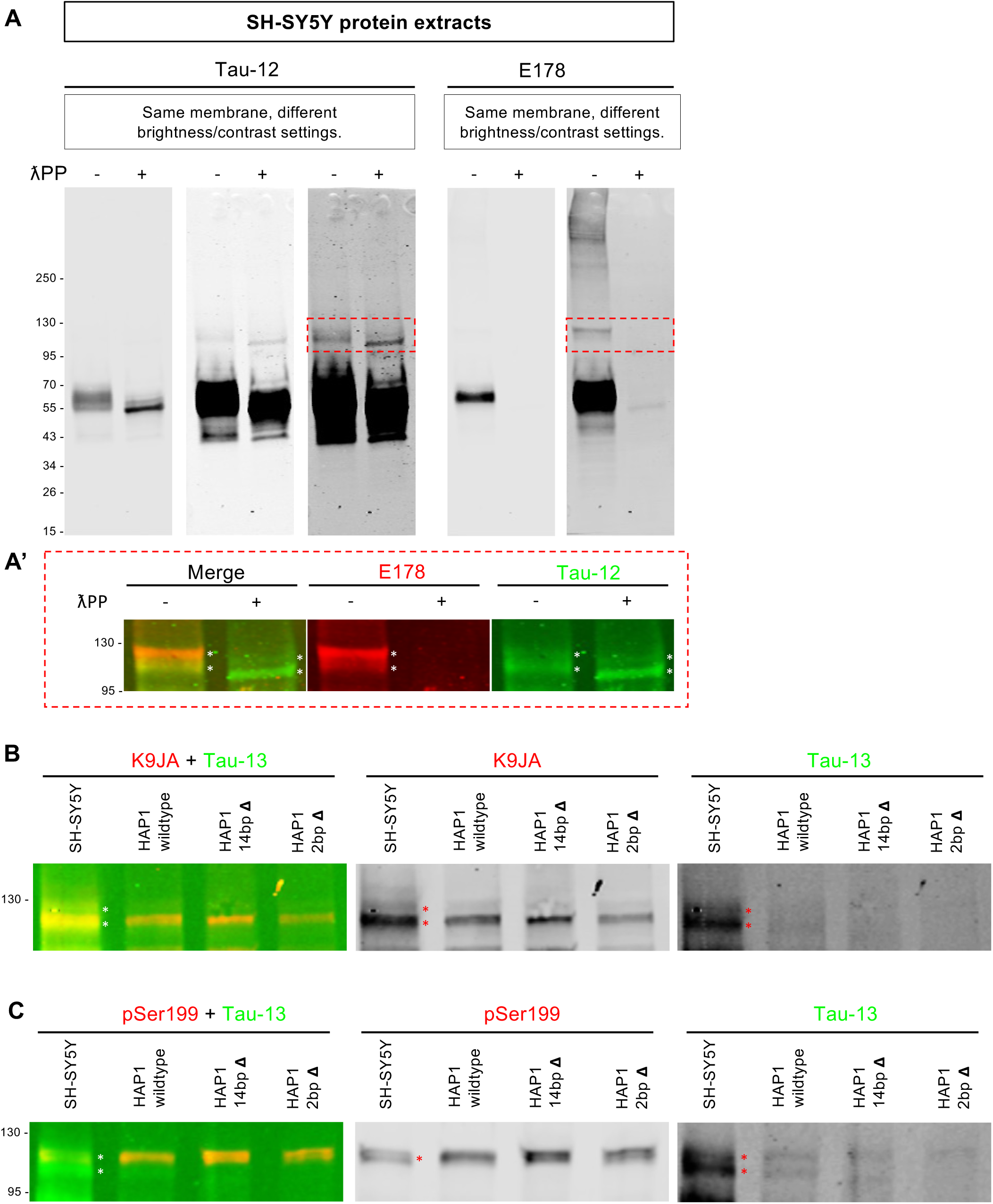
Several different Tau antibodies detect high-MW (HMW) Tau-immunoreactive bands in SH-SY5Y protein extracts. **A**: Fluorescent WB membranes of SH-SY5Y protein extracts, that were either untreated (-) or treated (+) with λPP, were sequentially probed with two different Tau antibodies, first with Tau-12 and then with E178. For each channel, the same membrane is shown at different exposure settings to allow the visualisation of HMW Tau-immunoreactive bands. **A’**: Magnified view of the region outlined with dashed, red line in panel A. The merged view of the two fluorescent WB detection channels as well as each individual channel are shown to illustrate the overlap of the HMW bands detected with the two antibodies (E178 was visualised in the 700 nm channel - red; Tau-12 was visualised in the 800 nm channel - green) **B**, **C**: Fluorescent WB membranes were co-probed with different combinations of mouse and rabbit Tau antibodies: Tau-13 + K9JA (**B**) or Tau-13 + pSer199 (**C**). For all panels, membranes were probed and scanned sequentially using the respective total Tau mouse antibody first, as the signal detected with these antibodies was weaker, in order to eliminate the possibility that the HMW band detected with these antibodies may result from cross-reactivity with the phospho-Tau or total Tau rabbit antibodies. Changing the order of the antibodies gave the same result (data not shown). We also confirmed that there was no bleed-through between the 800 nm (green) and the 700 nm (red) channels. Asterisks indicate the two HMW Tau-immunoreactive bands detected (**A’, B, C**).

**Supplementary Figure S9.**
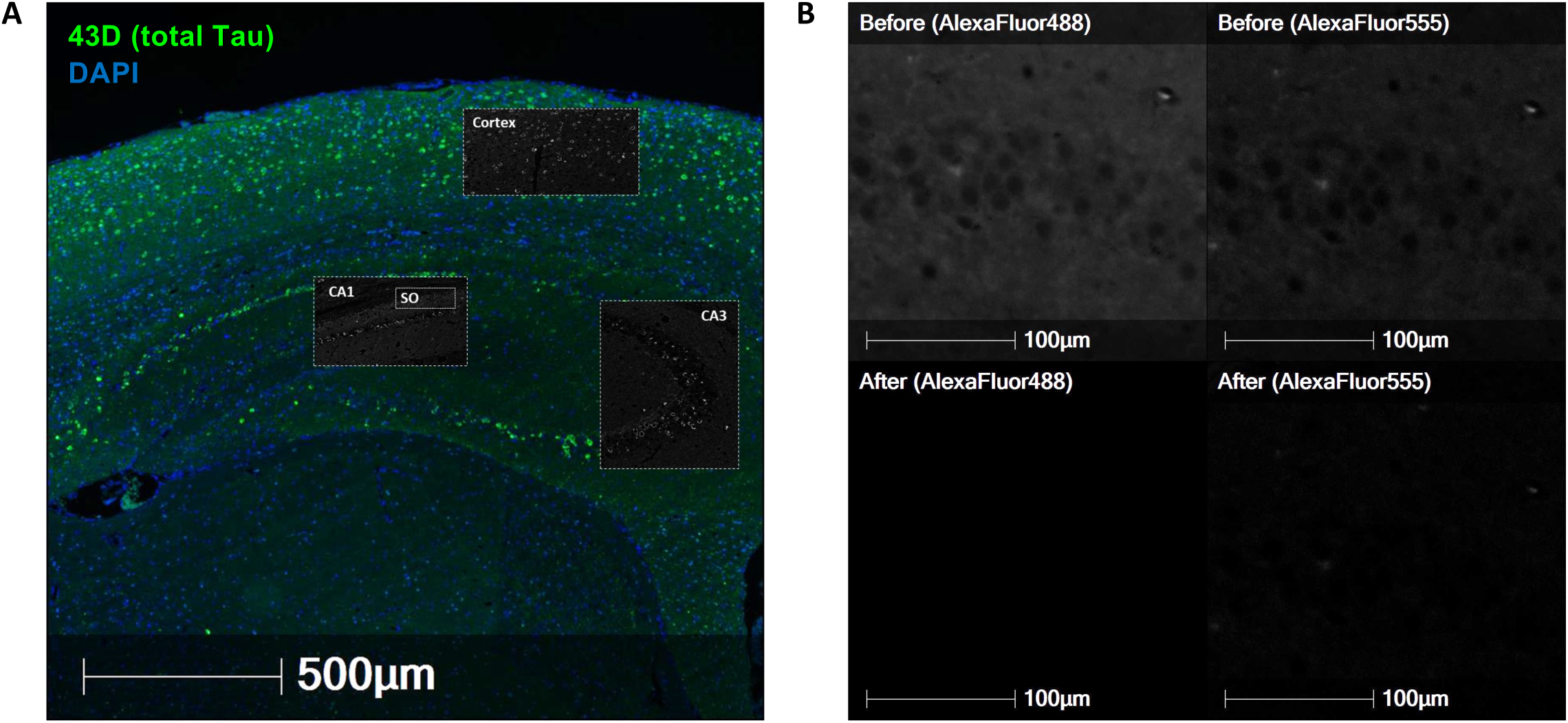
(**A**) Overview of mouse brain regions (cortex, CA1, CA3, SO - outlined with white dashed line) imaged for the validation of Tau antibodies by IHC. Brain section shown originates from rTg4510 mice, was treated with λPP and was stained with the 43D Total tau antibody (shown in green outside insets; shown in grayscale inside insets). Nuclei are stained with DAPI (blue, outside insets). Scale bar = 500 µm. CA = *cornu ammonis*, SO = *stratum oriens*. (**B**) Micrographs of CA3 brain region from rTg4510 mouse FFPE brain sections immunostained with AlexaFluor-conjugated secondary antibodies (AlexaFluor488 - left; AlexaFluor555 - right), in the absence of primary antibodies, before (top row) and after (bottom row) autofluorescence removal. Respective fluorescent signals are shown in grayscale. Scale bars = 100 µm.

**Supplementary Figure S10.**
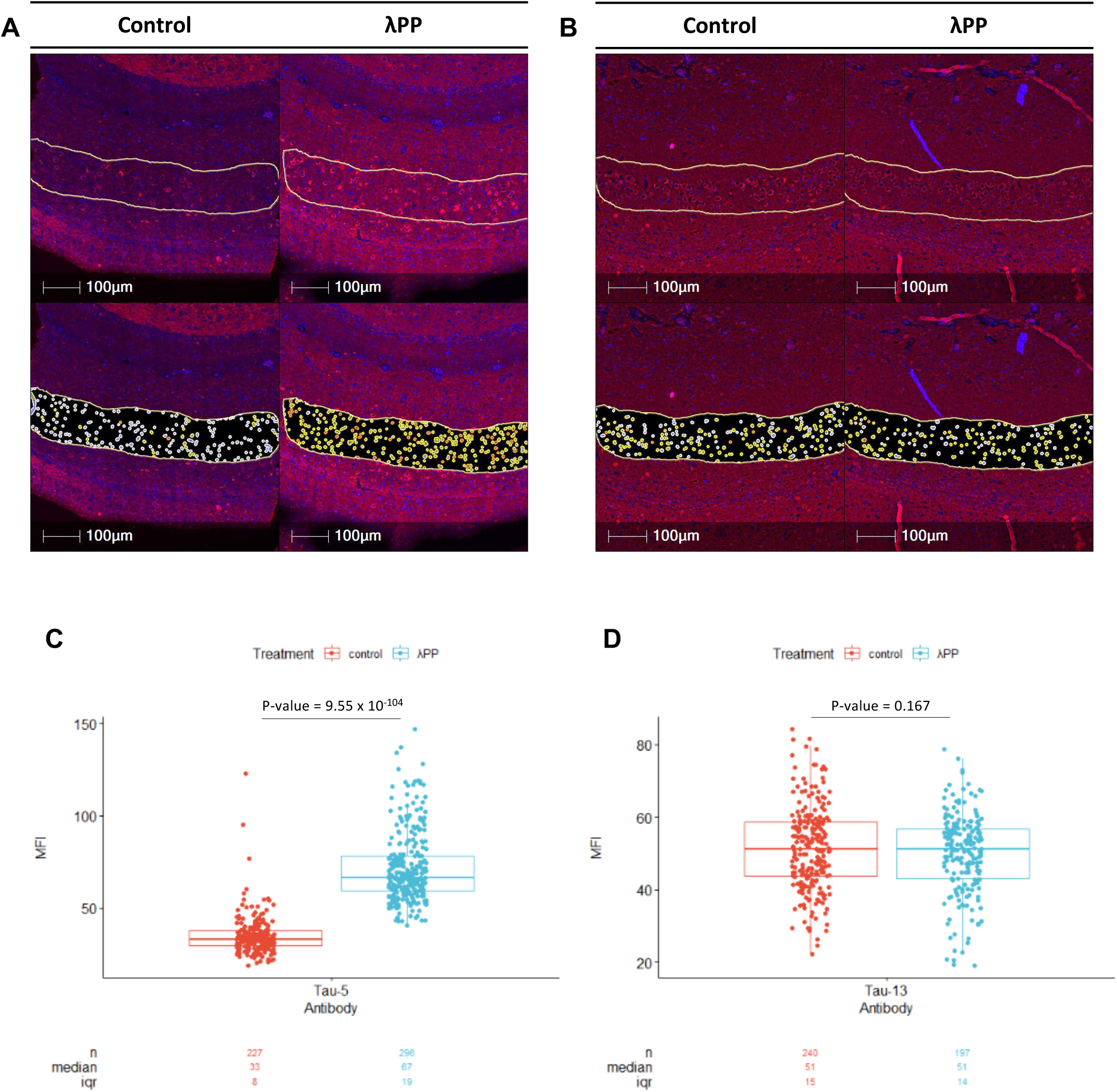
Phosphorylation partially inhibits the binding of the Tau-5, but not Tau-13, antibody clone in FFPE-IHC applications. Micrographs (**A, B**) and quantifications (**C, D**) of the signal mean fluorescence intensity (MFI) obtained in rTg4510 mouse brain cortex stained with the Tau-5 (**A, C**) or Tau-13 (**B, D**) antibodies, before (**A, B** - left; **C, D** - red) and after (**A, B** - right; **C, D** - blue) λPP treatment. Tau signal is shown in red (**A, B**). Nuclei are stained with DAPI (blue) (**A, B**). In A and B, the region used for quantifications is outlined and images in the bottom row highlight all the cells identified by the automated image analysis pipeline. Tau-negative cells are coloured in white (**A, B** - bottom row). Tau-positive cells are coloured in yellow and orange, indicating moderate and high Tau staining signal intensity, respectively (**A, B** - bottom row). Scale bars = 100 µm (**A, B**). n= total number of cells analysed, iqr = inter-quartile range (**C, D**). p-values were calculated using a two-tailed, independent *t test*.

**Supplementary Figure S11.**
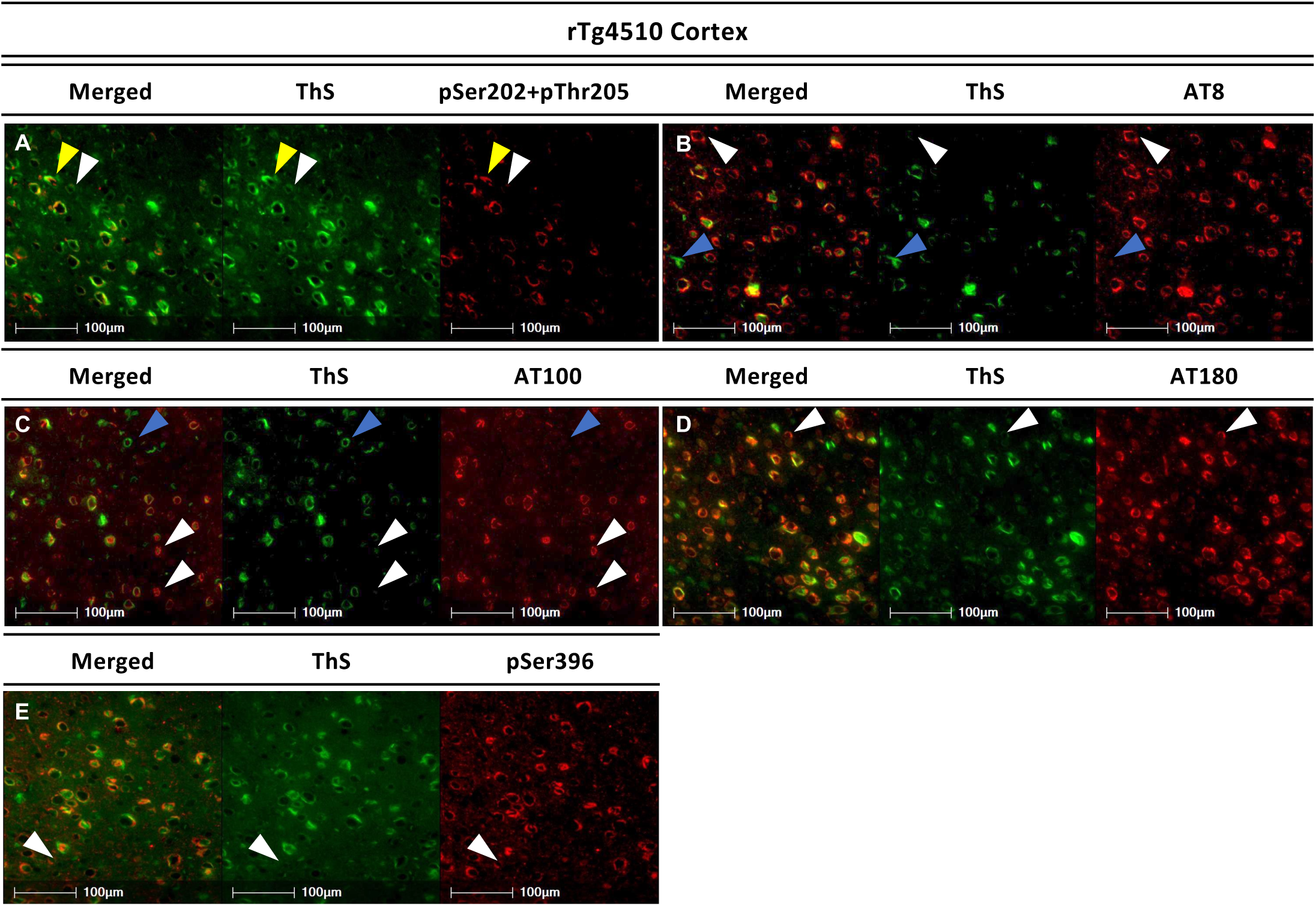
Majority of phosphorylated Tau in rTg4510 cortex is found in protein aggregates. FFPE rTg4510 mouse brain sections co-stained with Thioflavin S (ThS, green) and phospho-Tau antibodies (red): pSer202+pThr205 (**A**), AT8 (**B**), AT100 (**C**), AT180 (**D**), pSer396 (**E**). Cortex region outlined in Supp. Fig. 9A is shown. Scale bars = 100 µm. A small subset of cells were positive for phospho-Tau but negative for ThS (white arrowheads), or positive for ThS but negative for the respective phospho-Tau antibody (blue arrowheads).

**Supplementary Figure S12.**
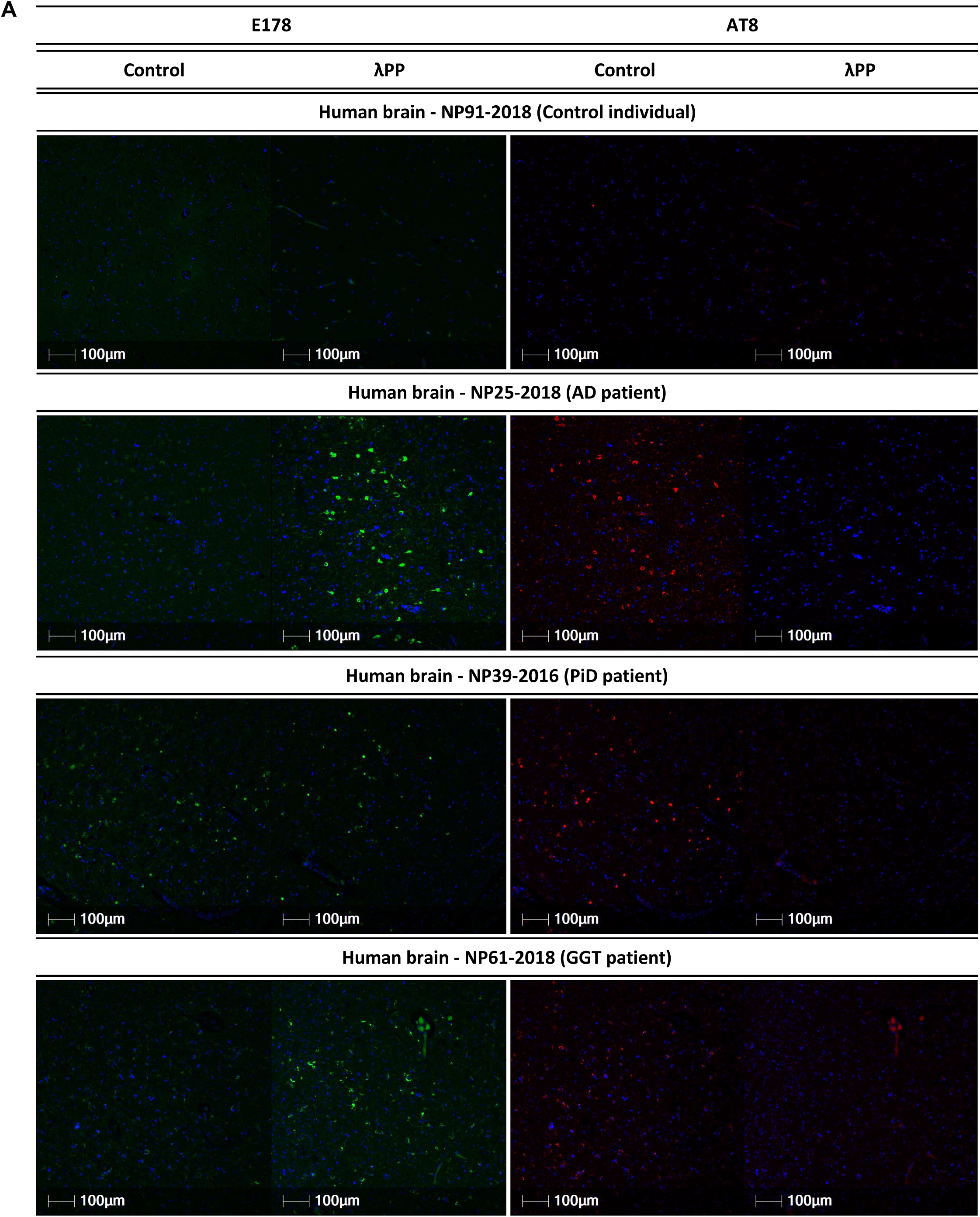
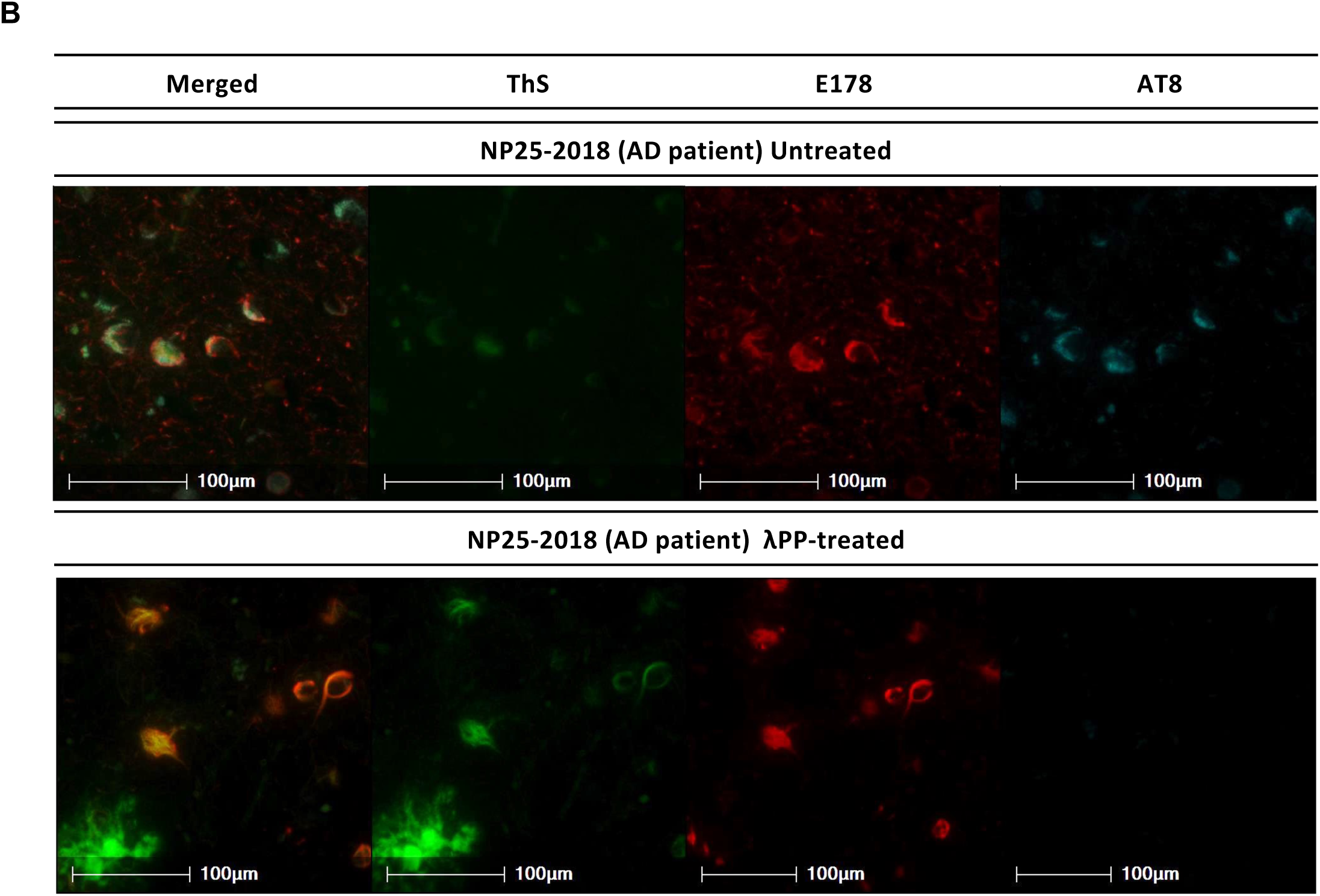
E178 immunoreactivity to pathological PHF-Tau present in human brain sections is enhanced, rather than abrogated, following λPP treatment. **A**: Serial FFPE brain sections from control, AD, PiD and GGT individuals were either untreated (control) or treated (λPP) with λPP prior to immunostaining with two different phospho-Tau antibodies: E178 (green) and AT8 (red). Nuclei were stained with DAPI (blue). Scale bars = 100µm. **B**: Serial FFPE brain sections from an AD patient were pre-incubated with λPP (bottom row) or left untreated (top row), before immunostaining with two different phospho-Tau antibodies: E178 (red) and AT8 (cyan). Sections were then stained with ThS (green) to identify aggregated proteins. Areas that appear positive for ThS staining but negative for phospho-Tau immunostaining are presumed to represent amyloid deposits. Scale bars = 100µm.

**Supp. Fig. S13:**
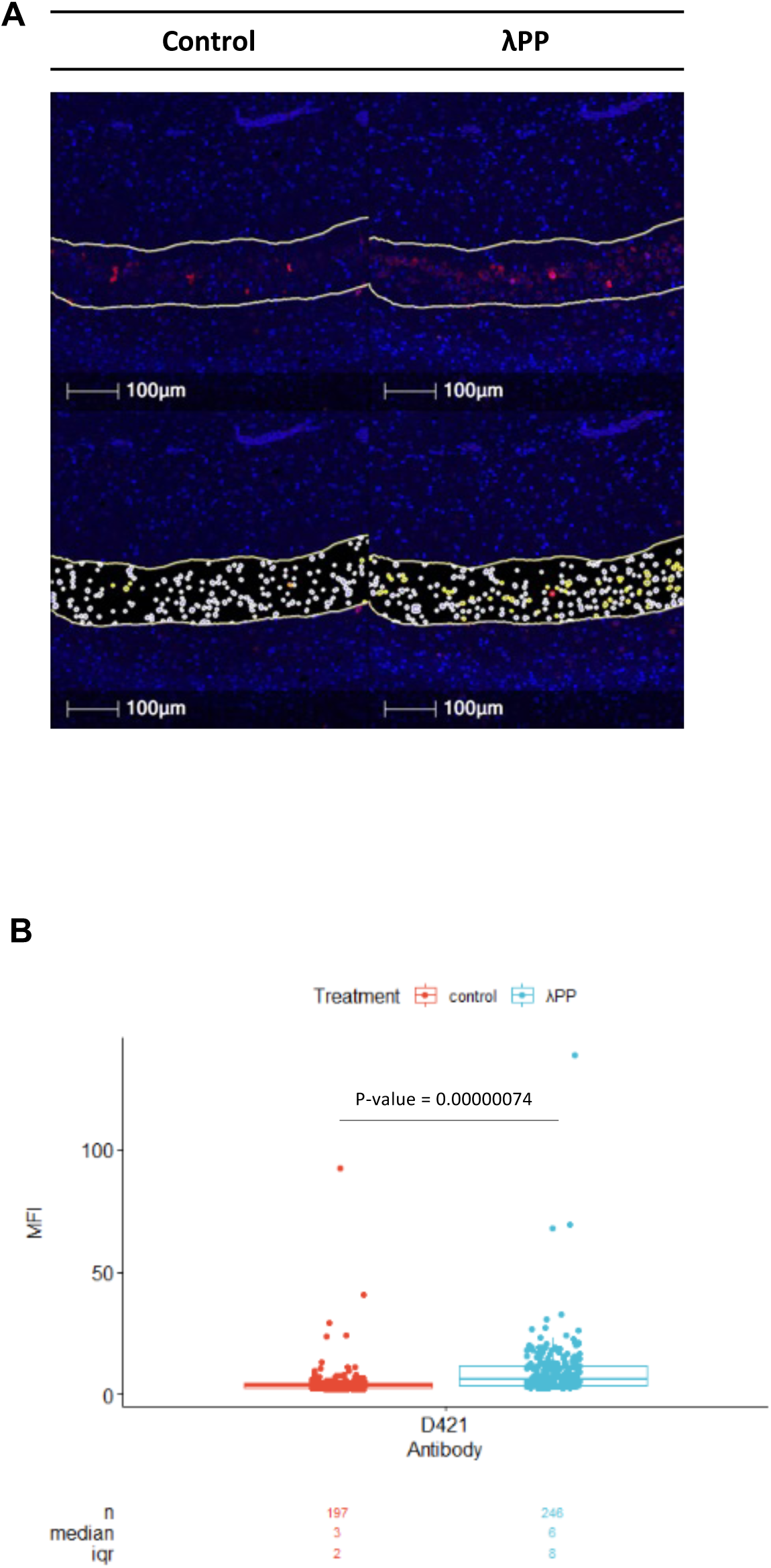
Phosphorylation partially inhibits binding of the D421 antibody clone in FFPE-IHC applications. Micrographs (**A**) and quantifications (**B**) of the signal mean fluorescence intensity (MFI) obtained in rTg4510 mouse brain cortex stained with the D421 antibody, before (**A** - left; **B** - red) and after (**A** - right; **B** - blue) λPP treatment. Tau signal is shown in red (**A**). Nuclei are stained with DAPI (blue) (**A**). The area used for quantifications is outlined (**A**). In A, bottom rows show all the cells identified by the image analysis pipeline. Tau-negative cells are coloured in white (**A** - bottom row). Tau-positive cells are coloured in yellow and orange, indicating moderate and high Tau staining signal intensity, respectively (**A** - bottom row). Scale bars = 100 µm (**A**). n = total number of cells of cells analysed, iqr = inter-quartile range (**B**). p-values were calculated using a two-tailed, independent *t test*.

